# Epigenome-informed Polymer Modeling Reveals Chromosome Folding Landscapes across Distinct Genetic and Epigenetic Contexts

**DOI:** 10.64898/2026.11.04.764273

**Authors:** Saman Bazmi, Alexander Turco, Sushant Kumar

## Abstract

Three-dimensional (3D) genome organization is critical for gene regulation, enabling long-range interactions between genes and distal regulatory elements. Genomic structural variations and epigenetic differences can perturb this organization, contributing to phenotypic variation and tissue-specific gene regulation. Prior studies have assessed the impact of these genetic and epigenetic changes on 3D genome folding, primarily using population-averaged features such as topologically associating domains and loops. However, growing evidence from single-cell and live-imaging studies indicates that chromosomes are highly dynamic and exist as heterogeneous ensembles of conformations. Here, we applied an epigenetic-state-informed polymer modeling framework to generate chromosomal conformational ensembles directly from cell- and tissue-specific epigenomic data. The resulting ensembles reproduced experimental Hi-C maps across cell lines and tissue types. Comparison of these conformational ensembles revealed that, although polymer dynamics were conserved across cell lines, their folding landscapes diverged. These differences arose primarily from cell-line-specific chromatin-state interaction networks and were concentrated in rare, long-lived contacts. Furthermore, modeling a TAD-boundary-disrupting deletion without re-optimizing the model parameters indicated a shift in the chromatin conformational ensemble from predominantly insulated to predominantly fused states. The modeled deletion also selectively strengthened active chromatin contacts and differentially disrupted enhancer contacts involving neighboring genes. Finally, comparison of two tissue types from the same donor revealed epigenetically driven rewiring of chromatin-state interaction networks. These findings establish chromosomal conformational ensembles as a sensitive readout of genome reorganization across distinct genetic and epigenetic contexts.

## BACKGROUND

Three-dimensional (3D) genome organization facilitates gene regulation by enabling interactions between genes and their regulatory elements^1,2^. Large-scale functional genomics initiatives, including the ENCODE^3^, 4D Nucleome^4^, and ENTEx^5^ projects, have produced comprehensive maps of chromatin accessibility, histone modifications, DNA methylation, transcription factor occupancy, and 3D genome architecture across diverse cell lines and tissue types. These datasets have enabled increasingly detailed reconstructions of gene regulatory networks by identifying cis-regulatory elements and linking them to their target genes^6^. Among these molecular features, 3D genome organization plays a particularly important role by mediating physical interactions between genes and distal regulatory elements, thereby coordinating transcriptional activity. Therefore, a mechanistic understanding of the principles governing chromosome folding is critical for explaining how gene regulation varies across cell types, individuals, and disease states.

The canonical view posits a hierarchical organization of 3D genome architecture within the cell nucleus across multiple levels, including DNA loops^1^, topologically associating domains (TADs)^7,8^, compartments^9^, and chromosome territories^10^. These structural components are considered essential for proper gene regulation^11^. For instance, chromatin loops facilitate interactions between gene promoters and distal regulatory elements, enabling appropriate gene expression^1,12^. Similarly, TADs insulate regions of high internal interaction, potentially restricting communication between specific enhancers and promoters^7,8,13^. Perturbations to these organizational features are often associated with altered gene regulation that can promote a wide range of developmental^14,15^ and disease phenotypes^16,17^, highlighting the functional importance of higher-order chromosome architecture.

Both genetic and epigenetic variation can influence chromosome organization, thereby reshaping gene regulatory networks^15,18^. For example, studies have shown that structural variants (SVs) in the genome (including deletions, duplications, inversions, and translocations) can alter the dosage and positioning of regulatory elements by disrupting TAD boundaries and creating new chromatin contacts^14,19^. These large-scale genomic rearrangements can lead to enhancer hijacking^20,21^, TAD fusion, loss of regulatory insulation^14,17^, and other forms of regulatory rewiring in various diseases, including cancer^17,21^. Similarly, epigenetic differences among cell and tissue types can alter chromatin states and interaction patterns, thereby influencing chromatin compartmentalization and large-scale chromosome folding^16,18,22^. Collectively, these findings indicate that both genetic and epigenetic variation affect gene regulation not only through local sequence- or chromatin-level changes but also by disrupting higher-order genome architecture.

Our current understanding of chromosomal organization stems largely from chromosome conformation capture technologies such as Hi-C and related assays^1,9,23^. These methods have provided unprecedented insights into genome architecture and identified many organizational features observed across distinct organisms^7,24^. Nonetheless, Hi-C contact maps represent population-averaged interaction frequencies across large numbers of cells and therefore provide only a limited view of the underlying structural heterogeneity^25^. In contrast, recent advances in single-cell genomics^25–27^ and imaging-based techniques^28,29^ reveal a dynamic and heterogeneous view of chromosome organization, in which chromosomes can sample a diverse range of conformational states over time^30^. These observations challenge the traditional view of chromosomes as static structures and instead support a model in which chromosomes exist as ensembles of conformations^28,29,31^.

While experimental techniques such as single-cell Hi-C^25,26^, multiplexed imaging^28^, and FISH-based assays^29^ can reveal chromosomal conformational heterogeneity, comprehensive characterization of chromatin conformational ensembles remains challenging due to high experimental costs, technical complexity, and sparse data^28,32^. Consequently, computational approaches offer a complementary strategy for inferring chromatin conformational landscapes from genomic and epigenomic data^33–35^. Existing computational methods can be broadly classified into data-driven and mechanistic modeling frameworks. Data-driven methods primarily reconstruct chromosome structures directly from experimentally measured contact maps^36–39^. In contrast, mechanistic modeling methods use polymer physics and biologically plausible intra-chromosomal interactions to simulate the chromosomal folding process^34,40–44^. This allows polymer modeling frameworks to generate biophysically grounded ensembles of chromosome conformations and provide mechanistic insights into genome organization^38,42,44–47^.

Recent studies have used polymer-based models to replicate experimentally observed chromatin architecture across organisms^48–50^, chromosomes^42,44,51^, and cell types^33,34,38^. These polymer models have also been used to investigate how SVs reconfigure TADs, create ectopic enhancer–promoter contacts, and switch loci between regulatory conformations^52^. However, these studies were largely locus-specific, relied on parameters inferred from wild-type contact data, and assessed SV impact using ensemble-averaged contact maps. More broadly, these studies have primarily focused on the effects of SVs^53–56^ and chromatin-state differences^1,22,57,58^ on canonical 3D-genome features such as chromatin loops, TADs, and compartments derived from ensemble-averaged Hi-C maps. Consequently, we still lack comprehensive insight into how SVs and tissue-type-specific chromatin states alter the distribution of accessible chromosomal conformations, contact persistence, compartment segregation, energetic heterogeneity, and other biophysical properties derived from chromosomal folding ensembles. Characterizing these ensemble-level properties is critical for gaining mechanistic insights into how genetic and epigenetic perturbations reshape chromatin interactions and ultimately rewire gene regulatory programs.

Therefore, we used an epigenetic-state-informed polymer modeling framework to generate chromosomal conformational ensembles directly from diverse cell lines and tissue-specific epigenomic data. This approach allowed us to systematically examine how chromatin-state composition, SVs, and tissue-specific epigenetic differences shape chromosomal folding landscapes. Overall, we observed that chromosomes with similar global polymer dynamics could adopt distinct conformational states, driven primarily by differences in chromatin-state interaction networks and compartmental organization. Additionally, polymer modeling of a TAD boundary-disrupting deletion revealed a redistribution of conformational-state populations, shifting chromosomes from insulated to fused folding states and reorganizing interactions among active chromatin regions. Furthermore, a systematic evaluation of tissue-specific epigenetic differences revealed significant changes in 3D contact persistence profiles, chromatin segregation, and conformational heterogeneity, despite modest changes in ensemble-averaged contact maps. Collectively, these findings underscore the critical role of chromatin-state interaction networks in shaping chromosomal folding landscapes and show that both genetic and epigenetic perturbations influence genome organization by modifying the distribution and stability of accessible chromosomal conformation states.

## RESULTS

### Overview of the polymer framework to model chromosomal folding

Similar to prior data-driven approaches^38^, we leveraged cell-type-specific epigenomic data with a physics-based polymer model to reconstruct three-dimensional chromosomal conformational ensembles. Briefly, we modeled each chromosome as a coarse-grained polymer chain, with non-overlapping genomic bins represented as beads connected by springs. We assigned each genomic bin (bead) an epigenetic state^59^ by combining histone modifications (H3K4me3, H3K27ac, H3K27me3, H3K9me3) and architectural protein (CTCF) profiles, discretizing the linear sequence of beads into a small set of epigenetic states (E1–E5) that define the block copolymer (**Fig. 1A**). We then performed polymer simulations to characterize the chromatin conformational ensemble using an energy function, U_chrom_ (r, *α*_i_) = U_CIC_ + U_GSC_ + U_ALS_, which combines Chromatin Identity Coupling (CIC), Genomic Separation Compaction (GSC), and Anchor-Mediated Loop Stabilization (ALS) terms (see Methods). We employed an iterative approach^42,60^ to update the state-interaction parameters *α*_i_ until the simulation-derived, ensemble-averaged contact maps matched the experimental Hi-C map for a given cell line or tissue type (**Fig. 1B**). Subsequently, we applied these optimized models to probe chromatin dynamics (e.g., compaction monitored by the radius of gyration R_g_ over time) across distinct cell lines and tissue types. Additionally, we assessed how structural variations reshape the chromatin folding landscape (comparing wild-type to deletion genotypes) and systematically characterized tissue-specific chromosomal conformational ensembles (**Fig. 1C**). Overall, this framework enabled us to investigate how genomic and epigenetic perturbations, in the form of SVs and tissue-specific epigenetic state differences, influence genome architecture and chromosomal folding ensembles.

**Figure 1.**
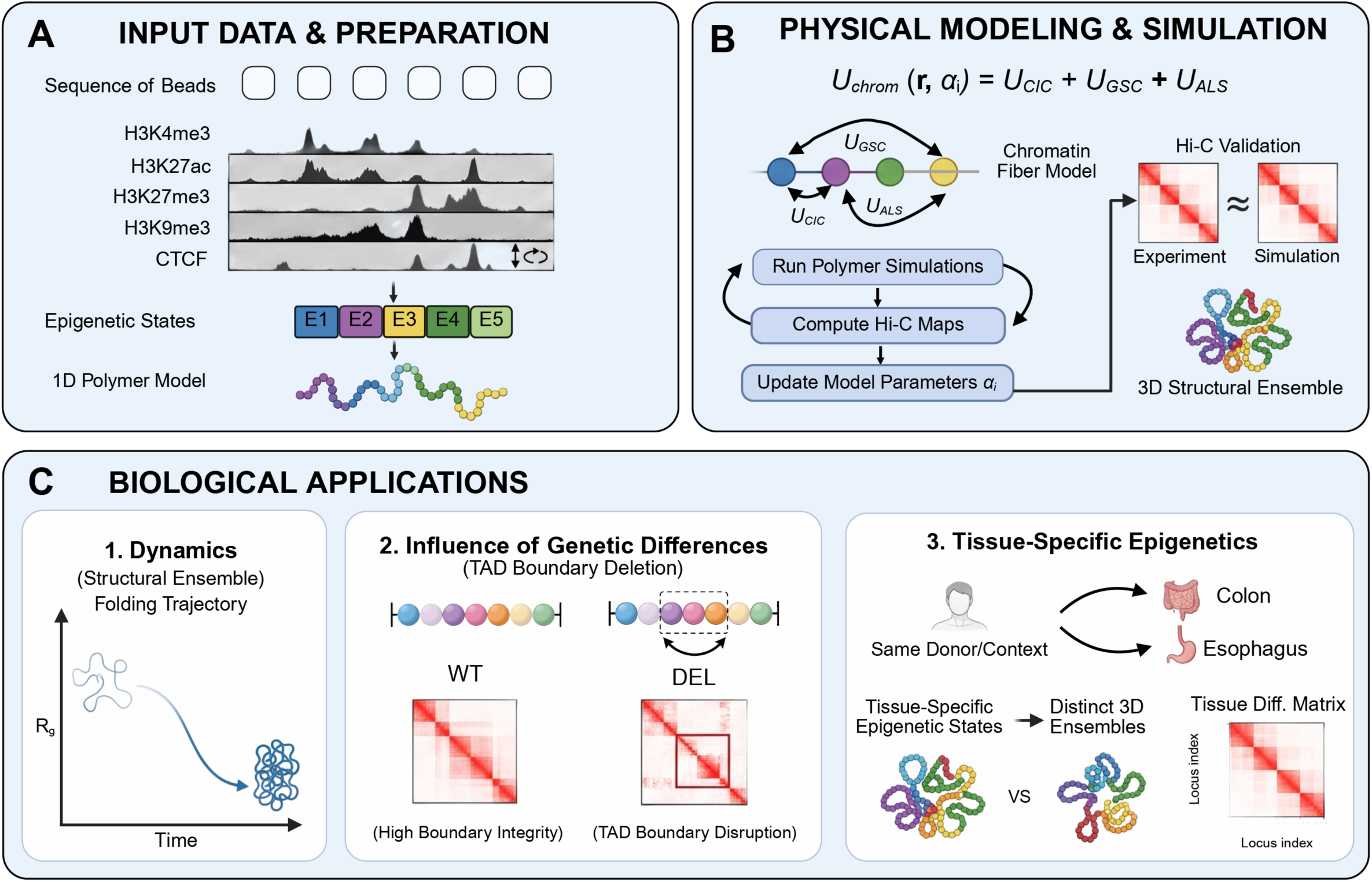
An epigenetic-state-informed polymer model reconstructs 3D chromosomal conformational ensembles from cell-type-specific epigenomic data. **(A)** Each chromosome is modeled as a coarse-grained polymer chain of non-overlapping genomic bins (beads) connected by springs. Histone-modification and architectural-protein tracks (H3K4me3, H3K27ac, H3K27me3, H3K9me3, and CTCF) are combined to assign each bead one of five epigenetic states (E1–E5), discretizing the linear sequence into a block copolymer. **(B)** Polymer simulations characterize the chromatin conformational ensemble using the energy function U_chrom(r, α_i) = U_CIC + U_GSC + U_ALS, comprising Chromatin Identity Coupling (CIC), Genomic Separation Compaction (GSC), and Anchor-Mediated Loop Stabilization (ALS) terms. State-interaction parameters α_i are updated iteratively until simulation-derived, ensemble-averaged contact maps reproduce the experimental Hi-C map for a given cell line or tissue type. **(C)** Optimized models are applied to investigate chromatin dynamics, including compaction (radius of gyration, R_g, over time) and 3D conformational collapse; to assess how structural variants (deletion vs. wild-type) reshape local contact frequencies and the chromatin folding landscape; and to characterize tissue-specific chromosomal conformational ensembles, revealing differential locus-locus contacts across cell lines and tissues.

### Model benchmarking and epigenetic state interactions during chromosomal folding

We first assessed how closely our polymer model-derived contact maps reproduced the 3D genome organization measured by experimental Hi-C across three cell lines^61^, including GM12878 (lymphoblastoid), H1-hESC (embryonic stem cell), and IMR90 (fetal fibroblast). After parameter optimization, we observed a strong correlation between simulation-derived and experimental Hi-C contact maps for each autosomal chromosome. For example, we compared the experimental map with the simulated map of chromosome 8 in the GM12878 cell line at 50-kb resolution (1 bead = 50 kb) and found that the model recovers the hallmark features of the experimental data. We observed strong concordance for the compartmental plaid pattern, long-range interaction blocks, and domain-level organization, yielding a high Pearson correlation (*ρ* = 0.97) between the simulated and experimental contact maps (**Fig. 2A**). We observed these agreements genome-wide, as the Pearson correlation between simulated and experimental contact maps remained uniformly high across all autosomes (chr1–chr22) and all three cell lines (**SI Fig. 1-Fig. 21**). This consistency between simulation- and experimental Hi-C-derived contact maps demonstrates that the model was able to accurately reconstruct compartment strength and preserve the genome-wide interaction-frequency distributions. Furthermore, the absolute error supported the same conclusion, as the root-mean-square error (RMSE) between experimental and simulated contact-probability curves remained low across all chromosomes and cell lines, ranging from approximately 0.004 to 0.017 (**SI Fig. 22A**). The RMSE for most chromosomes fell at the low end of this range (≈0.004–0.006), indicating stable performance with no systematic chromosome-specific bias. Only a few chromosomes showed modestly elevated error, including chromosome 4 across all three cell lines (≈0.016–0.017, the maximum of the scale), with localized increases at chromosome 19 in GM12878 and chromosome 8 in IMR90 (≈0.010–0.012). Our simulation-derived chromosomal conformational ensembles also accurately captured the polymer-scaling behavior of the genome. For instance, the simulated contact probability (P(s)) as a function of genomic distance tracked the experimental curve across the full range of genomic separations and followed the expected power-law decay for chromosome 8 in the GM12878 cell line (**Fig. 2B**). Fitting this decay yields a scaling exponent of *κ* = −0.965 for the experiment and *κ* = −1.007 ± 0.003 for the simulation, a difference of less than 5%. The two curves overlap from short range (s ≈ 1) through the mid-range decay and diverge only at the largest separations, where experimental noise dominates. Together, these high Pearson correlations, low RMSE values, and matched scaling exponents show that each cell-line-specific model was well optimized, capturing the correct polymer-compaction regime and the proper balance between short-range looping and long-range compartmental interactions.

**Figure 2.**
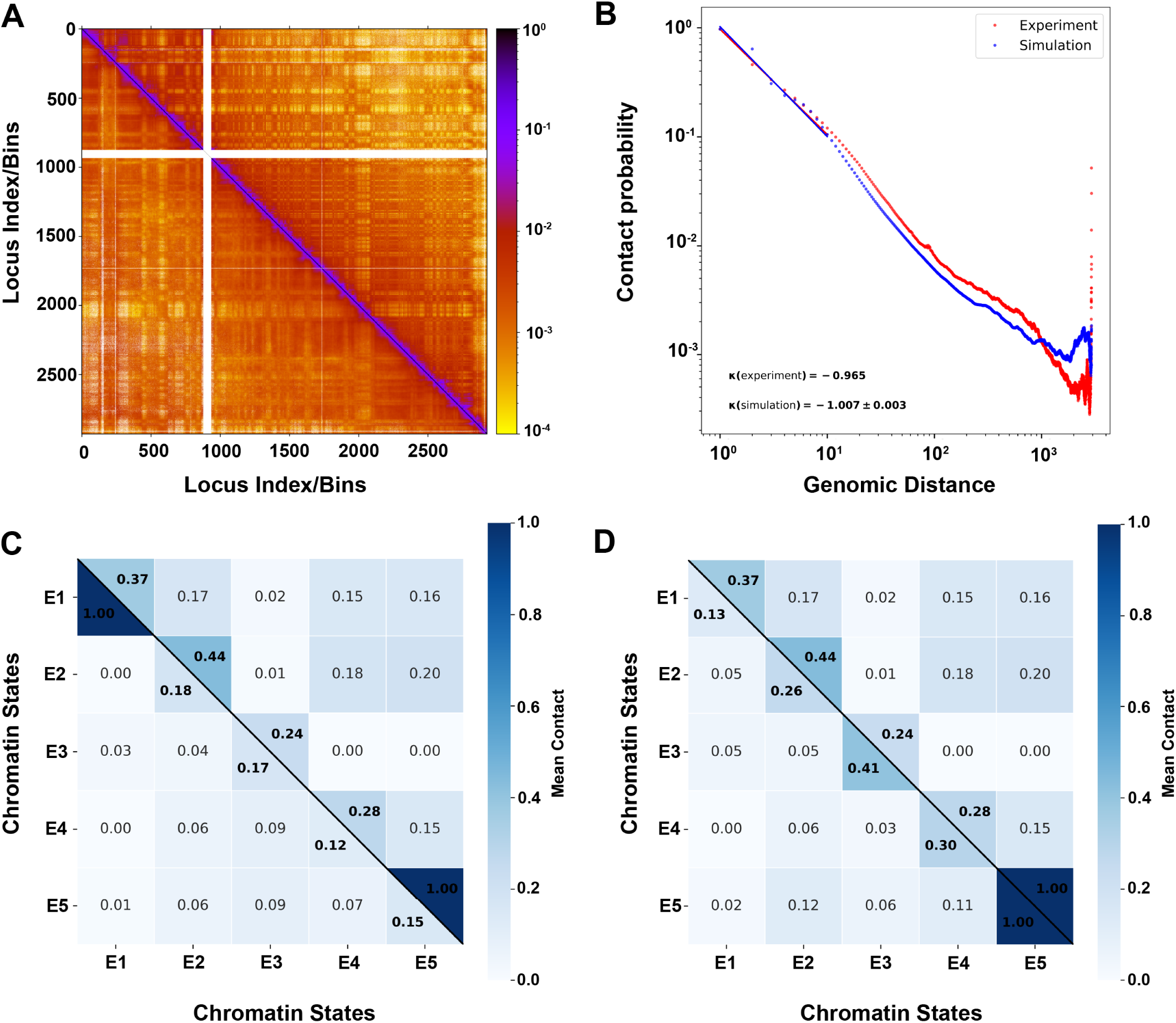
The optimized polymer model reproduces experimental Hi-C organization and reveals cell-type-specific chromatin-state interactions. **(A)** Experimental (upper triangle) and simulated (lower triangle) Hi-C contact maps for chromosome 8 in GM12878, showing compartmental plaid patterns, long-range interaction blocks, and domain-level organization (Pearson r = 0.97); maps for all remaining chromosomes across the three cell lines are in the Supporting Information. **(B)** Contact probability P(s) versus genomic distance for chromosome 8 GM12878 (red, experiment; blue, simulation), with fitted scaling exponents α(experiment) = −0.965 and α(simulation) = −1.007 ± 0.003. **(C)** State–state interaction heatmaps (normalized mean contact, 0–1) for epigenetic states E1–E5; diagonal = self-interactions, off-diagonal = cross-state mixing. Each panel overlays two cell lines (GM12878 (upper) vs. H1-hESC (lower triangle)). **(D)** State–state interaction heatmaps (normalized mean contact, 0–1) for epigenetic states E1–E5; diagonal = self-interactions, off-diagonal = cross-state mixing. Each panel overlays two cell lines (GM12878 (upper) vs. IMR90 (lower triangle)).

Next, we systematically compared global polymer dynamics and mobility across the three cell lines by quantifying the mean-squared-displacement exponent, *ς* (MSD ∼*τ*^ς^). We observed that MSD remained in the sub-diffusive regime (*ς* < 1) across all cell lines (**SI Fig. 23**). GM12878 showed a tight distribution, with its A and B compartments both centered near *ς* ≈ 0.49, whereas IMR90 and H1-hESC sampled a broader distribution (spanning *ς* ≈ 0.44–0.50) with means near *ς* ≈ 0.48. However, overall differences in global mobility were not statistically significant, suggesting that the cell lines share conserved polymer-backbone physics and comparable short-time dynamic constraints, and that their structural differences do not arise from global polymer mobility. Nonetheless, the slightly elevated distribution of GM12878 relative to the broader distributions of IMR90 and H1-hESC might indicate heterogeneity in interaction persistence. Given the conserved pattern in global polymer dynamics across the three cell lines, we reasoned that their structural differences instead arise from epigenetic organization. Therefore, we quantified both the abundance of each epigenetic state and the strength of interactions between these states in each cell line. We observed that cell lines differed substantially in genome-wide state composition (**SI Fig. 22C**). For instance, E3 (≈53%) in GM12878, E4 (≈51%) & E5 (∼19%) in H1-hESC, and E1 in IMR90 (≈42%) were the predominant chromatin states. Furthermore, interaction probabilities among beads corresponding to distinct chromatin states during chromosomal folding also differed across cell lines (**Fig. 2C and 2D**). A systematic comparison of normalized state-state interaction matrices across cell lines indicated that the off-diagonal contacts most clearly separated the folding regimes. For instance, chromatin states in the GM12878 cell line exhibit extensive mixing, leading to numerous cross-state contacts (E4–E5 ∼0.15 & E2–E5 ∼0.20). In contrast, IMR90 and H1-hESC chromosomal ensembles, on average, showed suppressed cross-chromatin-state contacts (**SI Fig. 24**). These chromatin-state interaction patterns among chromosomal ensembles across cell lines indicate two distinct chromatin regimes. For example, GM12878 adopted a diffuse interaction profile characterized by strong cross-state mixing and weak self-association, consistent with a heterogeneous, weakly phase-separated landscape. In contrast, H1-hESC and IMR90 conformational ensembles were enriched for same-state interactions along the diagonal, suggesting stronger compartmental segregation and more cooperative phase separation. Together, these results indicate that cell-type-specific differences in genome folding primarily arise from divergent epigenetic state composition and state-state interactions rather than from global polymer dynamics, implying that chromosomal folding heterogeneity is epigenetically encoded.

### Mapping persistent contacts to delineate conformational heterogeneity among cell lines

After characterizing global chromosomal dynamics across cell lines, we next examined how their conformational ensembles differ structurally. First, we quantified contact persistence separately for beads assigned to euchromatin and heterochromatin states. We extracted contact dwell lengths (see **Methods**) from the simulated ensembles and computed the upper quantiles of the dwell-length distribution (Q90) (**Fig. 3A)** and the survival fractions of long-lived contacts (S≥10) (**Fig. 3B)** for each compartment independently across chromosomes. We observed compartment-specific persistence profiles across three cell lines. For instance, in GM12878, euchromatin contacts persist slightly longer than heterochromatin contacts (**Fig. 3A-B)**. In contrast, in IMR90, this ordering reverses, with heterochromatin contact persistence exceeding that of euchromatin. Similarly, in the H1 cell line, contacts involving heterochromatin beads strongly dominate those involving euchromatin (**Fig. 3A-B)**. Moreover, compartmental-level contact asymmetry increases monotonically along the distribution tail, from Q90 to Q99 (**Fig. 3A & SI Fig. 25A-B**) and from S≥5 to S≥20 (**Fig. 3B & SI Fig. 25C-D**), indicating that the most persistent contacts exhibit the greatest compartment-specific separation. These observations suggest that persistent contacts are distributed non-uniformly across chromatin and that compartment-specific differences in persistent contacts primarily drive chromatin contact heterogeneity across cell lines, with progressive stabilization of the heterochromatic compartment in IMR90 and, most prominently, in H1-hESC. Furthermore, these quantile and survival metrics indicated a heavy-tailed dwell-time distribution, with some contacts persisting beyond the average interaction lifetime. The dwell-time distributions of the H1 and IMR90 cell lines showed pronounced tail amplification, indicating selective enrichment of rare, long-lived chromatin interactions that promote enhanced compartmental stabilization and reduced conformational rearrangement. In contrast, the GM12878 cell line showed weaker tails, suggesting fewer persistent contacts and facilitating a more dynamic chromatin state with frequent reorganization.

**Figure 3.**
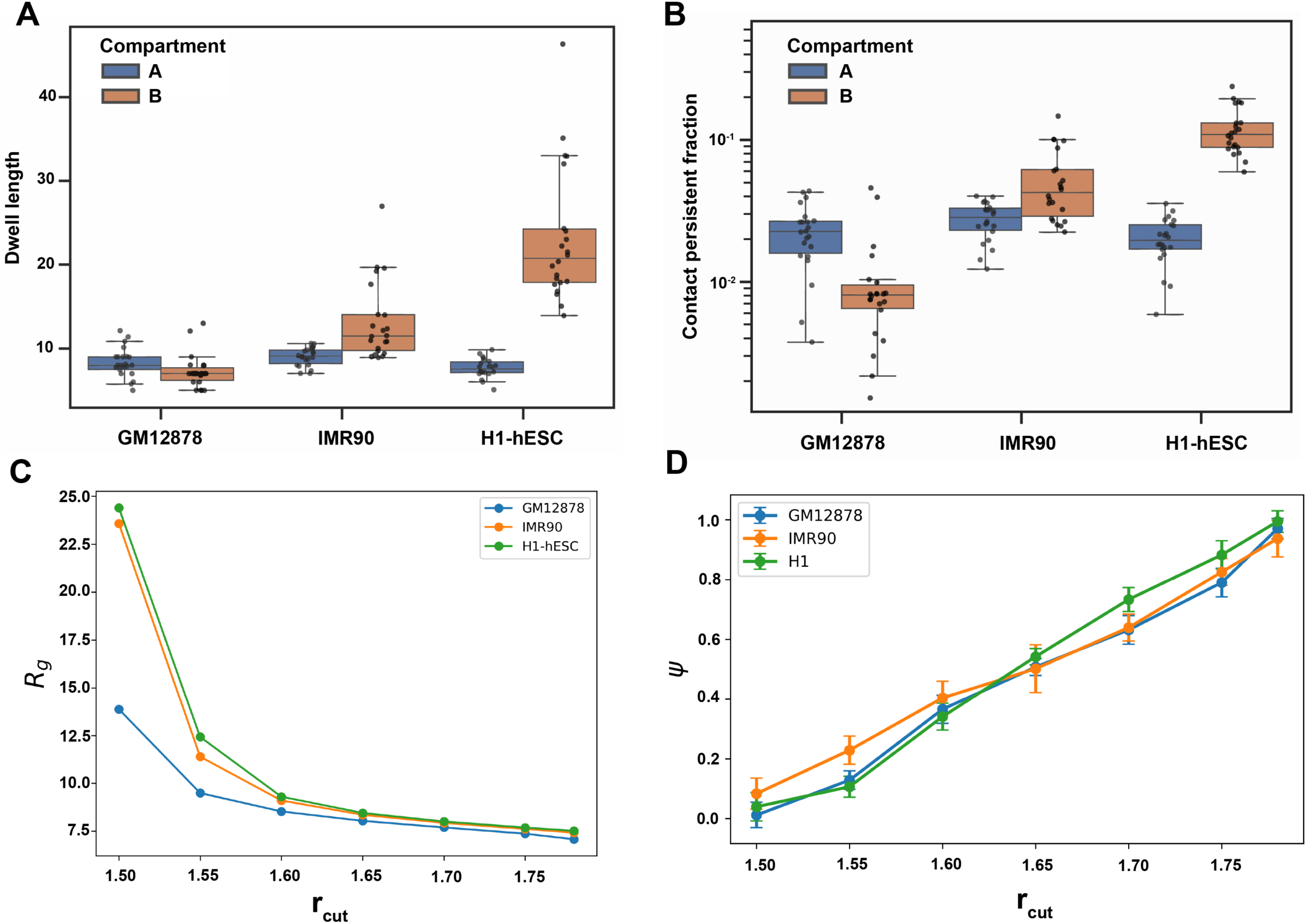
Cell-line heterogeneity is associated with compartment-specific contact persistence and modulated by interaction range. **(A)** Compartment-resolved contact dwell length metrics (Q90) for GM12878, IMR90, and H1-hESC, shown separately for A (euchromatin) and B (heterochromatin) compartments. **(B)** Long-lived-contact survival fractions (S≥10, log scale) for the same cell lines and compartments. Euchromatin persistence slightly exceeds heterochromatin in GM12878; heterochromatin exceeds euchromatin in IMR90; heterochromatin strongly exceeds euchromatin in H1-hESC, with the asymmetry growing from Q90→Q99 and S≥5→S≥20 (See SI). **(C)** Ensemble-averaged radius of gyration R_g_ versus interaction cutoff r_cut_ at μ = 3.15. R_g_ falls steeply between r_cut_ = 1.50 (R_g_ ≈ 13.8 for GM12878; ≈ 23.5 for IMR90; ≈ 24.5 for H1-hESC) and r_cut_ ≈ 1.60, then plateaus at R_g_ ≈ 7.2–7.5 by r_c_ ≈ 1.75–1.78. **(D)** Average normalized A/B compartment strength versus r_c_ for GM12878, IMR90, and H1-hESC; strength rises approximately linearly from ≈ 0.0–0.1 (r_cut_ = 1.50) to ≈ 0.93–1.00 (r_cut_ ≈ 1.78), with H1-hESC reaching the strongest segregation. Error bars denote ± statistical error (≈ ±0.03–0.05).

Next, we investigated whether these compartment-level differences in cell lines extend to the genome scale by reconstructing each persistence metric across the genome using compartment-specific statistics as weighted averages, with weights proportional to the number of contributing contacts (**SI. Fig. 26A-F).** Across all metrics (Q90 to Q99, and S≥5 to S≥20), the reconstructed means follow a consistent ordering (GM12878 < IMR90 < H1-hESC), with the largest separation observed for the extreme-tail metric Q99 (**SI. Fig. 26C)**. This suggests that enrichment of persistent contacts at the compartment level propagates to the genome scale, where H1-hESC shows the strongest genome-wide stabilization of chromatin contacts and GM12878 shows the greatest fluctuations. The reconstructed distributions further reinforce this observation, with H1-hESC and IMR90 maintaining higher and more stable fractions of long-lived contacts, as reflected in higher medians and narrower spreads across chromosomes, indicating a more stable structural regime (**SI. Fig. 26D-F)**. In contrast, GM12878 displays a broader distribution, a lower median, and greater chromosome-to-chromosome variability. These results indicate that GM12878 chromatin fluctuates more dynamically, whereas H1 and IMR90 show more stable contact patterns. This distinction arises from differences in contact persistence rather than overall contact abundance.

To link each cell line’s contact persistence profile to the underlying polymer physics, we next examined how the chromatin conformational ensemble responds to the model’s interaction parameters. We found that varying the interaction cutoff (r_cut_) at a fixed mean interaction strength produced a sharp collapse in the radius of gyration (R_g_), shifting the system from extended to folded conformations and identifying r_cut_ as the primary control of global compaction (**Fig. 3C & SI Fig. 27**). This compaction tracked a saturating energy landscape, in which compact conformations occupied deeper energetic basins and extended ones occupied a flatter, less sensitive regime (**SI Fig. 28**), consistent with stable, long-lived structures residing in deeper basins. These relationships were robust across the mean interaction strength (μ=3.00–3.25), with r_cut_ remaining the dominant control over compaction, while μ acted only as a weak secondary modulator (**SI Fig. 28**). Next, we measured the normalized euchromatin/heterochromatin compartment strength (see **Methods**) as a function of r_cut_ to investigate its influence on chromatin compartmentalization (**Fig. 3D**). In all three cell lines, compartment strength rose steadily and nearly linearly with r_cut_, from ≈0 at r_cut_ = 1.50 to ≈0.93–1.00 at r_cut_ ≈ 1.78 (mean ± SEM across chromosome 10). The curves tracked closely across the mid-range and separated at high r_cut_, with H1-hESC showing the strongest segregation (approaching 1.00). Increasing the interaction range therefore enhanced same-state attraction and stabilized compartmental organization, establishing compartmentalization as a tunable, phase-segregation-like property governed by interaction parameters. Together, these analyses indicate that chromatin-folding heterogeneity manifests primarily through rare, persistent contacts, and that interaction range governs compaction, energetics, and compartment strength, linking long-lived contacts to deeper energetic basins and stronger compartmental segregation.

### Modeling the Effects of Structural Variation on Chromatin Dynamics

We next examined how SVs reshape chromatin organization and chromosomal folding ensembles. To enable detailed sampling of folding ensembles, we focused on a well-characterized locus where an SV has been experimentally shown to disrupt TAD architecture. A prior study^53^ showed that the HGSVC^62^ sample HG00864 carries a deletion at chr8:143,618,315–143,632,522 that disrupts TAD architecture in this region, whereas no SVs were identified in this interval in GM19036, which served as the wild-type (WT) reference. We modeled the surrounding 5 Mb region (chr8:140,000,000–145,000,000) at 5 kb resolution, yielding a polymer of 1,000 beads, for both lymphoblastoid cell lines. To represent the deletion in HG00864, we removed the four consecutive beads spanning the deleted interval and reconnected the flanking segments to preserve chain continuity. Additionally, we used GM12878 ChromHMM annotations as a proxy for HG00864, given their shared lymphoblastoid origin (see Methods). For each system, we obtained 1.5 × 10⁷ conformations from polymer simulations and used the resulting ensembles to construct in silico Hi-C maps and compare the structural and energetic features of the WT and SV-affected chromosomes.

The optimized polymer model accurately recapitulated Hi-C contact patterns in both systems, with simulated WT and SV maps each showing a high Pearson correlation coefficient (*ρ*=0.97) relative to their experimental counterparts (**SI Fig. 29A-B**). This correlation holds without retuning the potential energy function beyond the deletion itself, confirming that the polymer directly incorporates the sequence rearrangement. Comparison of the ensemble-averaged contact maps from these simulations reveals local chromosomal reorganization (**Fig. 4A**), in which loci previously separated by the deleted region exhibited increased contacts across the deletion junction in the SV-affected model. Furthermore, the contact probability decay, P(s), separates the two ensembles (**SI Fig. 29C**), with the SV-affected ensemble maintaining higher contact probability than the WT across all genomic distances. Because the two systems derive from different individuals, part of this offset likely reflects intrinsic differences between GM19036 and HG00864 rather than the rearrangement alone. Therefore, we consider the separation in the P(s) profile suggestive of the ensembles occupying distinct contact regimes, rather than evidence that the deletion propagates to long range.

**Figure 4.**
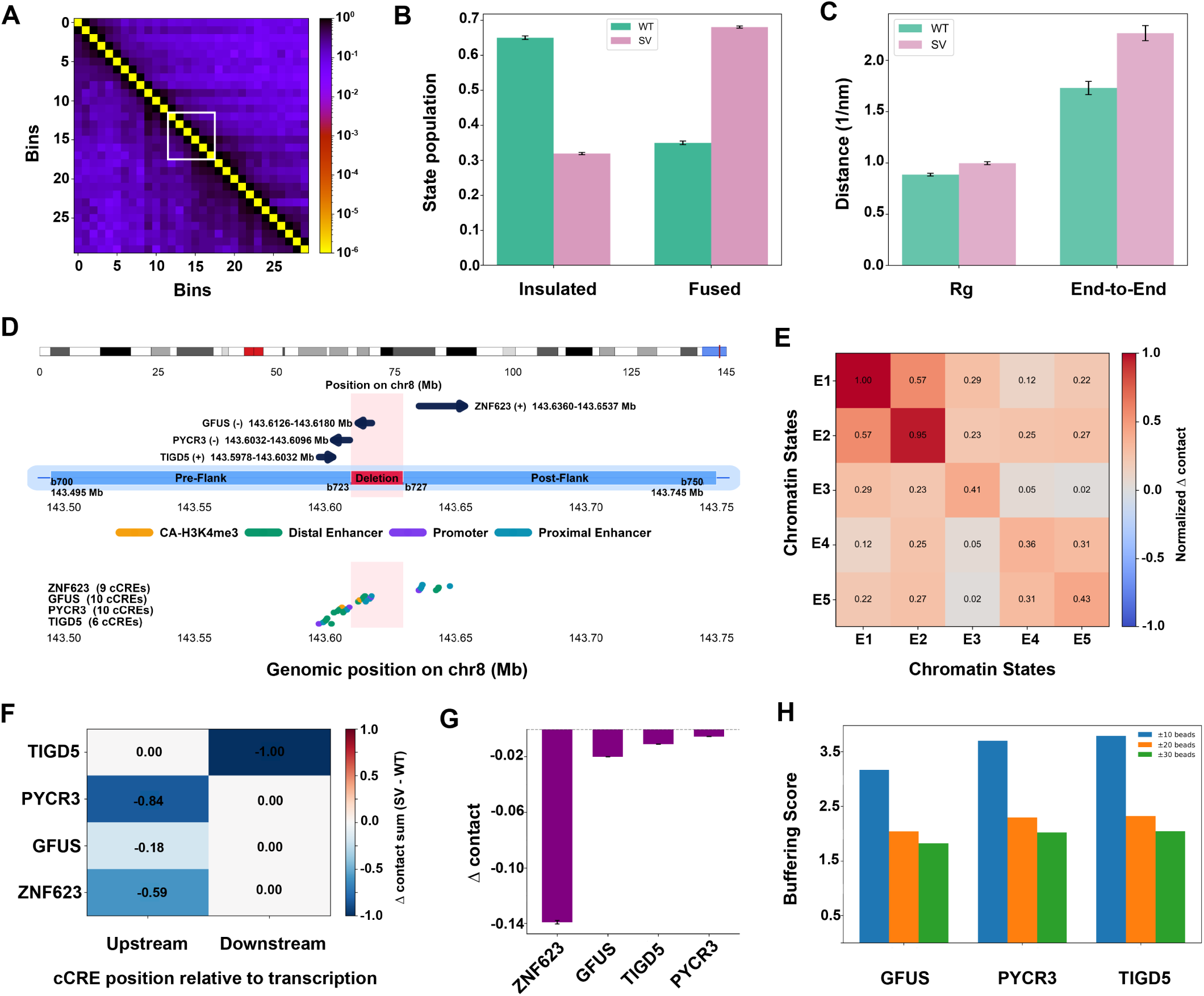
Structural variation remodels local chromatin folding states and gene-regulatory interactions at a chromosome 8 deletion breakpoint. **(A)** Coarse-grained (30-bin) contact matrix of the breakpoint-spanning locus, with the white box highlighting the bins that flank the deletion breakpoint. **(B)** Fraction of each ensemble classified as insulated or fused in the wild-type (WT) and SV-affected ensembles. The WT ensemble is predominantly insulated (0.65), whereas the SV ensemble is predominantly fused (0.68). **(C)** Radius of gyration (Rg) and end-to-end distance of the locus for each ensemble (nanometers (nm)). The SV-affected ensemble adopts a more extended end-to-end configuration. **(D)** Genomic context of the modeled region. Top, chromosome 8 (hg38) with the simulated 143.50–143.75 Mb segment highlighted in red. Middle, zoomed locus showing the bead model, the chr8:143,618,315–143,632,522 deletion (DEL; four beads, 723-726, red shading) separating the left flank (beads 700-722, blue shading) from the right flank (beads 727-750, blue shading), and genes TIGD5 (+), PYCR3 (−), GFUS (−), and ZNF623 (+). Bottom, gene-linked candidate cis-regulatory element (cCRE) intervals used in the model, colored by class (CA-H3K4me3, TF accessible, distal enhancer, promoter, proximal enhancer); the number of cCREs assigned to each gene is indicated. **(E)** Normalized change in contact frequency between chromatin states E1–E5, scaled to the most strongly affected pair (E1 − E1 = 1.00). The SV-affected ensemble shows the largest contact gains among the active states E1 and E2 (E1–E1 = 1.00, E2 − E2 = 0.95, E1 − E2 = 0.57). **(F)** Direction-aware contact change (Δ = SV − WT) for each gene, decomposed into upstream and downstream components relative to the direction of transcription; color encodes the signed Δ contact sum. **(G)** Gene-level enhancer contact rewiring, plotted as the change in enhancer contact between the SV and WT ensembles (Δ = SV − WT) for each gene. **(H)** Local regulatory buffering score for each breakpoint-associated gene (GFUS, PYCR3, TIGD5, ZNF623), computed as the distance-weighted sum of active cCREs within ±10, ±20, and ±30 beads (≈50, 100, and 150 kb), B = Σ aᵢ/(1 + dᵢ), where dᵢ is the bead separation between the gene and cCRE i, and normalized to ZNF623; higher values denote denser local active neighborhoods. Bars and heatmap values represent the mean across independent simulation replicas, and error bars in (B), (C), and (G) denote the statistical errors.

We classified chromosomal configurations from our simulations of WT and SV-affected polymer models as insulated (anchor–anchor distance < 2.5 σ) or fused (≥ 2.5 σ). In the insulated state, the domain flanking the deletion breakpoint remains segregated from neighboring domains, whereas in the fused state, these domains collapse into a single, contact-rich neighborhood. We calculated state populations and their mean lifetimes from simulation trajectories and observed that the insulated state predominates (65 ± 1%) and is long-lived, consistent with stable domain insulation, whereas the fused state is short-lived and represents only 35 ± 1% of structures in the WT simulation (**Fig. 4B**). In contrast, in the SV-affected simulation, the fused state prevails (68 ± 1%) with an extended lifetime (**see Fig. S30A**), whereas the insulated state is reduced to 32 ± 1% and becomes transient. Global conformational metrics further corroborated this shift, with the SV-affected polymer sampling more extended configurations. For instance, the mean radius of gyration (R_g_) and end-to-end distance(d_E-E_) of configurations from the SV-affected polymer simulation were higher (R_g_^SV^ = 1.0 nm & d^SV^_E-E_= 2.3 nm) compared to the WT model (R_g_^WT^ = 0.88 nm & d^WT^ = 1.7nm) (**Fig. 4C**). Accordingly, projecting the ensemble onto a switching coordinate defined by the distance between loci flanking the breakpoint (**Fig. SI 30B**) reveals a clear redistribution of structural probability. The WT ensemble peaks near ∼1.3 within a narrow basin, whereas the SV-affected ensemble centers near ∼2.5 with a broader profile and only partial overlap. This shift shows that the SV relocates the dominant structural basin rather than simply amplifying fluctuations around a shared minimum, thereby potentially reshaping the energetic accessibility of chromosomal conformations.

To link these structural differences to underlying chromatin states, we computed the normalized differential contact matrix (SV-WT), stratified by the five ChromHMM states^59^ (E1-E5) and excluding centromeres (**Fig. 4E**). We observed strong gains in homotypic contacts among the most active states, particularly E1 (1.00) and E2 (0.95), whereas E3-E5 showed more modest increases (0.36-0.43). These patterns indicate that the SV-affected ensemble likely comprises configurations that favor enhanced interactions among transcriptionally permissive chromatin states. Additionally, we found the off-diagonal terms in the differential contact matrix to be positive but weaker, with the E1-E2 interaction leading (0.57), while intermediate-state pair differences ranged from 0.02 to 0.31. Notably, positive mixed interactions between active (E1, E2) and intermediate (E3, E4) states suggest that the rearrangement brings previously segregated chromatin neighborhoods into closer spatial proximity. Together, these observations suggest that the presence of an SV likely reshapes chromatin organization by selectively enhancing interactions between specific epigenetic states, rather than uniformly increasing contacts, thereby creating a distinct epigenetic interaction signature compared to the WT.

### Assessing the Impact of Structural Variants on Gene Regulatory Interactions

We further investigated how the selected deletion (chr8:143,618,315–143,632,522) alters regulatory interactions among four genes flanking the breakpoint and their annotated candidate cis-regulatory elements (cCREs)^63^, in the polymer model. This deletion removes 4 beads (beads 723-726), separating the pre-flank (beads 700-722) from the post-flank (beads 727-750) region. Of the four affected genes, TIGD5, PYCR3, and GFUS lie immediately upstream of the deletion, whereas ZNF623 lies immediately downstream (**Fig. 4D**). We classified each gene-linked cCRE as promoter-like, proximal enhancer-like, distal enhancer-like, or TF-bound/accessible. We then calculated and compared mean interaction frequencies between linked cCREs and their corresponding genes across SV-affected and WT chromatin conformational ensembles. Because the four genes differ in strand orientation and position relative to the deletion, we quantified normalized contact changes (the ratio of the original to the maximum contact difference) and separated them into upstream and downstream components (**Fig. 4F**). This direction-aware comparison showed that contact loss was concentrated on the deletion-facing side of each gene. For PYCR3 (−0.84), ZNF623 (−0.59), and GFUS (−0.18), this corresponded to the upstream side, whereas for TIGD5, it corresponded to the downstream side. The opposite side of each gene remained largely unchanged. Therefore, contact loss was confined to the side of each gene facing the breakpoint, consistent with the removal of sequences it previously contacted. Comparison of gene-enhancer contacts between the WT and SV-affected ensembles revealed reduced enhancer contact frequencies for all four genes, although the magnitude of loss varied substantially (**Fig. 4G**). For example, ZNF623 showed the largest reduction in these interactions (Δ = −0.140 ± 0.001), approximately 7-fold greater than that of GFUS (Δ = −0.0201 ± 0.0001), 13-fold greater than that of TIGD5 (Δ = −0.0105 ± 0.0001), and 27-fold greater than that of PYCR3 (Δ = −0.0051 ± 0.0001). This suggests that ZNF623 experiences the most pronounced disruption of enhancer contacts following the deletion, potentially reflecting the altered regulatory neighborhood created by the SV. Closer inspection of contacts between genes and active cCREs showed that all four genes experienced the greatest losses in their immediate genomic neighborhoods (**Supplementary Figure S30D**). Contacts between regulatory elements increased, notably between pre-flank enhancers and post-flank promoters (Δ = 1.15 ± 0.01) and between pre-flank promoters and post-flank enhancers (1.10 ± 0.01). In contrast, gene-body contacts decreased, including gene–gene (0.58 ± 0.01) and gene–regulatory element interactions (0.65–0.95). These findings indicate that the deletion strengthens regulatory-element contacts across the junction while weakening gene-body interactions during chromosomal folding.

The pronounced differences in enhancer-contact loss observed for ZNF623 compared with other affected neighboring genes within the same deletion may indicate a dense local network of active regulatory elements that buffers genes against deletion-induced contact loss. Therefore, we quantified the active regulatory neighborhood for each gene and examined whether regulatory-element density was associated with the magnitude of contact loss. For each gene, we counted active cCREs within ±10, ±20, and ±30 beads (≈50, 100, and 150 kb) and weighted them by inverse genomic distance to calculate a buffering score, such that proximal regulatory elements contribute more strongly than distal elements. Briefly, B = Σ aᵢ / (1 + dᵢ), where aᵢ is the number of active cCREs in bead i and dᵢ is the distance (in beads) between bead i and the gene. We then normalized each score relative to ZNF623 (reference = 1.00). Within ±10 beads, GFUS, PYCR3, and TIGD5 occupied markedly denser active neighborhoods than ZNF623 (3.2-, 3.7-, and 3.8-fold, respectively; Fig. **4H**). This advantage narrowed monotonically as we widened the window, to 2.0–2.3-fold at ±20 beads and 1.8–2.0-fold at ±30 beads. This convergence shows that ZNF623 distributes its active elements distally, whereas GFUS, PYCR3, and TIGD5 concentrate them around their gene bodies. Therefore, local regulatory density, rather than total regulatory content, distinguishes ZNF623 from its neighbors.

If local density buffers chromatin contacts, genes with the densest neighborhoods should retain the most contacts after the deletion. Therefore, we computed each gene’s retention score as the ratio of SV-affected to WT contact frequency, R = C_SV_ / C_WT_, where R = 1.00 denotes full preservation. We found that GFUS, PYCR3, and TIGD5 retained 0.81–0.86 of their WT contacts across all windows, whereas ZNF623 retained only 0.54 locally, losing nearly half of its proximal contacts (**SI Fig. 30C**). This ranking matched the buffering scores (TIGD5 ≈ PYCR3 ≈ GFUS ≫ ZNF623), consistent with the buffering hypothesis. Additionally, ZNF623’s contact retention increased as we widened the window (0.54 → 0.70 → 0.82), indicating that the deletion strips its local regulatory layer while distal elements maintain residual connectivity and prevent the genes from becoming fully isolated (**SI Fig. 30C**). This observation suggests that dense local neighborhoods preserve enhancer contacts, whereas dependence on distal interactions leaves a gene potentially more vulnerable to the impact of genomic rearrangement.

To express these differences as absolute contact loss rather than ratios, we computed ΔC = C_SV_ − C_WT_ for each gene and window. Every gene showed a net loss of contacts, and none showed a net gain, confirming that the deletion is broadly disruptive (**Supplementary Fig. 30D**). The magnitude of contact loss followed a similar pattern, with GFUS, PYCR3, and TIGD5 exhibiting relatively modest reductions (0.09–0.10 locally, decreasing to 0.03 at wider windows) compared with ZNF623 (ΔC = −0.140 ± 0.001; **Fig. 4G**), which eased to −0.02 distally (**Supplementary Fig. 30D**). The consistent ordering across buffering score, retention, and absolute loss suggests a mechanism in which a dense local neighborhood absorbs the impact of the deletion. In this buffering mechanism, when the deletion removes one active contact, a redundant nearby element can mitigate its impact. These observations suggest that GFUS, PYCR3, and TIGD5 carry that redundancy, whereas ZNF623, embedded in a sparser local neighborhood, lacks it and rewires accordingly, accounting for its outsized loss (**Fig. 4G).**

Because both WT (GM19036) and SV-affected (HG00864) models share identical chromatin-state annotations and parameters, differences between the ensembles are attributable solely to the deletion. However, while this design isolates the effect of the SV, it does not capture other sources of inter-individual variation, such as additional variants or sample-specific chromatin states, that may further shape chromatin architecture in vivo. Within this framework, the consistent ordering of the four genes across buffering score, retention, and absolute contact loss identifies local regulatory density as a major determinant of susceptibility to deletion-induced contact loss. Together, these analyses support a model of local regulatory buffering in which dense active chromatin neighborhoods confer relative resilience to SV-induced disruption, whereas genes that rely on distal regulatory elements, such as ZNF623, are more likely to exhibit pronounced contact loss and reorganization.

### Tissue-Specific Analysis of Chromosomal Dynamics

Prior studies^14,19^ have implicated 3D genome architecture in tissue-specific regulatory divergence, yet population-averaged Hi-C maps offer limited mechanistic insight into how chromosomal folding ensembles differ across tissues. To directly link tissue identity to changes in chromosomal organization, we applied our polymer modeling framework to chromosome 10 in sigmoid colon (SC) and esophageal muscularis mucosa (EMM) tissues from the same ENTEx donor^5^. We focused on a single chromosome to enable extensive conformational sampling in both tissues. We selected chromosome 10 for these simulations because it is a mid-sized chromosome with both gene-dense and gene-poor regions, making it representative of genome-wide chromatin organization. This paired-tissue design controls for genetic background, isolating epigenetically driven differences in folding and their regulatory consequences. Overall, our simulation-based ensemble-averaged contact maps were highly concordant across tissues (Pearson r = 0.96 for each; **Supplementary Fig. 31A-B).** The simulated contact maps also reproduced compartment-level checkerboard patterns, TAD-scale block structures, and focal contact enrichments, including within three regions of interest (RoI1, 34–50 Mb; RoI2, 44–47 Mb; RoI3, 78–86 Mb), indicating that the model accurately captures tissue-specific chromatin architecture.

Next, we computed contact map differences (SC – EMM) between two tissues directly from Hi-C data (**Fig. 5A**) and found that tissue-specific differences are concentrated within spatially defined regions rather than distributed globally. Moreover, these contact differences clustered into discrete sub-blocks of the matrix that aligned with regions of interest (RoI) boundaries (**Fig. SI 31 A-B**), indicating localized rewiring of the 3D contact network. We performed a sliding-window (±1 Mb) analysis of mean relative absolute contact differences, which confirmed this heterogeneity and revealed sharp peaks separated by extended regions of minimal divergence (**Fig. SI 31E**). These peaks coincided precisely with the predefined RoIs, motivating targeted analysis of their conformational ensembles. In particular, we divided chromosome 10 into bins, annotated them with four epigenetic states (E1–E4), and quantified epigenetic-state compositions within each RoI (**Fig. 5B**) to identify the epigenetic origins of these structural differences. We observed that tissue identity reshaped epigenetic-state composition in a region-specific manner. For instance, EMM tissue was enriched for active chromatin states, whereas SC shifted the same loci toward repressive states (particularly E3 and E4) in RoI1 and RoI2. In RoI3, genomic bins in SC and EMM tissues were distributed between active and repressive states in markedly different proportions, despite sharing the same underlying DNA sequence. We next asked how these compositional shifts altered chromatin state-to-state contact preferences. We calculated chromatin state-interaction differences from simulations (SC – EMM; **Fig. 5C**), which revealed selective strengthening of homotypic E1–E1 interactions in SC compared to EMM chromatin conformational ensembles. In contrast, we observed weakening of E2–E2 interactions and several heterotypic interactions, including E1–E3 and E2–E3, in SC compared to EMM conformations. These observations suggest that tissue-specific architecture emerges from targeted modulation of compartment–compartment interactions rather than uniform compaction or expansion of the contact network. Further analysis of chromatin-state transitions between tissues revealed heterogeneous changes in contact frequency across distinct chromatin state-switching categories (**Fig. SI 31F**).

**Figure 5.**
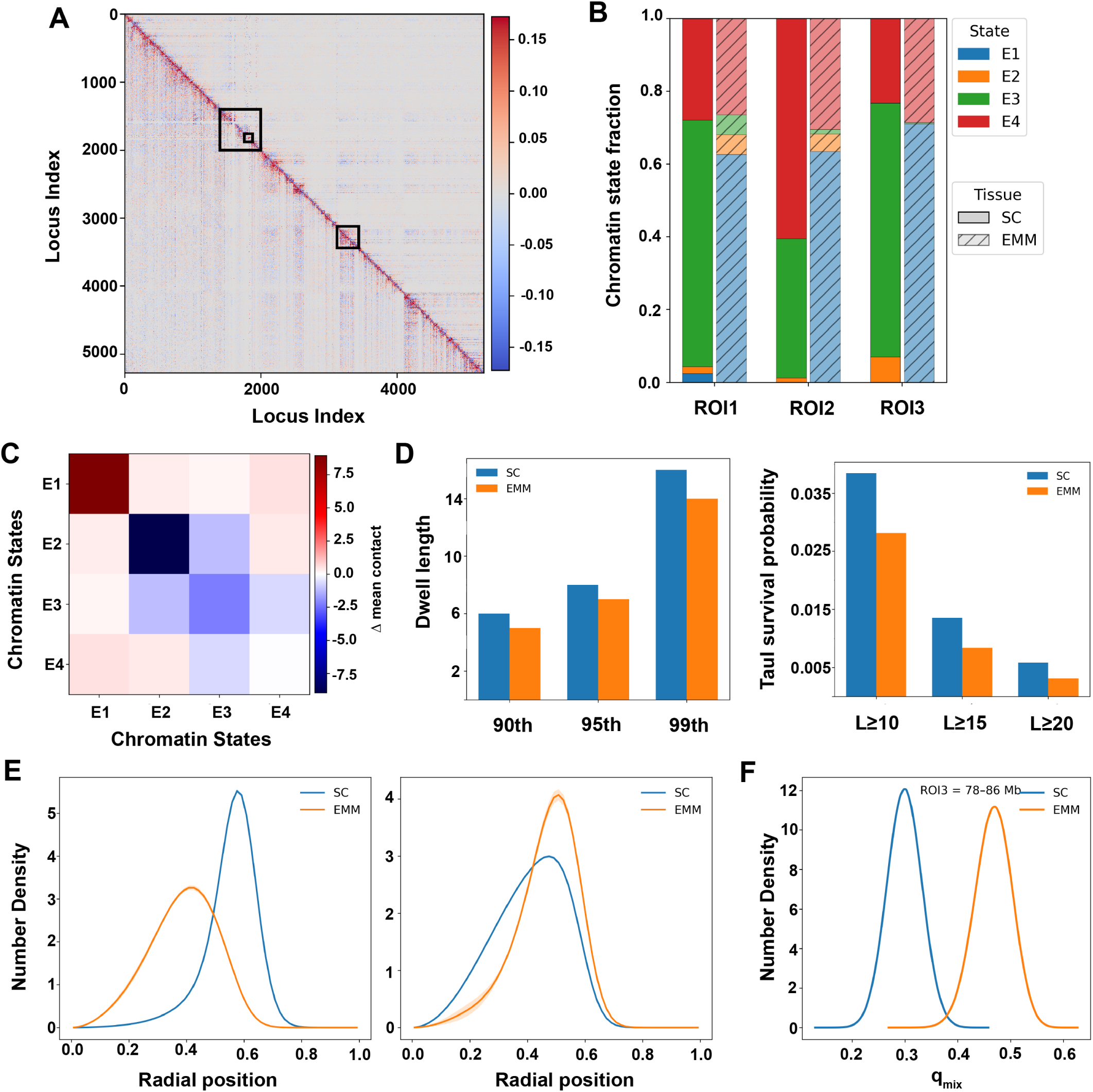
Tissue-specific epigenetic differences reshape chromosomal interactions, contact persistence, and conformational organization. **(A)** Locus-by-locus relative contact difference (EXP ↑ / SIM ↓) between colon and muscle; red indicates stronger colon contacts and blue stronger muscle contacts. **(B)** Stacked fractions of chromatin states E1–E4 for three regions of interest (ROI1–ROI3) in colon and muscle (experimental annotation). **(C)** State-interaction difference heatmap (colon − muscle) showing the change in mean contact between chromatin-state pairs E1–E4. **(D)** Dwell length at the 90th, 95th, and 99th percentiles (left) and tail survival probability for dwell durations L ≥ 10, 15, and 20 (right) for ROI1, colon versus muscle. **(E)** Density of normalized radial position (ρ = r/R) for E1 and E3 states, colon versus muscle. **(F)** Distribution of the mixing parameter q_mix for ROI3, colon versus muscle.

We next examined how compartment-level differences manifest at the polymer scale by calculating the contact probability distribution (P(s)) as a function of genomic distance, stratified by chromatin class (euchromatin and heterochromatin) for each tissue (Fig. SI 31C). In both SC and EMM, euchromatin and heterochromatin exhibited distinct scaling regimes, with heterochromatic contacts decaying more slowly over intermediate-to-long genomic distances, consistent with compartment-mediated phase separation. The two tissues diverged most strongly at intermediate genomic distances within each chromatin class, indicating tissue-specific shifts in polymer compaction rather than uniform rescaling of a shared decay profile. Notably, chromatin mobility remained comparable across tissues despite differences in the ensemble-averaged contact map. For example, ensemble-averaged mean-squared displacements for loci within RoI1 and RoI3 showed indistinguishable sub-diffusive scaling in SC and EMM (Fig. SI 31D). Because chain connectivity and excluded volume primarily govern backbone mobility, this similarity suggests that tissue-specific folding differences arise from changes in the effective energy landscape rather than from altered polymer dynamics. Therefore, both tissues are likely sampling distinct conformational basins on a topologically similar landscape.

To characterize tissue-specific chromosomal folding dynamics, we measured contact dwell times within RoI1 (**Fig. 5D**) and found that SC consistently exhibited longer-lived contacts than EMM across the distribution. For example, the 90th, 95th, and 99th percentiles of contact dwell times increased from 5.0, 7.0, and 14.0 frames in EMM to 6.0, 8.0, and 16.0 frames in SC, respectively. Likewise, the probability of contacts persisting beyond 10, 15, and 20 frames was higher in SC at all thresholds. These long-lived contacts indicate deeper energetic minima and stronger cooperative stabilization in SC, whereas EMM likely favors more transient interactions. Since locus mobility does not differ between tissues (**Fig. SI 31D**), these differences might reflect changes in interaction energetics rather than global dynamics.

We next assessed whether tissue-specific chromosomal folding coincides with differences in nuclear radial organization (**Fig. 5E**). In both tissues, heterochromatin localized preferentially toward larger normalized radii (*r/R*) relative to euchromatin, consistent with canonical radial segregation. However, SC exhibited sharper, more separated radial distributions, with higher peak densities and stronger euchromatin–heterochromatin separation, whereas EMM showed broader, more overlapping profiles with a modest inward shift of both chromatin classes (peak densities ≈ 4.05 and ≈ 3.30). These patterns place SC closer to a phase-separated regime and EMM closer to a partially mixed state. The contact-mixing order parameter *q_mix_* for RoI3 supports this conclusion (**Fig. 5F**). Overall, the mixed-state distributions of SC (*q_mix >_*∼0.3) and EMM (*q_mix>_* ∼0.47) showed minimal overlap, indicating stronger same-state clustering in SC and increased cross-compartment mixing in EMM within the same genomic region. Finally, the radius-of-gyration (*R_g_*) distribution analysis of the RoI1 conformational ensemble (**Fig. SI 32**) revealed distinct compaction regimes between the two tissues. In particular, SC sampled broader conformations with higher *R_g_* (peak at *R_g_* ≈ 5.7) than the compact and more uniform conformational ensembles (peak at *R_g_* ≈ 5.2) sampled in the EMM tissue. This observation suggests that even within a region of comparable epigenetic complexity, SC and EMM populate distinct conformational ensembles, consistent with their underlying tissue-specific energetic minima.

Taken together, these results indicate that tissue-specific chromatin organization of chromosome 10 in the ENTEx donor is associated with underlying differences in the composition of epigenetic states. These epigenetic variations extend to compartment-level contact preferences, polymer-scale scaling behavior, nuclear positioning, compartment mixing, and overall compaction. In contrast, the backbone mobility of model chromatin remains unchanged. Overall, the epigenetic-state–informed polymer model captures tissue-specific chromosomal folding within a unified framework and reproduces the contact maps of both SC and EMM, along with their structural differences.

## DISCUSSION

In this work, we used an epigenetic-state-informed polymer modeling framework to generate three-dimensional chromosomal conformational ensembles directly from cell-line- or tissue-type-specific epigenomic data^33,34,42,44^. We applied this framework to systematically investigate how genetic and epigenetic variations affect the chromosomal folding landscape. Toward this goal, we constructed and applied distinct polymer models to generate chromosomal conformational ensembles for three human cell lines (GM12878, H1-hESC, and IMR90), using chromatin states derived from reference epigenomes^64,65^. Subsequently, we benchmarked the simulation-derived ensemble-averaged properties of these cell lines against their experimentally derived Hi-C features. Overall, these models accurately captured genome organization features consistent with experimental Hi-C data, yielding Pearson correlation coefficients up to 0.97, low root-mean-square errors (0.004–0.017), and contact-probability scaling exponents close to experimental values^1^. Furthermore, comparison of chromosomal conformational ensembles revealed consistent polymer dynamics across cell lines. For example, we found global polymer dynamics indistinguishable among the three cell lines, with mean-squared-displacement exponents remaining sub-diffusive and statistically overlapping (α ≈ 0.48–0.49), indicating conserved backbone physics and comparable short-time dynamic constraints^44,47^. In contrast, we observed striking differences in chromosomal folding landscapes among the three cell lines that tracked with their underlying variation in chromatin-state composition and, more importantly, in chromatin state-specific interaction patterns^18,58,66^. For instance, we observed a diffuse conformational state profile characterized by strong inter-chromatin-state interactions and weak intra-chromatin-state associations, indicating a heterogeneous, weakly phase-separated chromosomal folding landscape in GM12878. In contrast, H1-hESC and IMR90 showed stronger enrichment of same-chromatin-state contacts along the diagonal of the interaction matrix, indicating more pronounced compartmental segregation^45,49,67^. These observations suggest that, despite similarities in polymer dynamics, a given chromosome across different cell lines occupies distinct folding regimes. Together, these findings indicate that effective interactions between chromatin states, rather than changes in the polymer chain’s mechanical properties, primarily shape cell-type-specific chromosomal folding.

Beyond quantifying ensemble-average metrics, our study also examined conformational dynamics and energetic heterogeneity across distinct cell lines and tissue types. For instance, we observed that large differences in contact persistence across cell lines arise from rare, long-lived contacts rather than from the overall contact distribution. These differences were compartment-specific, with euchromatin contacts persisting slightly longer in GM12878, whereas heterochromatin contacts were progressively stabilized in IMR90 and, most prominently, in H1-hESC. The observed compartmental asymmetry increased toward the tail of the distribution, indicating that the most persistent contacts showed the greatest compartment-specific separation. Similarly, differences in the upper quantiles of dwell-time distributions and in long-lived-contact survival fractions across cell lines were most pronounced between compartments in a cell-type-specific manner. These observations extended across the genome, with a consistent ordering of contact persistence (GM12878 < IMR90 < H1-hESC) and the largest separation in the extreme tail (Q99). These findings suggest relatively compact, highly segregated conformations with more persistent contacts in the IMR90 and H1 cell lines. In contrast, the mixed organization of GM12878 conformations, with stronger inter-chromatin state interactions, is consistent with a chromosomal folding energy landscape that permits more frequent conformational reorganization. Furthermore, a systematic parameter-sweep analysis of polymer potential energy functions revealed the interaction-range cutoff as the principal regulator of chromosome organization. Together, these results indicate that chromatin architecture emerges from the chromatin-state interaction network, where rare but persistent contacts potentially deepen energetic basins that stabilize segregated conformations and promote chromatin compartment segregation^38,45,67^.

Prior studies have shown that SVs can perturb 3D genome organization and rewire gene regulation by disrupting TAD boundaries and altering enhancer-promoter interactions^14,19^. However, these studies have primarily examined the impact of SVs on ensemble-average 3D genome structure^41,52,68^. A systematic assessment of how SVs influence chromosomal conformational landscapes remains underexplored. To address this gap, we applied our polymer modeling framework to comprehensively characterize differences in chromosomal conformational ensembles induced by a previously reported chromosome 8 deletion that overlaps a TAD boundary^53^. Consistent with our analyses of other human cell lines, the polymer model accurately reproduced both wild-type and SV-affected ensemble-averaged Hi-C contact maps (ρ = 0.97 in both systems) without re-optimizing interaction parameters beyond introducing the deletion itself ^41^. This finding suggests that the SV-induced change in chromosomal architecture is sufficient to recapitulate much of the observed structural reorganization.

Comparison of chromosomal conformational ensembles between wild-type and SV-affected models revealed a substantial redistribution of chromosomal folding states. For instance, within the wild-type ensemble, 65% of conformations adopted predominantly insulated states, whereas 68% of SV-affected conformations occupied a fused folding state. The observed transition in the SV-affected chromosomal ensemble can be attributed to an increased persistent contact fraction in fused conformations and a shift in the dominant structural basin along the breakpoint-flanking coordinate. Furthermore, the SV-affected ensemble sampled more extended structures, exhibiting a larger mean radius of gyration (1.0nm versus 0.88nm) and end-to-end distance (2.3 versus 1.7) than the wild-type ensemble. These observations align with prior studies showing that boundary-disrupting deletions weaken domain insulation and promote interactions between neighboring chromatin regions^69,70^. A systematic shift in the SV-affected model’s conformational ensemble relative to the wild-type was accompanied by selective rewiring of chromatin-state interactions. For example, we observed the strongest gains among active chromatin states, with E1-E1 and E2-E2 contacts showing the largest increases in the SV-affected ensemble, whereas E3-E5 states exhibited substantially smaller changes. Moreover, we found positive increases in interactions between active and intermediate chromatin states, particularly E1-E2 contacts. These patterns suggest that the modeled deletion preferentially reorganizes interactions among transcriptionally permissive chromatin regions rather than uniformly increasing contact frequency across the locus^71,72^. The observed enrichment of active chromatin interactions aligns with prior studies examining the consequences of boundary disruption and enhancer rewiring following genomic rearrangements^69,70,73^.

A close examination of regulatory element interactions, including contacts potentially affected by the deletion, revealed heterogeneous gene-level effects, even though all four genes reside near the same deletion breakpoint. For example, TIGD5, PYCR3, and GFUS exhibited modest reductions in enhancer contacts across conformations. In contrast, ZNF623 showed a substantially larger loss (Δ = −0.140), nearly an order of magnitude greater than the decreases observed for the other three genes. The observed variability in distal element interactions was accompanied by marked variation in local regulatory context. In particular, within a ±10-bead window, genes such as GFUS, PYCR3, and TIGD5 had approximately three- to fourfold higher densities of nearby active regulatory elements than ZNF623. Consistent with this difference, these genes retained 81-86% of their wild-type contacts following the deletion, whereas ZNF623 retained only 54% of its local contacts. Similar context-dependent responses have been observed in studies of enhancer hijacking and TAD boundary disruption, where the regulatory consequences of structural variation depend strongly on the local distribution of cis-regulatory elements^69,73,74^. Together, these findings suggest that SVs can influence the conformational states available to a chromosomal region, with measurable shifts in chromatin-state populations leading to differential degrees of enhancer contact disruption across neighboring genes.

Beyond investigating the impact of a deletion, we systematically assessed how epigenetic perturbations influence chromosomal folding. We analyzed differences in the folding landscape of chromosome 10 in sigmoid colon (SC) and esophageal muscularis mucosa (EMM) tissues from the same ENTEx donor to isolate epigenetic differences from genomic variation. As with our SV analyses, we reconstructed and compared the chromosomal folding landscapes for each tissue by integrating tissue-specific epigenetic annotations into our polymer modeling framework. Despite overall similarity in contact maps between tissue types (Pearson correlation: *ρ* ∼0.96), the largest differences were concentrated in discrete genomic regions that also showed substantial shifts in chromatin-state composition. A close examination of epigenetic states within these regions revealed a higher propensity for homotypic interactions (E1-E1) in SC, whereas EMM showed a higher propensity for homotypic interactions (E2-E2) and various heterotypic interactions (E1-E3 and E2-E3), suggesting that tissue-specific epigenetic states alter the interaction network underlying chromosome folding.

Importantly, comparing SC and EMM chromosomal folding ensembles revealed differences that were not readily apparent in ensemble-averaged contact maps. For example, despite nearly identical chromatin mobility, SC conformations consistently exhibited longer-lived chromatin contacts than EMM and higher probabilities of contact persistence across all measured thresholds. These observations indicate that the two tissues occupy distinct folding landscapes with different contact stabilities, consistent with recent studies suggesting that rare and persistent interactions contribute disproportionately to higher-order genome organization and may be an important determinant of tissue-specific chromatin structure. Furthermore, tissue-specific conformational ensembles differed in the extent of chromatin segregation and structural heterogeneity. For instance, the SC conformational ensemble exhibited stronger radial separation of euchromatin and heterochromatin, lower compartment mixing (<Q_mix_> ≈ 0.30 versus ≈ 0.47 in EMM), and enhanced same-state clustering. At the same time, SC sampled a broader conformational ensemble, with a larger radius-of-gyration peak (R_g_ ≈ 5.7nm) than EMM (R_g_ ≈ 5.2nm). Together, these observations suggest that tissue-specific epigenetic states influence chromosomal folding not only by altering contact frequencies but also by reshaping the ensemble of accessible conformations, their contact persistence profile, and the degree of compartmental segregation.

Although our polymer modeling framework provides mechanistic insights into chromosomal folding landscapes, several limitations warrant consideration. First, we optimized the model to reproduce population-averaged Hi-C contact maps within an equilibrium framework^38,75^. Consequently, the conformational ensembles, contact dwell times, and persistence measures reported here describe properties of the inferred folding landscape rather than direct measurements of in vivo chromatin dynamics. Moreover, we express dwell times and persistence measures in simulation units and do not calibrate them to physical timescales. Incorporating non-equilibrium mechanisms, such as loop extrusion and active chromatin remodeling, could help bridge the gap between inferred structural landscapes and biological dynamics. Second, we infer conformational heterogeneity from polymer ensembles rather than observing it directly. Because Hi-C captures population-averaged contact frequencies, multiple structural ensembles can be consistent with the same contact map. Future integration with single-cell Hi-C^25,27^, multiplexed imaging^28,76,77^, and live-cell chromatin tracking^30,78^ will be important for validating key model predictions, including distinct conformational basins and rare persistent contacts. Third, we examined a limited set of biological scenarios, including a single structural variant, a single donor, and a single tissue pair, each restricted to one genomic region or chromosome. We prioritized depth of conformational sampling over breadth, focusing on well-characterized systems with independent experimental evidence to anchor model predictions, such as a deletion previously shown to disrupt TAD architecture and paired tissues sharing an identical genetic background. Finally, our current approach does not explicitly model transcriptional output but uses chromatin contacts as structural proxies for regulatory potential^74,79,80^. Expanding the current study to additional donors, tissues, disease states, and diverse classes of SVs, while integrating structural ensembles with transcriptional and imaging measurements^76,77^, will help establish the generalizability and functional relevance of the relationships observed here between epigenetic state, chromatin folding, and regulatory rewiring.

Despite these limitations, our study uses a polymer modeling framework that links epigenetic information to chromosomal folding landscapes across cell types and tissues. By bridging quantitative epigenomics and polymer physics, the polymer model provides a mechanistic explanation for how one-dimensional chromatin states give rise to higher-order genome organization and serves as a practical tool for investigating the structural consequences of genetic and epigenetic perturbations. More broadly, these results highlight the potential of ensemble-based polymer modeling to complement experimental genomics by revealing the conformational heterogeneity and folding landscapes underlying observed 3D genome architecture.

## MATERIALS & METHODS

### Datasets

We obtained functional genomics and Hi-C contact map datasets from publicly available sources, including the 4D-nucleome, ENCODE, and ENTEx projects, to perform our systematic polymer simulations. Specifically, we retrieved Hi-C contact maps for multiple human cell lines, including GM12878, IMR90, and H1-hESC, from the 4D-nucleome project^61,81^. For each cell line, we extracted normalized contact matrices for all autosomal chromosomes (1-22) using the Juicer Tool^82^. We generated these normalized contact matrices with the Knight–Ruiz (KR) algorithm, which corrects for systematic biases, including coverage variability and genomic mappability. We also obtained chromatin state annotations and CTCF binding profiles from the ENCODE Project Consortium database for all three cell lines to define the epigenetically informed polymer model state. We used histone modification ChIP-Seq datasets mapped to the human reference genome Build 38 (GRCh38) for each cell line to infer the corresponding genome-wide chromatin state with the ChromHMM tool. Moreover, we used CCCTC-binding factor (CTCF) ChIP-seq peaks to delineate loop extrusion anchor sites^83,84^ in our polymer model.

For SV analysis, we used Hi-C data from one HGSVC sample, HG00864, which carries a previously characterized large deletion that affects the TAD boundary^53^. We extracted an intra-chromosomal contact map for chromosome 8 from this sample at 5-*kb* resolution using the Juicer Tool to simulate wild-type and mutant polymers (after modeling the deleted region). Similarly, we used Hi-C contact maps, CTCF binding profiles, and histone modification ChIP-Seq data for Sigmoid Colon (SC) and Esophagus Muscularis Mucosa (EMM) tissues from a single donor (donor ID: ENCDO845WKR) in the ENTEx resource^5^. These tissue-specific epigenomic and chromosomal contact profiles from the same donor enabled us to perform polymer simulations to systematically investigate tissue-specific chromatin organization and folding.

### Chromosomal Representation

We implemented a coarse-grained polymer modeling framework to systematically model the chromosomal folding process. We divided each chromosome into fixed-size, non-overlapping genomic bins, each represented by a single bead/monomer in our polymer model. We defined a chromatin state for each genomic bin and, by extension, for its corresponding monomer bead, using ChromHMM-based annotations^59,65^. Briefly, ChromHMM uses histone modifications and chromatin accessibility data (obtained via DNase-seq/ATAC-seq) to define genome-wide chromatin status for each genomic bin. In this framework, the polymer backbone maintains chromatin bead connectivity by linking adjacent monomers, whereas three-dimensional chromosomal conformations arise from biophysical interaction potentials (see supplemental method). Incorporating chromatin-state-aware interaction strengths into our potential energy function enables chromosomal compartmentalization, sub-compartmentalization, and other dynamic structural features observed during chromosome folding. To capture chromatin state for polymer interactions, we defined six model-specific epigenetic classes (E1-E6), corresponding to various levels of compaction and transcriptional activity based on histone marks (including H3K4me1, H3K9me3, H3K27ac, H3K27me3, etc.). For instance, the E1 state corresponds to highly accessible chromatin enriched for promoter and strong enhancer marks, which is transcriptionally active, gene-dense, and structurally open. In contrast, the E2 state is characterized by moderately active chromatin that is less accessible than in E1 but still transcriptionally permissive. Similarly, the E3 state is assigned to regions enriched in H3K27me3, consistent with facultative heterochromatin, whereas E4 regions indicate compacted heterochromatin. The E5 state includes regions with actively transcribed gene bodies and elongated chromatin structure. Finally, the E6 chromatin state corresponds to centromeres, which form very tightly packed structures and tend to group together dynamically.

Beyond chromatin states, we delineated CTCF binding sites that overlap with beads in our polymer model using their genomic coordinates. More specifically, we defined loop anchors as pairs of convergently oriented CTCF sites for each cell line and tissue type, which impose constraints consistent with loop extrusion-mediated chromosome folding. Including these anchors enables the formation of loops and domain-like structures that mirror the chromatin organization observed in experimental Hi-C data^1,40,41^. We used ChIP-seq datasets for each cell line to identify CTCF binding sites by filtering peak coordinates to retain only the most reliable sites. A CTCF label was assigned to a bead when a peak overlapped a genomic bin. In cases where peaks overlapped a single bead, they were merged into a single effective binding site. Previous studies show that CTCF motif orientation is critical for loop extrusion-mediated chromosomal folding. Therefore, we accounted for motif orientation when defining loop anchors, using convergent CTCF binding sites with a forward motif upstream and a reverse motif downstream. These anchors delineate specific loci that can stabilize chromatin loops, aligning with experimental findings of loop enrichment between convergent CTCF motifs^85^. Overall, our bead/monomer representation integrates genomic sequence connectivity, epigenetic state identity, and CTCF-mediated architectural constraints into a unified polymer framework that enables the physical modeling of the three-dimensional organization of chromosomes.

### Molecular Dynamics Simulations

We implemented our modeling framework in Python 3.11 and OpenMM^86^ to perform molecular dynamics simulations and generate ensembles of three-dimensional chromosomal conformations using the polymer and genomic energy functions (described in the supplementary information). Briefly, we represented each chromosome as a bead-on-spring polymer with a resolution of 50kb per bead, unless otherwise specified. Each bead corresponds to a chromatin segment of size σ, and we set the bead diameter to 1, defining the system length scale in model units. We applied periodic boundary conditions to avoid technical artifacts and allow unrestricted chromatin motion. We placed chromatin, modeled as chains of beads, in a spherical box, ensuring a volume fraction of 0.1^87^. Furthermore, we included CTCF loop coordinates and chromatin states derived from experimental datasets as inputs to label each bead in the chromatin chains.

We generated initial chromosome conformations as random-walk polymer chains and then equilibrated the system using a temperature-annealing protocol. First, we collapsed the system at a high temperature T = 5 in reduced units to randomize the initial structure. We then applied the annealing procedure by gradually reducing the temperature from T = 5 to T = 1. At T = 1, the chromosome remains folded but dynamically fluctuates, reproducing experimental Hi-C maps^42^. This approach enabled us to generate independent, uncorrelated configurations that sample a broader chromatin conformational landscape. We performed these simulation runs using Langevin dynamics with a damping coefficient of 10.0 *τ*, where *τ* is the simulation time unit. We applied a time step of Δ*t* = 0.01*τ* for all simulations. To recapitulate Hi-C contact maps, each sampling simulation for any chromosome in all three cell lines was run for 5×10^8^ steps, and chromosome conformations were stored every 10^3^ steps. This procedure generated 500,000 conformations per simulation run. We performed 30 independent production runs, yielding 15×10^6^ sampled conformations per chromosome for downstream structural analysis.

#### Parameter Optimization Simulations

We used an iterative parameter optimization approach to determine the parameters that best reproduce experimental Hi-C contact maps. To this end, we performed 40 independent runs to generate chromosomal conformations for each chromosome. Each run produced 50000 conformations, yielding 2×10^6^ structural conformations used to generate an ensemble-average-based *in silico* Hi-C contact map. We iteratively updated the interaction parameters in our potential energy function until the Pearson correlation coefficient between the simulated and experimental maps exceeded 0.90, ensuring quantitative agreement with the experiments. The 26 optimized parameters include the chromatin-state interaction (*M*_ij_), the genomic separation compaction parameter (*G*_i_), and additional polymer parameters (see Supplement Table 1). The optimal parameters *ζ* and *r*_c_ were 3.22 and 1.78, respectively, to reproduce experimental Hi-C maps^42,60^.

#### Unfolded chromosome simulations

We also performed additional simulations to explore unfolded chromosomal conformations by varying the parameters of Eq. 8 (see Supplementary Information). In particular, we varied ζ and r_cut_ over the ranges ζ = 3.00 − 3.25 and rcut = 1.50 − 1.78, while the optimal values for reproducing experimental Hi-C maps were ζ = 3.22 and rcut = 1.78. This switching protocol yields partially unfolded chromosomal conformations. These simulations were performed for chromosome 10 across all three cell lines (GM12878, IMR90, and H1-hESC) using the same molecular dynamics protocol described above. Each system included 30 production runs, each with 5 × 108 steps, yielding 15 × 106 conformations per condition. We used these ensembles to analyze large-scale chromatin expansion and transitions between folded and unfolded chromosomal states.

#### Structural Variation Simulations

To investigate the effects of an SV, we performed an additional set of polymer simulations for chromosome 8 in the HGSVC sample HG00864, which contains a deletion. The simulations were restricted to a genomic region spanning 140-145 Mb, which, at a resolution of 5 kb per bead, produced a polymer chain of 1000 beads. After removing the genomic segment (chr8: 143618315-143632522)^53^, which maps to four consecutive beads (723-726) in the SV-affected polymer representation of chromosome 8, we reconstructed the truncated polymer chain by removing these beads and reconnecting the neighboring segments to preserve chain continuity. Because we lacked matching ChIP-seq data for the rearranged sample, we used chromatin-state annotations from the wild-type GM12878 dataset as a proxy, since GM19036 and HG00864 are both lymphoblastoid cell lines. We optimized parameters with 40 independent replicas, generating 2×10^6^ conformations, followed by 30 production simulations, each with 5×10^8^ integration steps, yielding 15×10^6^ conformations for both wild-type and truncated chromosomes. We used these ensembles to directly compare structural organization and in silico Hi-C contact maps, enabling quantitative analysis of how the deletion affects the local and global structural properties of the resulting chromatin conformational ensembles.

#### Tissue-specific simulations

We performed molecular simulations of chromosome 10 in sigmoid colon (SC) and Esophagus Muscularis Mucosa (EMM) tissues from the same individual in the ENTEx project to investigate tissue-specific differences in chromosomal folding. We represented chromosome 10 at a 25-kb-per-bead resolution, constructing a polymer chain of 5,277 beads. We incorporated tissue-specific epigenomic and chromatin organization data from ENTEx using the same protocol as in the cell line simulations, including ChromHMM-derived^59,65^ chromatin state annotations and CTCF binding site information. We obtained ChromHMM segmentations from histone modification data (H3K4me1, H3K27ac, H3K4me3, H3K27me3, H3K9me3, and H3K36me3) and mapped these emission states onto the chromatin classes used in the polymer model. States enriched for H3K4me3 and promoter-associated annotations defined the active promoter class (E1), whereas states enriched for H3K4me1 and H3K27ac defined the active enhancer/open chromatin class (E2). We assigned Polycomb-repressed regions enriched in H3K27me3 to E3 and constitutive heterochromatin enriched in H3K9me3 to E4. We independently assigned centromeric regions to a centromeric state (E5), treating them separately from ChromHMM-derived epigenetic classes. This mapping directly links tissue-specific epigenetic segmentation to chromatin-state–dependent interaction potentials in the simulations.

We obtained tissue-specific CTCF binding sites from ChIP-Seq datasets and mapped them onto the polymer to define loop-anchor positions. Using pairs of convergently oriented sites, we imposed pairwise loop constraints to reconstruct the loop extrusion term in the effective energy function (**SI Eq. 13**), thereby capturing tissue-specific looping interactions consistent with experimental data. We performed 40 independent optimization runs per tissue (yielding 2 × 10^6^ in total) to achieve quantitative agreement with the corresponding Hi-C maps. After optimization, we carried out 30 independent production simulations per tissue, each comprising 5×10^8^ integration steps and using the same sampling frequency as in the main simulations. The resulting ensembles (15×10^6^ chromosomal conformations per tissue) enabled quantitative analysis of tissue-specific differences in chromosome 10 folding, compartment organization, and contact map structure.

### Downstream analyses

We computed a comprehensive set of observables from equilibrated molecular dynamics trajectories to quantitatively characterize chromatin organization, structure, and dynamics. This enabled us to investigate the association between simulated conformations and underlying biological mechanisms. For instance, we constructed a state–state interaction matrix directly from the simulation trajectories to quantify effective interactions among chromatin states (E1–E6). This approach captured the energetic contributions encoded in the effective energy function, including epigenetic interactions, long-range potentials, and loop-extrusion constraints, rather than relying solely on contact-based Hi-C measures. Similarly, we characterized chromatin dynamics by computing the mean squared displacement (MSD) using a fast Fourier transform–based implementation for numerical efficiency over long trajectories. We also quantified global chromosomal compactness using the radius of gyration and end-to-end distance. To assess structural heterogeneity in regions affected by structural variants, we identified distinct conformational states in the simulation trajectories and quantified their population frequencies and lifetimes. We performed these analyses separately for euchromatic and heterochromatic loci, enabling compartment-specific characterization of chromatin structure and mobility. Across all analyses, we averaged each observable across independent replicas of the same simulation to ensure statistical robustness.

Furthermore, we biologically contextualized the simulation-derived structural and dynamic changes by annotating the modeled region with gene and regulatory element information. The interval spanned chr8:140,000,000-145,000,000, corresponding to a 5 Mb genomic region represented at 5,000 bp per bead, for a total of 1,000 beads in the model. We first identified protein-coding genes within and near the deletion region using GRCh38 gene annotations downloaded from Ensembl (Release 115)^88^. We then used SCREEN v4 to retrieve candidate cis-regulatory elements (cCREs) overlapping these gene regions and retained cCREs linked to the corresponding genes in SCREEN through linked-gene evidence^63^. SCREEN provides access to a unified ENCODE cCRE registry that integrates chromatin accessibility, histone modification, and CTCF-binding data from experiments across diverse cell and tissue contexts. We classified these cCREs into regulatory categories, including promoter-like signatures, proximal enhancer-like signatures, distal enhancer-like signatures, CTCF-only elements, and DNase-H3K4me3 elements. Each gene and cCRE annotation was mapped to its corresponding 5 kb simulation bead based on genomic coordinates, allowing regulatory features to be summarized at bead resolution. We then used these bead-level biological annotations as metrics to compare with simulation-derived observables, including changes in bead interactions, chromatin-state contacts, mobility, compactness, and conformational-state behavior. This framework allowed us to assess whether regions showing altered structural or dynamic properties in the simulation were enriched for, depleted of, or spatially associated with gene-linked regulatory elements, thereby linking polymer-level chromatin behavior to potential biological consequences of the structural variant.

## Source code & Data Availability

The processed HI-C data generated by the Human Genomic Structural Variation Consortium (HGSVC) were obtained from the IGSR data portal (https://ftp.1000genomes.ebi.ac.uk/vol1/ftp/data_collections/HGSVC2/working/20230515_Shi_hic_files/) ^53^. 4D-nucleome data including Hi-C contact maps for multiple human cell lines GM12878, IMR90, and H1-hESC was obtained from the 4D-nucleome data portal (https://data.4dnucleome.org/). ENCODE data were accessed through the SCREEN (v4) portal (https://screen-v4.wenglab.org/). ENTEx data related to sigmoid colon (SC) and esophageal muscularis mucosa (EMM) were obtained from the same ENTEx donor (donor ID: ENCDO845WKR) from the ENTEx portal (http://entex.encodeproject.org/)

All relevant source code for this study, including scripts for molecular dynamics simulations and downstream analyses, can be accessed on the project GitHub page (https://github.com/kumarlab-compomics/ChromoDyn)

## Funding

SK, AT, and SB acknowledge support from the Natural Sciences and Engineering Research Council of Canada, Princess Margaret Cancer Foundation, and Canada Research Chair Program.

## Conflict of Interest

The authors declare that they have no conflict of interest.

## Acknowledgements

This research was made possible by the availability of Hi-C data and high-quality, haplotype-resolved genome assemblies generated by the Human Genomic Structural Variation Consortium (HGSVC) and the ENTEx project. We also acknowledge helpful discussions with Drs. Timur Galeev and Mark Gerstein regarding tissue-specific Hi-C data from the ENTEx project.

## Notes

### Competing Interest Statement

The authors have declared no competing interest.

