## supplementary information for "Epigenome-informed Polymer Modeling Reveals Chromosome Folding Landscapes across Distinct Genetic and Epigenetic Contexts"

### Supplementary Methods

**Polymer Energy Function:** $U_p$

To maintain polymer connectivity and shape, regulate local binding rigidity, and also impose excluded volume, a coarse-grained bead-spring polymer model has been featured with four energy components: harmonic bond, harmonic angle, a soft core repulsive nonbonded potential, and the confinement potential as below,

$$U_p = U_{\text{bond}} + U_{\text{angle}} + U_{\text{soft}} + U_{\text{confine}}. \quad (1)$$

Equation 1 ensures the chromatin fiber stays connected, is sufficiently flexible, and maintains the excluded volume effect. A harmonic spring potential connects adjacent beads on the chromatin chain.

$$U_{\text{bond}} = \Sigma_{i=1}^{N-1} \frac{k_{\text{b}}}{2} (r_{\text{i,i+1}} - r_0)^2, \quad (2)$$

where $N$ is the number of beads, $r_{\text{i,i+1}}$ is the distance between consecutive beads, $k_{\text{b}}$ is the stiffness bond constant, $r_0$ is the equilibrium bond length equal to $\sigma$ as the bead's diameters. The chromatin fiber's local bending rigidity is implemented through a harmonic angle potential applied to triplets of consecutive beads.

$$U_{\text{angle}} = \Sigma_{i=2}^{N-1} \frac{k_\theta}{2} (\theta_{\text{i}} - \theta_0)^2, \quad (3)$$

where $\theta_{\text{i}}$ is the angle formed by three consecutive beads, $k_\theta$ is the angular constant, and $\theta_0 = \pi$ is the equilibrium angle.

To prevent beads from overlapping while still letting chromatin rearrangement happen smoothly, a soft-core repulsive interaction is used between all nonbonded beads. This interaction is characterized as a truncated repulsive Lennard-Jones potential integrated with a soft-core region with a minimal distance,

$$U_{\text{soft}}(r) = \begin{cases} \epsilon U_{\text{LJ}}(r) + \frac{\epsilon}{4}, & r_0 < r < r_{\_c} \\ \frac{E_{\text{cut}}}{2}[1 + \tanh(\frac{2U_{\text{LJ}}(r)}{E_{\text{cut}}} - 1)], & r \le r_0 \\ 0, & r \ge r_{\text{c}} \end{cases} \quad (4)$$

where,

$$U_{LJ}(r) = (\frac{\sigma}{r})^{12} - (\frac{\sigma}{r})^6, \quad (5)$$

and $r$ is the distance between beads, $\epsilon$ is the repulsive strength, $E_{\text{cut}}$ controls the soft-core overlap energy, and $r_{\text{c}} = 2^{\frac{1}{6}}\sigma$ is the cut off for purely repulsive forces. The tanh function smooths out the interaction energy at short distances, which stops numerical instabilities that would happen because the $U_{\text{LJ}}$ decays rapidly. Therefore, equations 2-4 define the energy of the homopolymer backbone that is used to mimic chromatin fiber.

To simulate nuclear confinement, chromatin beads were restricted within a spherical volume with a Lennard-Jones wall potential on each bead. This confinement is determined in relation to the radial distance from the nuclear center. The confinement energy is written as

$$U_{\mathrm{conf}} = \Sigma_{i=1}^{N}\Theta(r_{\mathrm{c}} - \Delta r_{\mathrm{i}})[\epsilon U_{\mathrm{LJ}} + \epsilon/4], \tag{6}$$

where $R$ is the confinement radius, and $\Theta$ is the Heaviside step function that truncates the interaction at the purely repulsive cutoff $r_c$, and $\Delta r_i$ is the distance between beads. The confinement radius is determined based on the chromatin density in the nucleus [17].

$$R = (\frac{3N}{4\pi\rho})^{1/3}, \tag{7}$$

where $\rho$ is the base pair density in the nucleus. This formulation keeps the chromatin volume fraction the same in all simulations.

**Biophysical Chromosome Potentials**

To elucidate chromosome folding beyond polymer connectivity, we define three sequence-informed interaction terms that explain chromatin identity compatibility, genome-distance-dependent compaction, and CTCF-mediated looping. These contributions are known as Chromatin Identity Coupling (CIC), Genomic Separation Compaction (GSC), and Anchor-mediated Loop Stabilization (ALS). All interactions are modulated by distance with the contact-switching function that promotes spatial proximity while preserving distinguishable forces,

$$f(r_{ij}) = 0.5[1 + \tanh\zeta(r_c - r_{ij})], \tag{8}$$

where $r_{ij}$ denotes the spatial distance between loci $i$ and $j$, $r_c$ characterizes the contact distance, and $\zeta$ controls the sharpness of the transition. This hyperbolic-tangent form smoothly interpolates between contact and non-contact regions, avoiding discontinuities while preserving a physically meaningful interaction scale. Hence, the total chromosome-specific interaction energy is written as

$$U_{\mathrm{chrom}} = U_{\mathrm{CIC}} + U_{\mathrm{GSC}} + U_{\mathrm{ALS}}. \tag{9}$$

***Chromatin Identity Coupling ($U_{CIC}$)***

The Chromatin Identity Coupling term models interactions between loci based on their chromatin states ($E_1$–$E_6$). Each bead is assigned to a chromatin identity $t_i$ and interactions between bead pairs depend on a symmetric identity coupling matrix $M_{t_i t_j}$. The resulting contribution to the energy is

$$U_{\mathrm{CIC}} = \Sigma_{i<j} M_{t_i t_j} f(r_{ij})\Theta(r_{ij} - r_{\mathrm{lim}}), \tag{10}$$

where Θ denotes the Heaviside step function, and $r_{\text{lim}}$ defines the lower bound preventing overlap-driven contribution. The Matrix $M$ determines the effective interaction between chromatin classes and reflects their spatial association, driving compartment-like segregation and organizing chromatin into structurally distinct domains.

*Genomic Separation Compaction ($U_{GSC}$)*

To account for long-range folding dependency of chromatin identity, we introduce a genomic separation-dependent interaction. For loci separated by genomic distance

$$d_{ij} = |i - j|$$

, where the interaction strength is defined as

$$G(d_{ij}) = \frac{G_1}{\log(d)} + \frac{G_2}{d^2}. \quad (11)$$

Therefore, the corresponding energy term will be written as

$$U_{GSC} = \Sigma_{i<j} G(d_{ij}) f(r_{ij}) \Theta(d_{ij} - D_{L1}) \Theta(D_{L2} - d_{ij}), \quad (12)$$

where $D_{L1}$ and $D_{L2}$ define the lower and upper bounds that limit the range of interaction, respectively. This term captures baseline polymer folding of chromosomes, imposing genomic-distance-dependent compaction that reproduces the decay of contact probability with increasing genomic distance. The combination of logarithmic and power-law effects provides a flexible representation of hierarchical chromatin organization.

*Anchor-Mediated Loop Stabilization ($U_{ALS}$)*

Specific looping interactions between genomic loci are incorporated through the Anchor-Mediated Loop Stabilization term. Given a set of loop anchor pairs

$$L = (i,j),$$

the energy term is

$$U_{ALS} = \Sigma_{(ij)\in L} \xi f(r_{ij}), \quad (13)$$

where $\xi$ controls the strength of loop stabilization, set to -1.61 in the model unit [18]. This interaction provides an effective representation of CTCF/cohesin-mediated loop formation. Loop anchors are stabilized without enforcing fixed contacts, allowing structural flexibility while obtaining loop architecture. The set of anchor pairs has been determined from experimentally identified CTCF binding sites.

**Parameter Optimization**

We used the Maximum Entropy principle to determine the interaction parameter that governs chromatin folding. This framework finds the least-biased energy landscape that fits with the experimentally measured Hi-C contact probabilities. This method allows for the direct construction of an efficient chromosomal Hamiltonian from data, while maintaining the constraints of polymer physics. The principle has been widely used in previous studies for chromosome modeling to provide statistically rigorous inference of interaction parameters from experimental observables [18–20]. Within this framework, the chromatin conformational ensemble is described by a Boltzmann distribution, in which effective energy includes both the polymer Hamiltonian and data-driven interaction terms (See Biophysical Chromosome Potential). The latter are parametrized through the epigenetic interaction matrix $M$, which encodes pairwise affinities between chromatin states. Particularly, the optimization seeks the matrix $M$ and $G$s parameters that produce simulated contact probabilities consistent with experimental observations while maximizing configurational entropy of the ensemble. At the start of optimization, all parameters are set to zero, ensuring no interaction bias is imposed. Using molecular dynamics simulations, chromatin conformations are then generated, and contact probabilities are computed from the conformational ensemble. Using the Root Mean Square Error (RMSE) and Pearson correlation coefficient, the discrepancy between simulated and experimental HiC maps are used to iteratively update both $M$ and $G$ parameters. This method is repeated until convergence, yielding a self-consistent parameter set that reproduces the experimental contact map. In total, 26 parameters have been optimized to regenerate experimental Hi-C contact maps, and all are listed in Table 1.

| Interaction Parameters | Cell line | | | Structural Variation | | Tissues | |
|---|---|---|---|---|---|---|---|
| | GM12878 | IMR90 | H1-hESC | WT | DEL | SC | EMM |
| E1-E1 | -0.268028 | -0.268543 | -0.268098 | -0.268028 | -0.257779 | -0.254883 | -0.257970 |
| E1-E2 | -0.274604 | -0.274353 | -0.274222 | -0.274604 | -0.265322 | -0.271662 | -0.276986 |
| E1-E3 | -0.262513 | -0.262766 | -0.262871 | -0.262513 | -0.252753 | -0.269568 | -0.260876 |
| E1-E4 | -0.258880 | -0.258232 | -0.258156 | -0.258880 | -0.251852 | -0.251840 | -0.252646 |
| E1-E5 | -0.266760 | -0.266676 | -0.266107 | -0.266760 | -0.257831 | -0.269348 | -0.267065 |
| E1-E6 | -0.225646 | -0.225132 | -0.225100 | -0.225646 | -0.216645 | NA | NA |
| E2-E2 | -0.299261 | -0.299783 | -0.299102 | -0.299261 | -0.289202 | -0.306707 | -0.296779 |
| E2-E3 | -0.286952 | -0.286232 | -0.286907 | -0.286952 | -0.275725 | -0.298147 | -0.280189 |
| E2-E4 | -0.281154 | -0.281644 | -0.281090 | -0.281154 | -0.278087 | -0.293041 | -0.283196 |
| E2-E5 | -0.301320 | -0.301890 | -0.301323 | -0.301320 | -0.292391 | -0.293045 | -0.304625 |
| E2-E6 | -0.245080 | -0.245322 | -0.245235 | -0.245080 | -0.237766 | NA | NA |
| E3-E3 | -0.342020 | -0.342432 | -0.342232 | -0.342020 | -0.331936 | -0.344593 | -0.338169 |
| E3-E4 | -0.321726 | -0.321544 | -0.321434 | -0.321726 | -0.312407 | -0.321092 | -0.321181 |
| E3-E5 | -0.336630 | -0.336223 | -0.336655 | -0.336630 | -0.318837 | -0.339691 | -0.330781 |
| E3-E6 | -0.209919 | -0.209766 | -0.209123 | -0.209919 | -0.318559 | NA | NA |
| E4-E4 | -0.330443 | -0.330456 | -0.330223 | -0.330443 | -0.276521 | -0.344014 | -0.343952 |
| E4-E5 | -0.329350 | -0.329766 | -0.329986 | -0.329350 | -0.318837 | -0.321963 | -0.325686 |
| E4-E6 | -0.282536 | -0.282234 | -0.282766 | -0.282536 | -0.296451 | NA | NA |
| E5-E5 | -0.341230 | -0.341877 | -0.341543 | -0.341230 | -0.329569 | -0.348797 | -0.344116 |
| E5-E6 | -0.349490 | -0.349897 | -0.349212 | -0.349490 | -0.339062 | NA | NA |
| E6-E6 | -0.255994 | -0.255544 | -0.255433 | -0.255994 | -0.246247 | NA | NA |
| CTCF | -1.619170 | -1.619170 | -1.619170 | -1.619170 | -1.619170 | -1.619170 | -1.619170 |
| G1 | -0.056182 | -0.056655 | -0.056312 | -0.056182 | -0.056192 | -0.056182 | -0.056182 |
| G2 | -4.764573 | -4.764433 | -4.764322 | -4.764573 | -4.764578 | -4.764573 | -4.764573 |
| $K_b$ | 15.0 | 15.0 | 15.0 | 15.0 | 15.0 | 15.0 | 15.0 |
| $K_\theta$ | 1.0 | 1.0 | 1.0 | 1.0 | 1.0 | 1.0 | 1.0 |

**Table1: Optimal 26 parameters used to regenerate experimental Hi-C contact maps in cell line GM12878, IMR90, H1-hESC, mutated chromosome 8 (DEL), and various tissues (SC and EMM).**

**Contact Map Reconstruction**

To quantitatively compare simulated chromatin structures with experimental maps, we reconstructed contact probability maps from both Hi-C experiments and molecular dynamics trajectories using a consistent probabilistic framework. Experimental Hi-C maps measure the frequency of ligation events between genomic loci following crosslinking, digestion, proximity ligation, and sequencing. After read alignment and binning at a fixed genomic resolution, these interaction counts form a dense matrix representing pairwise contact frequencies between loci. To enable direct comparison with simulations, we convert the experimental matrix into a contact-probability map by normalizing each row to its maximum value, ensuring the nearest genomic neighbor has probability 1 and removing coverage bias across loci. This normalization produces a dimensionless matrix suitable for comparison with simulated contact probabilities and for extracting structural features such as compartments, TADs, and loops.

The simulated contact maps were constructed from ensembles of three-dimensional chromatin conformations generated by molecular dynamics simulations. For each configuration, the spatial distance between beads *i* and *j* was converted into a contact probability using a smooth crosslinking function as Eq. 8. This functional form corresponds to the probability of crosslinking between loci and is commonly used in chromatin modeling frameworks.

The ensemble-averaged simulated contact map is obtained by evaluating this function for all bead pairs across all frames and averaging over the molecular simulation trajectories,

$$P_{ij}^{\mathrm{sim}} = \frac{1}{N\mathrm{frames}} \sum_{n=1}^{\mathrm{frames}} 0.5[1 + tanh\zeta(r\mathrm{c} - r_{\mathrm{ij}}^{\mathrm{n}})]. \qquad (14)$$

We implement this procedure directly in the analysis by computing pairwise Euclidean distances for each configuration and converting them into probabilities at every simulation step using the hyperbolic-tangent switching function, then averaging over frames to generate the in silico Hi-C matrix.

Simulated and experimental contact maps were compared at identical genomic resolution, with simulation results typically shown in the lower triangle and experimental data in the upper triangle of the same matrix. This representation allows direct visualization of compartments, TAD boundaries, and loop enrichment patterns. This approach enables quantitative comparison between simulated chromatin and Hi-C experiments, forming the basis for parameter optimization and downstream structural analysis, including structural variations and tissue-specificity analysis.

**Observables**

A comprehensive set of observables from molecular dynamics trajectories was computed to characterize chromatin organization and structural conformations. The following observables will assist in enabling a direct connection between simulated chromatin conformations and the biological insight behind them. All quantities were calculated on the equilibrated ensemble and averaged across independent replicas to ensure statistical robustness.

*Chromatin State Interaction*

To quantify the effective interactions between chromatin states (E1-E6), we constructed a state-state interaction matrix directly from molecular dynamics trajectories. In contrast to contact-based measures in the Hi-C contact maps, this observable captures the underlying energetic interactions encoded in the energy function, including contributions from epigenetic interactions, long-range potentials, and loop-extrusion constraints. For each configuration, we obtain the pairwise interaction energy between loci i and j, $U_{ij}$, from the total potential energy decomposition. The effective interaction between chromatin states $\alpha$ and $\beta$ is then
defined as:

$$\mathcal{S}_{\alpha\beta} = \frac{1}{N_{\alpha\beta}} \Sigma_{i\in\alpha} \Sigma_{j\in\beta} \langle U_{ij} \rangle, \quad (15)$$

where $N_{\alpha\beta}$ is the number of beads pair belonging to states $\alpha$ and $\beta$. This observable provides a direct estimate of the effective free energy interaction between chromatin states, reflecting how epigenetic identity influences spatial organization through the full set of modeled potentials. To enable comparison across chromosomes and cell types, each interaction matrix is normalized using a min–max transformation. Diagonal elements quantify intra-state cohesion, while off-diagonal elements capture cross-state coupling.

*Chromatin Mobility and Diffusion*

Chromatin dynamics were computed and analyzed using the mean squared displacement (MSD):

$$MSD(\tau) = \left\langle |r_i(t+\tau) - r_i(t)|^2 \right\rangle_{i,t}. \quad (16)$$

The MSD was computed using a fast Fourier transform-based implementation to ensure numerical efficiency over long trajectories. For long timescales, the MSD demonstrates a power-law behavior:

$$MSD(\tau) \sim \tau^{\nu}, \quad (17)$$

where $\nu$ characterizes diffusion. Separate analyses were performed for loci belonging to euchromatin and heterochromatin compartments, enabling compartment-specific characterization of chromatin mobility.

*Tail Amplification of Contact Distributions*

We computed the heterogeneity of chromatin contacts between cell lines using the tail amplification metric, which probes the upper tail of the contact probability distribution. Let $P_{ij}$ denote the ensemble of contact probabilities evaluated over all locus pairs and sampled configurations. We define $Q_{99}$ as the 99th percentile of this distribution, given by:

$$P(P_{ij} \leq Q_{99}) = 0.99, \quad (18)$$

such that 99% of contact probabilities lie below $Q_{99}$, while the remaining 1% correspond to the

strongest interactions. The tail amplification ratio is then defined as:

$$T = \frac{Q_{99}}{\mathrm{P}_{\mathrm{ij}}}, \tag{19}$$

where $\mathrm{P}_{\mathrm{ij}}$ denotes the mean contact probability over the ensemble. This observable captures the contribution of rare, high-probability contacts and therefore provides a sensitive measure of structural heterogeneity within the chromatin ensemble across cell lines.

*Long-Lived Contact Fraction*

To characterize the temporal persistence of chromatin contacts, we define a binary contact variable:

$$C_{\mathrm{ij}} = \Theta(r_{\mathrm{c}} - r_{\mathrm{ij}}(t)), \tag{20}$$

where $\Theta$ is the Heaviside step function. The temporal stability of contact is quantified through the autocorrelation function:

$$A_{\mathrm{ij}} = \langle C_{\mathrm{ij}}(t) C_{\mathrm{ij}}(t+\tau) \rangle. \tag{21}$$

Where the long-lived contact fraction is then defined as:

$$f_{\mathrm{LL}} = \frac{\Sigma_{\mathrm{ij}} A_{\mathrm{ij}}(\tau > \tau_0)}{\Sigma_{\mathrm{ij}} \langle C_{\mathrm{ij}} \rangle}, \tag{22}$$

where $\tau_0$ is a threshold timescale used to isolate persistent interactions. This metric provides a measure of structural memory within chromatin conformations.

*Compartmentalization Strength*

The strength of euchromatin (EU)-heterochromatin (HE) compartmentalization is quantified using:

$$\Psi = \frac{P_{\text{EU-EU}} + P_{\text{HE-HE}}}{2P_{\text{EU-HE}}}, \tag{23}$$

where $P_{\text{EU-EU}}$, $P_{\text{HE-HE}}$, $P_{\text{EU-HE}}$ denote the average contact probabilities within euchromatin, within heterochromatin, and between euchromatin and heterochromatin compartments, respectively. Larger values of $\Psi$ indicate stronger segregation between euchromatin and heterochromatin domains.

*Radius of Gyration and End-to-End Distance*

To assess the chromosome's overall compactness, we quantify its size using the radius of gyration and End-to-End distance. The radius of gyration is quantified by:

$$R_g = \left(\frac{1}{N}\Sigma_{i=1}^{N}|\vec{r_i} - \vec{r_{cm}}|^2\right)^{(1/2)}, \quad (24)$$

where $r_{cm}$ is the center-of-mass position. The large-scale extension is characterized by the end-to-end distance:

$$R_{ee} = |\vec{r}_N - \vec{r}_1|. \quad (25)$$

*State Population and Lifetime*

The structural heterogeneity in the regions affected by structural variants has been quantified by the population and lifetime of conformational states identified from molecular dynamics trajectories. The population os state $\alpha$ is defined as:

$$p_\alpha = \frac{N_\alpha}{N_{total}}, \quad (26)$$

where $N_\alpha$ is the number of configurations assigned to the state $\alpha$ criterion. The lifetime of each state is computed as the mean residence time:

$$\tau_\alpha = \langle t_{res}^{(\alpha)} \rangle. \quad (27)$$

These quantities provide a kinetic description of structural transitions induced by genomic rearrangement.

*Tissue Specific Order Parameters*

To characterize chromatin architecture within tissue-specific regions of interest, we defined two order parameters based on local contact statistics. The compartment segregation order parameter is defined as:

$$q_\phi = \frac{N_{intra} - N_{inter}}{N_{intra} + N_{inter}}, \quad (28)$$

### Supplementary Figures

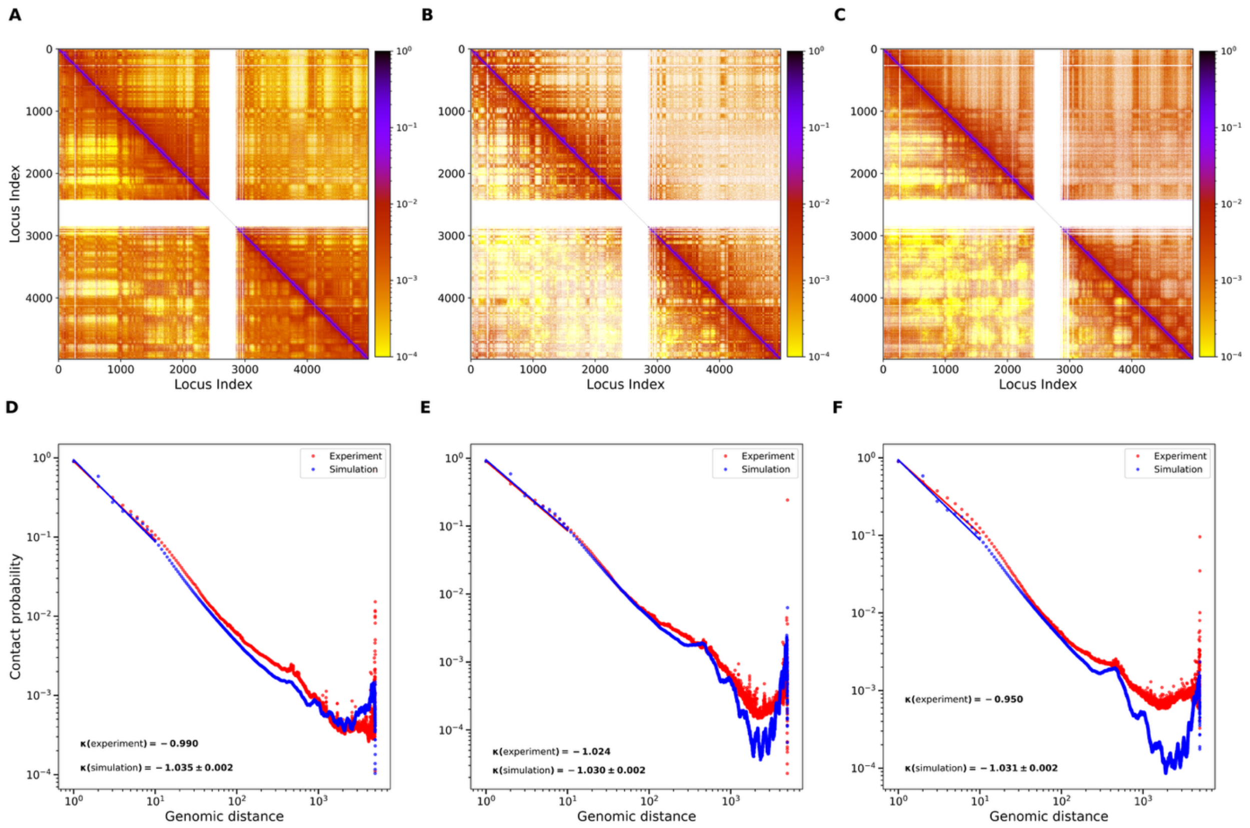


Fig S1. **Validation of chromosome 1 chromatin architecture simulated by the polymer model across three human cell types. (A–C)** Experimental Hi-C contact maps (upper triangles) are compared with polymer simulation contact maps (lower triangles) for chromosome 1 in **GM12878** (A), **IMR90** (B), and **H1-hESC** (C). Contact frequencies are displayed on a logarithmic color scale. The close visual agreement between experimental and simulated maps demonstrates that the model accurately reproduces chromosome-scale interaction patterns across diverse chromatin contexts. **(D–F)** Contact probability as a function of genomic distance, $P(s)$, derived from experimental Hi-C data (red) and polymer simulations (blue) for **GM12878** (D), **IMR90** (E), and **H1-hESC** (F). The fitted power-law scaling exponent ($\kappa$) is shown for both experiment and simulation in each panel, demonstrating that the simulations faithfully recapitulate the distance-dependent decay of chromatin contacts observed experimentally.

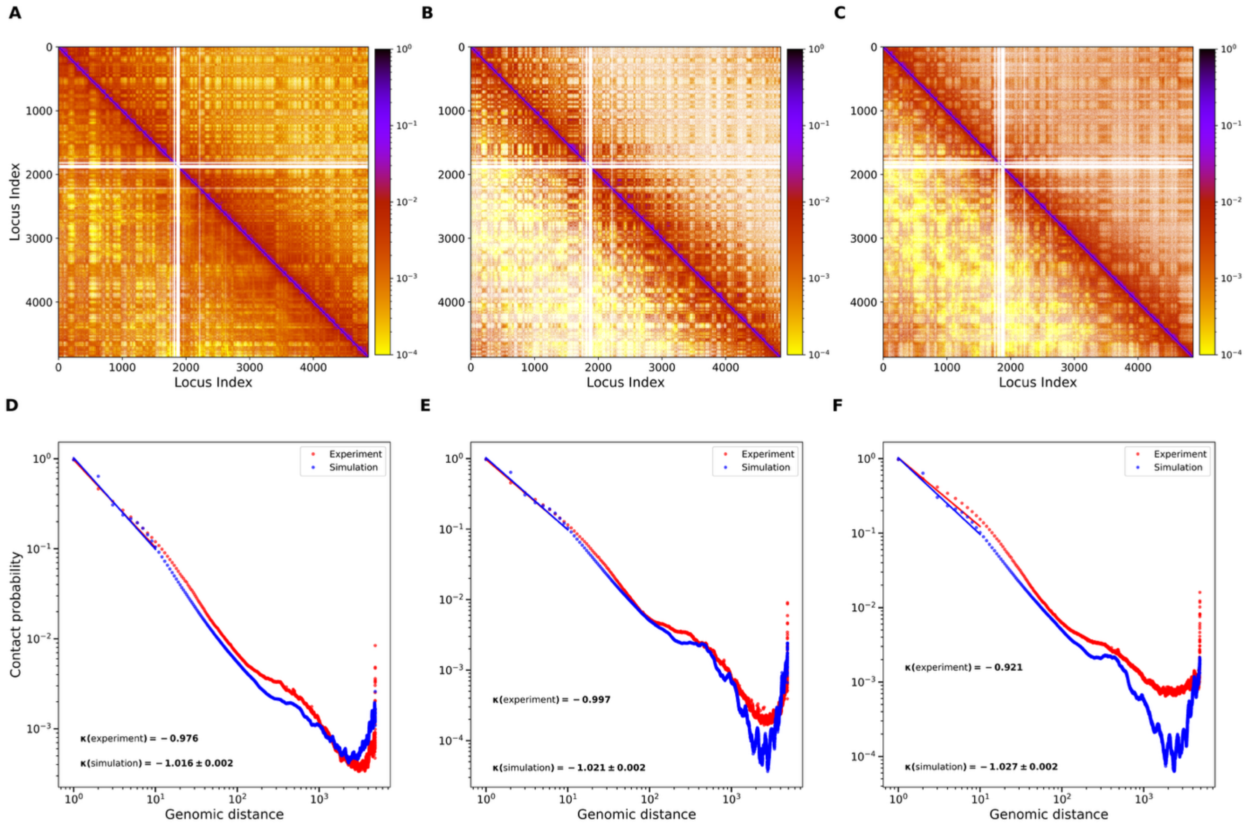


**Fig S2. Validation of chromosome 2 chromatin architecture simulated by the polymer model across three human cell types. (A–C)** Experimental Hi-C contact maps (upper triangles) are compared with polymer simulation contact maps (lower triangles) for chromosome 2 in **GM12878** (A), **IMR90** (B), and **H1-hESC** (C). Contact frequencies are displayed on a logarithmic color scale. The close visual agreement between experimental and simulated maps demonstrates that the model accurately reproduces chromosome-scale interaction patterns across diverse chromatin contexts. **(D–F)** Contact probability as a function of genomic distance, $P(s)$, derived from experimental Hi-C data (red) and polymer simulations (blue) for **GM12878** (D), **IMR90** (E), and **H1-hESC** (F). The fitted power-law scaling exponent ($\kappa$) is shown for both experiment and simulation in each panel, demonstrating that the simulations faithfully recapitulate the distance-dependent decay of chromatin contacts observed experimentally.

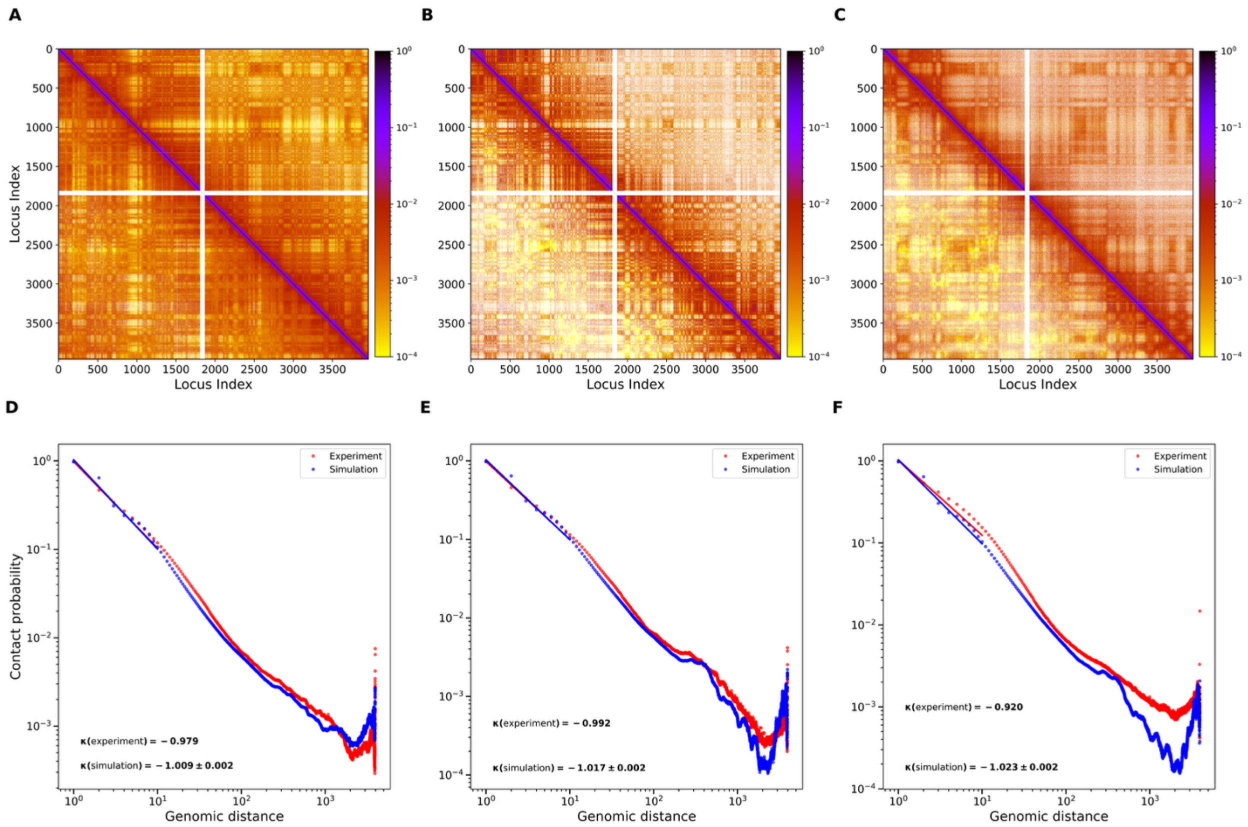


**Fig S3. Validation of chromosome 3 chromatin architecture simulated by the polymer model across three human cell types. (A–C)** Experimental Hi-C contact maps (upper triangles) are compared with polymer simulation contact maps (lower triangles) for chromosome 3 in **GM12878** (A), **IMR90** (B), and **H1-hESC** (C). Contact frequencies are displayed on a logarithmic color scale. The close visual agreement between experimental and simulated maps demonstrates that the model accurately reproduces chromosome-scale interaction patterns across diverse chromatin contexts. **(D–F)** Contact probability as a function of genomic distance, $P(s)$, derived from experimental Hi-C data (red) and polymer simulations (blue) for **GM12878** (D), **IMR90** (E), and **H1-hESC** (F). Each panel shows the fitted power-law scaling exponent ($\kappa$) for both experiment and simulation, demonstrating that the simulations faithfully recapitulate the distance-dependent decay of chromatin contacts observed experimentally.

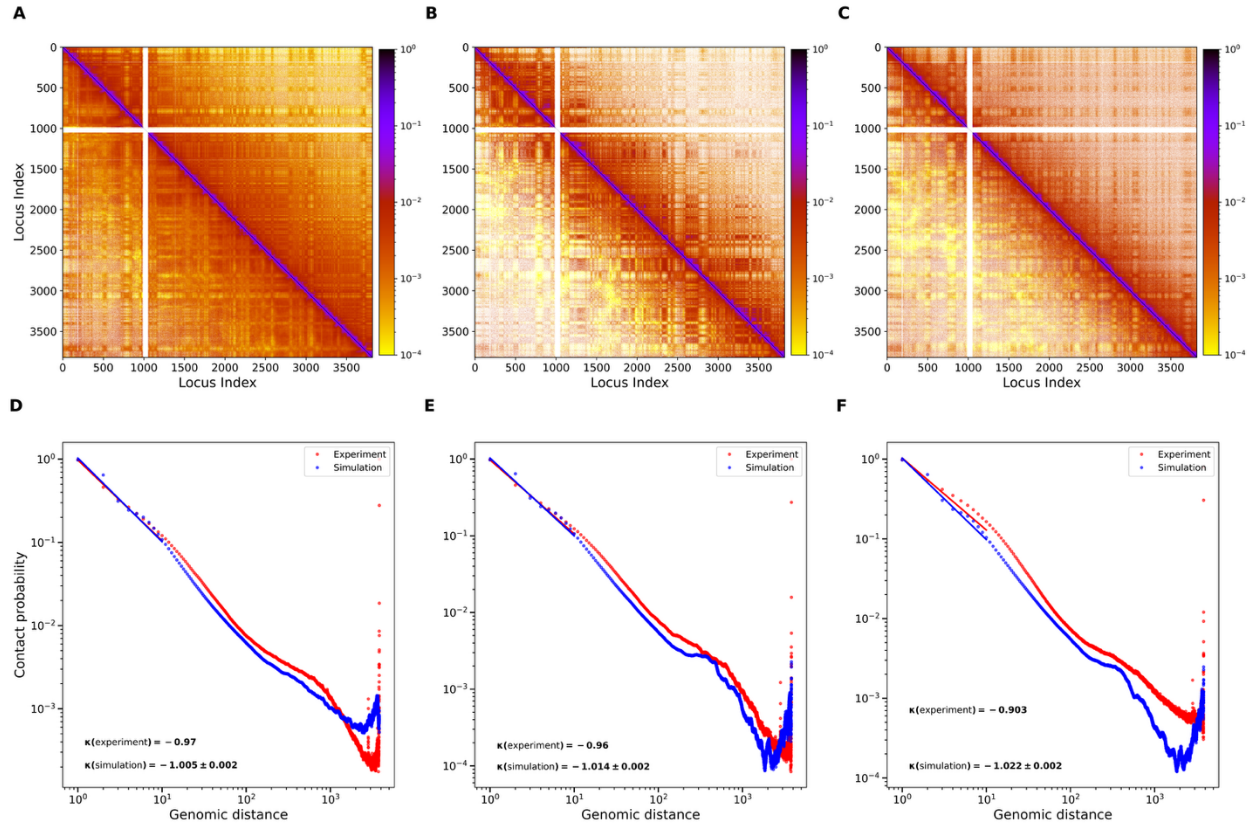


**Fig S4. Validation of chromosome 4 chromatin architecture simulated by the polymer model across three human cell types. (A–C)** Experimental Hi-C contact maps (upper triangles) are compared with polymer simulation contact maps (lower triangles) for chromosome 4 in **GM12878** (A), **IMR90** (B), and **H1-hESC** (C). Contact frequencies are displayed on a logarithmic color scale. The close visual agreement between experimental and simulated maps demonstrates that the model accurately reproduces chromosome-scale interaction patterns across diverse chromatin contexts. **(D–F)** Contact probability as a function of genomic distance, $P(s)$, derived from experimental Hi-C data (red) and polymer simulations (blue) for **GM12878** (D), **IMR90** (E), and **H1-hESC** (F). The fitted power-law scaling exponent ($\kappa$) is shown for both experiment and simulation in each panel, demonstrating that the simulations faithfully recapitulate the distance-dependent decay of chromatin contacts observed experimentally.

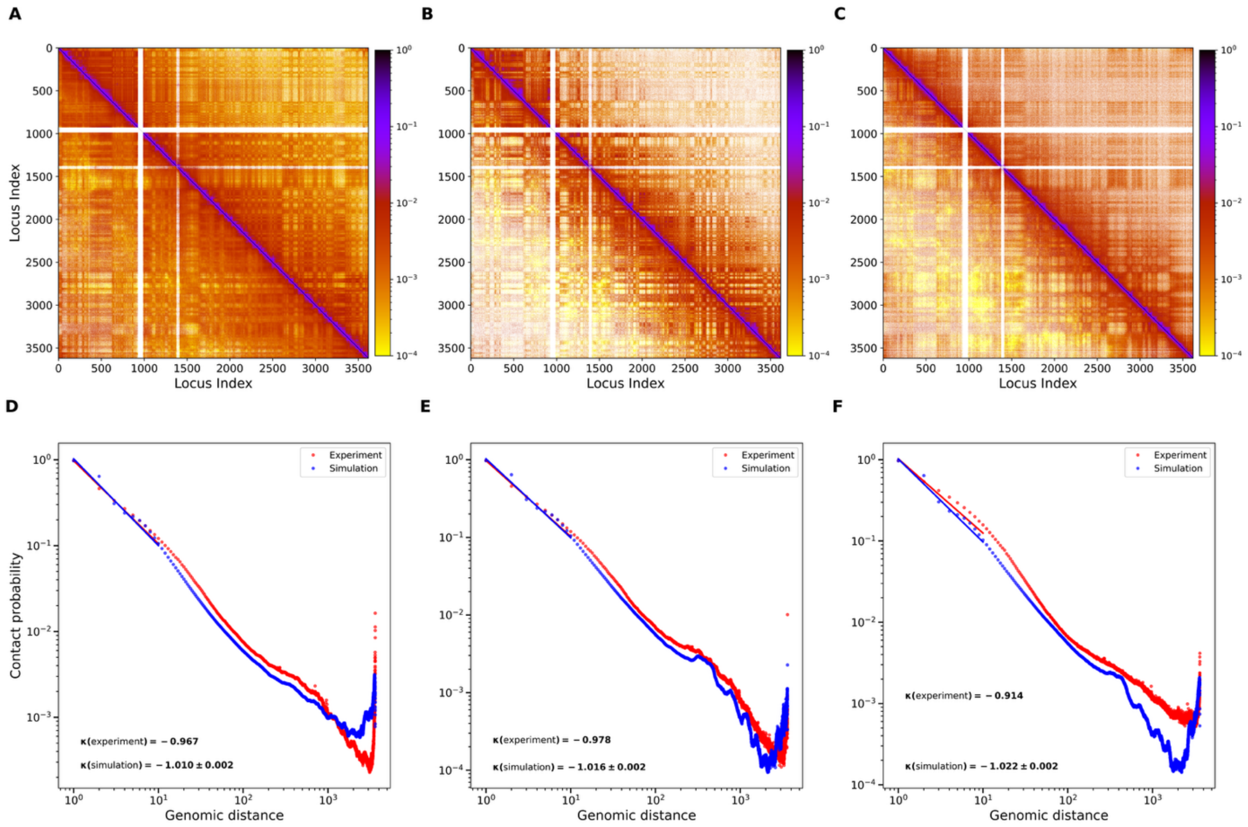


**Fig S5. Validation of chromosome 5 chromatin architecture simulated by the polymer model across three human cell types. (A–C)** Experimental Hi-C contact maps (upper triangles) are compared with polymer simulation contact maps (lower triangles) for chromosome 5 in **GM12878** (A), **IMR90** (B), and **H1-hESC** (C). Contact frequencies are displayed on a logarithmic color scale. The close visual agreement between experimental and simulated maps demonstrates that the model accurately reproduces chromosome-scale interaction patterns across diverse chromatin contexts. **(D–F)** Contact probability as a function of genomic distance, $P(s)$, derived from experimental Hi-C data (red) and polymer simulations (blue) for **GM12878** (D), **IMR90** (E), and **H1-hESC** (F). The fitted power-law scaling exponent ($\kappa$) is shown for both experiment and simulation in each panel, demonstrating that the simulations faithfully recapitulate the distance-dependent decay of chromatin contacts observed experimentally.

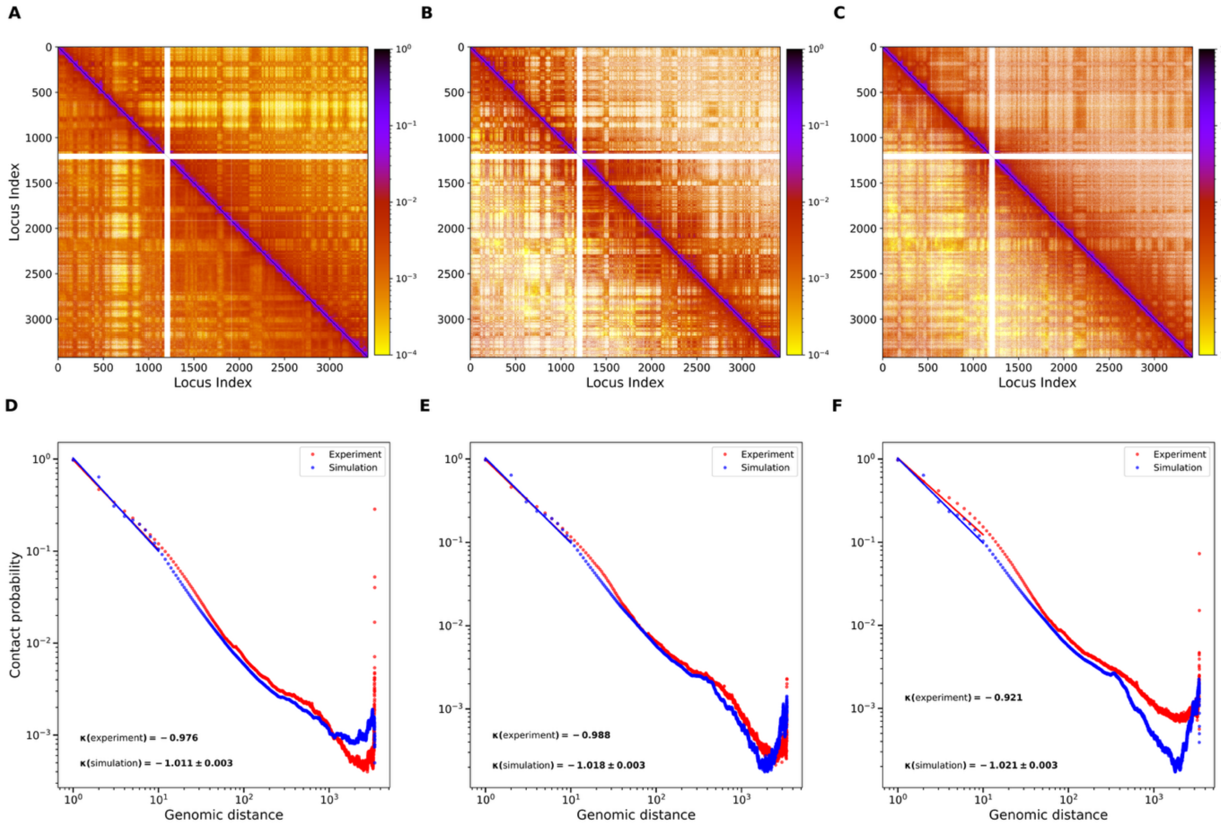


**Fig S6. Validation of chromosome 6 chromatin architecture simulated by the polymer model across three human cell types. (A–C)** Experimental Hi-C contact maps (upper triangles) are compared with polymer simulation contact maps (lower triangles) for chromosome 6 in **GM12878** (A), **IMR90** (B), and **H1-hESC** (C). Contact frequencies are displayed on a logarithmic color scale. The close visual agreement between experimental and simulated maps demonstrates that the model accurately reproduces chromosome-scale interaction patterns across diverse chromatin contexts. **(D–F)** Contact probability as a function of genomic distance, $P(s)$, derived from experimental Hi-C data (red) and polymer simulations (blue) for **GM12878** (D), **IMR90** (E), and **H1-hESC** (F). The fitted power-law scaling exponent ($\kappa$) is shown for both experiment and simulation in each panel, demonstrating that the simulations faithfully recapitulate the distance-dependent decay of chromatin contacts observed experimentally.

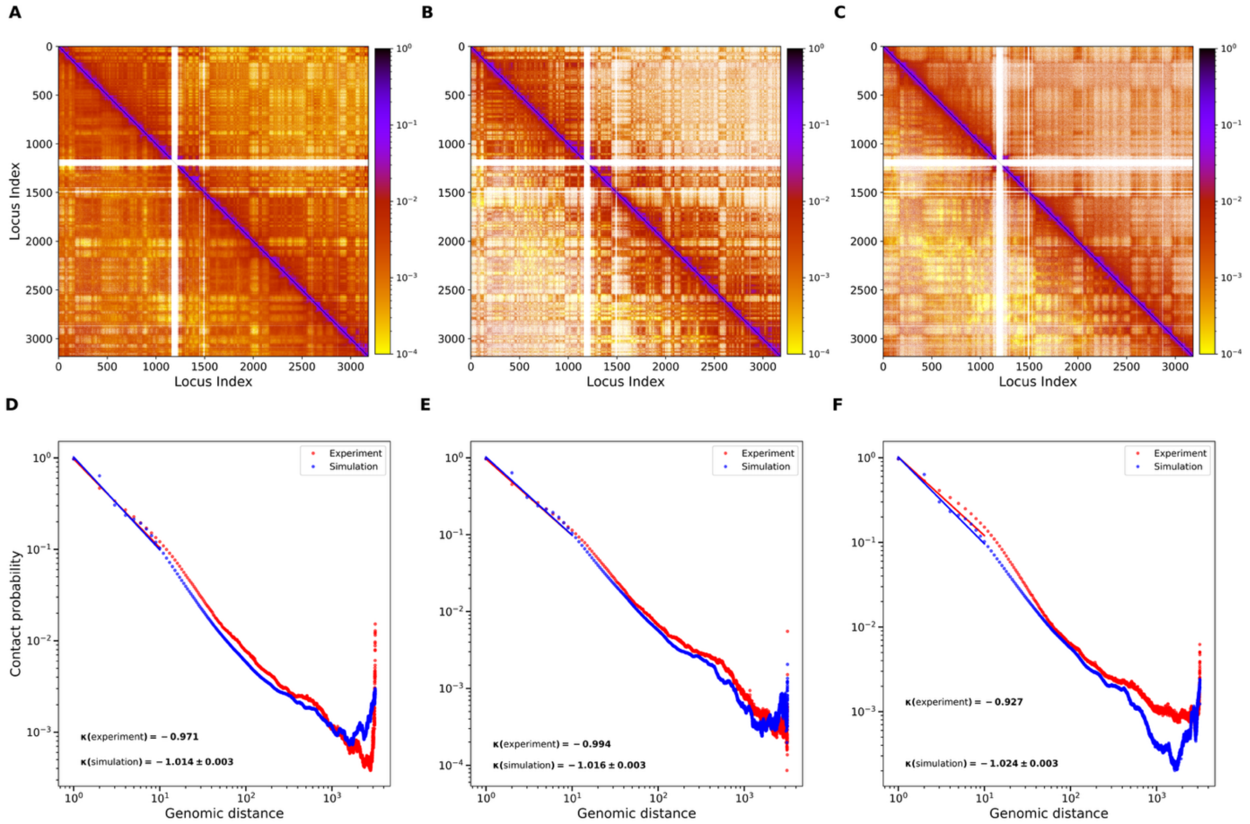


**Fig S7. Validation of chromosome 7 chromatin architecture simulated by the polymer model across three human cell types. (A–C)** Experimental Hi-C contact maps (upper triangles) are compared with polymer simulation contact maps (lower triangles) for chromosome 7 in **GM12878** (A), **IMR90** (B), and **H1-hESC** (C). Contact frequencies are displayed on a logarithmic color scale. The close visual agreement between experimental and simulated maps demonstrates that the model accurately reproduces chromosome-scale interaction patterns across diverse chromatin contexts. **(D–F)** Contact probability as a function of genomic distance, $P(s)$, derived from experimental Hi-C data (red) and polymer simulations (blue) for **GM12878** (D), **IMR90** (E), and **H1-hESC** (F). The fitted power-law scaling exponent ($\kappa$) is shown for both experiment and simulation in each panel, demonstrating that the simulations faithfully recapitulate the distance-dependent decay of chromatin contacts observed experimentally.

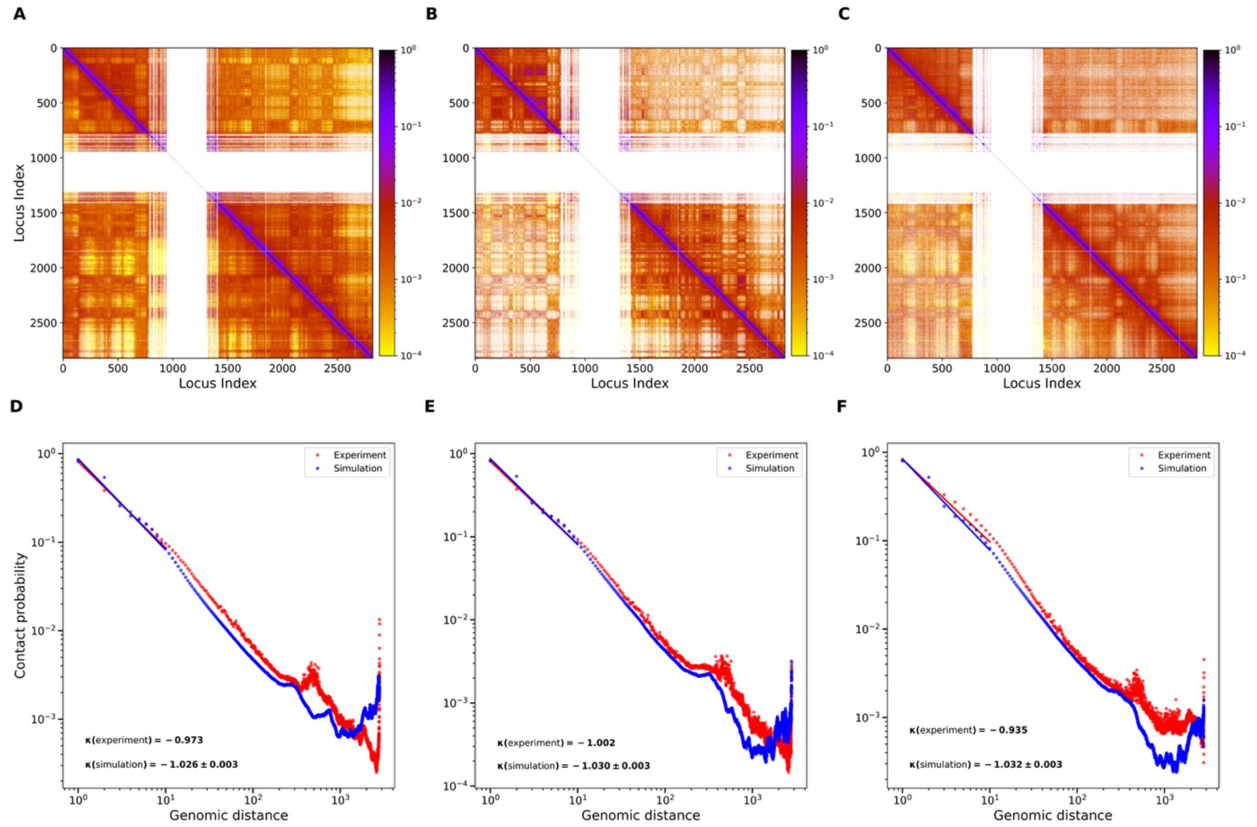


**Fig S8. Validation of chromosome 9 chromatin architecture simulated by the polymer model across three human cell types. (A–C)** Experimental Hi-C contact maps (upper triangles) are compared with polymer simulation contact maps (lower triangles) for chromosome 9 in **GM12878** (A), **IMR90** (B), and **H1-hESC** (C). Contact frequencies are displayed on a logarithmic color scale. The close visual agreement between experimental and simulated maps demonstrates that the model accurately reproduces chromosome-scale interaction patterns across diverse chromatin contexts. **(D–F)** Contact probability as a function of genomic distance, $P(s)$, derived from experimental Hi-C data (red) and polymer simulations (blue) for **GM12878** (D), **IMR90** (E), and **H1-hESC** (F). Each panel shows the fitted power-law scaling exponent ($\kappa$) for both experiment and simulation, demonstrating that the simulations faithfully recapitulate the distance-dependent decay of chromatin contacts observed experimentally.

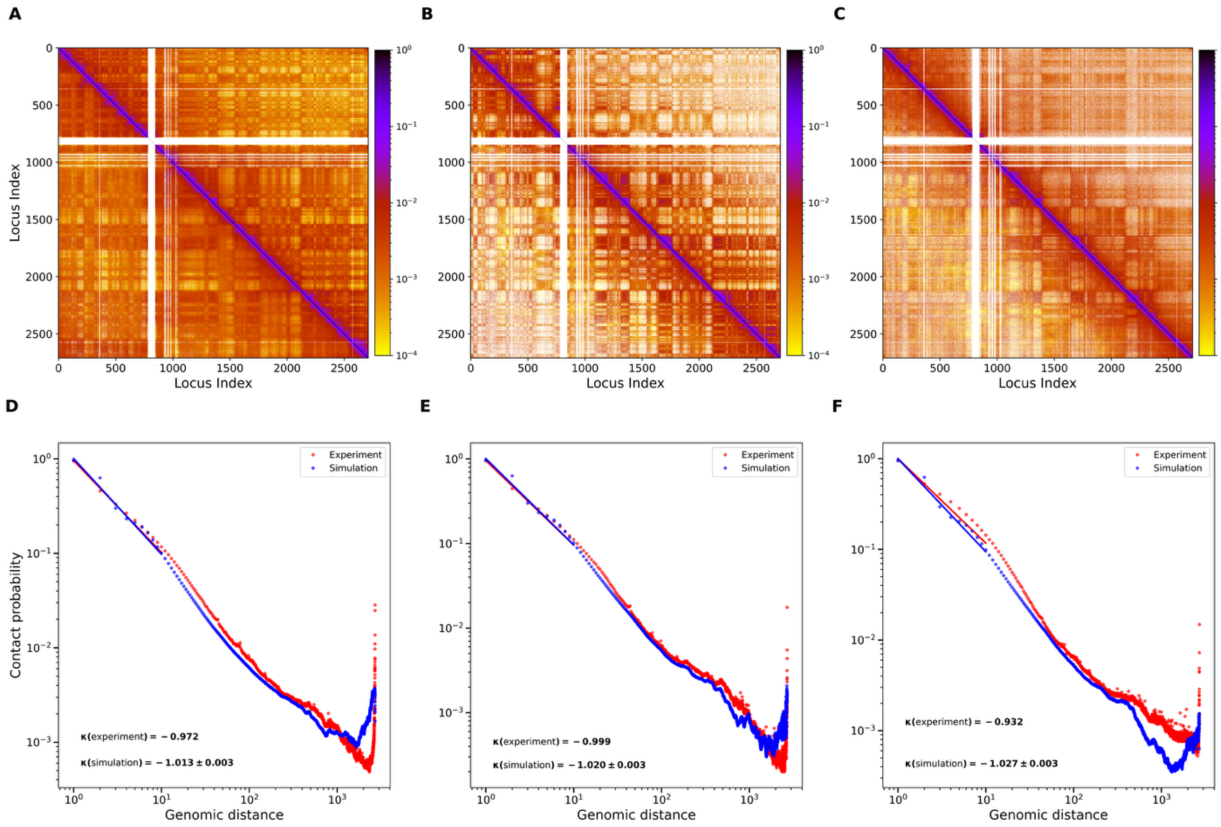


**Fig S9. Validation of chromosome 10 chromatin architecture simulated by the polymer model across three human cell types. (A–C)** Experimental Hi-C contact maps (upper triangles) are compared with polymer simulation contact maps (lower triangles) for chromosome 10 in **GM12878** (A), **IMR90** (B), and **H1-hESC** (C). Contact frequencies are displayed on a logarithmic color scale. The close visual agreement between experimental and simulated maps demonstrates that the model accurately reproduces chromosome-scale interaction patterns across diverse chromatin contexts. **(D–F)** Contact probability as a function of genomic distance, $P(s)$, derived from experimental Hi-C data (red) and polymer simulations (blue) for **GM12878** (D), **IMR90** (E), and **H1-hESC** (F). The fitted power-law scaling exponent ($\kappa$) is shown for both experiment and simulation in each panel, demonstrating that the simulations faithfully recapitulate the distance-dependent decay of chromatin contacts observed experimentally.

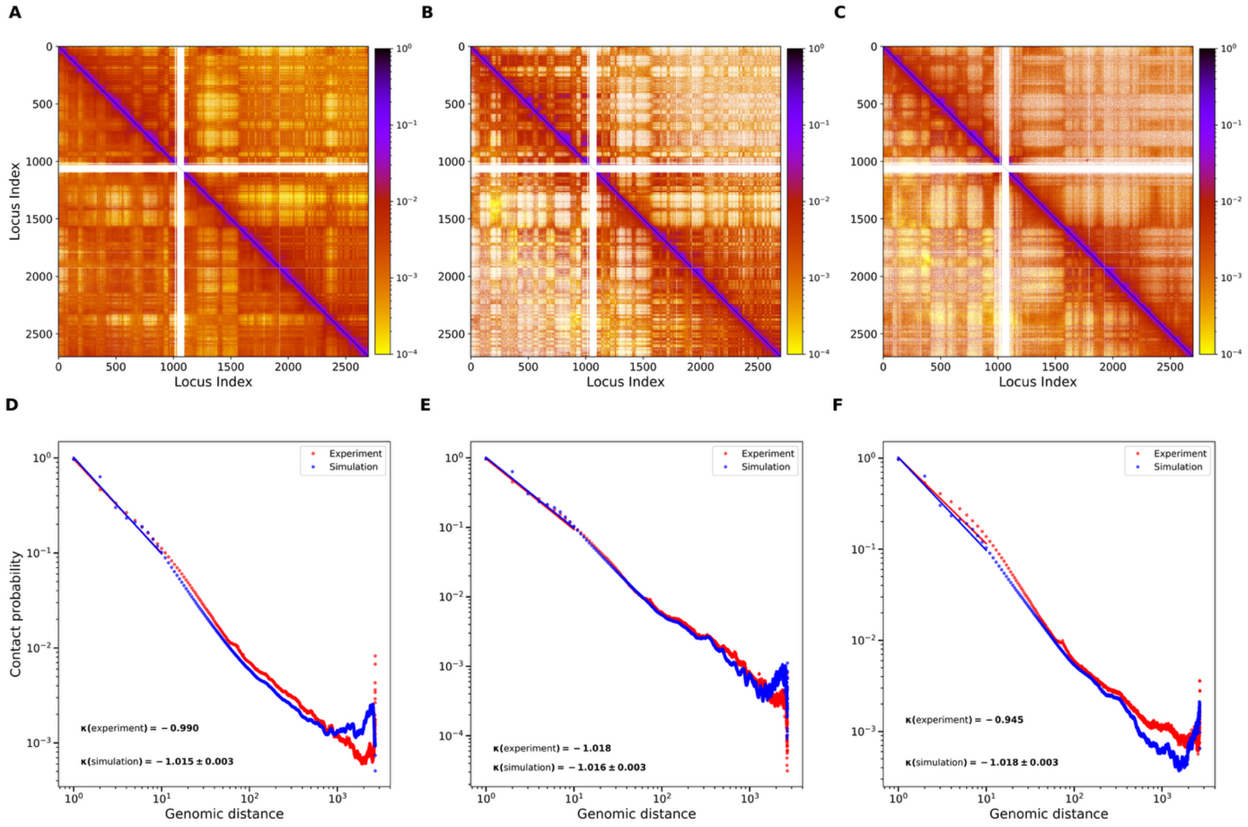


**Fig S10. Validation of chromosome 11 chromatin architecture simulated by the polymer model across three human cell types. (A–C)** Experimental Hi-C contact maps (upper triangles) are compared with polymer simulation contact maps (lower triangles) for chromosome 11 in **GM12878** (A), **IMR90** (B), and **H1-hESC** (C). Contact frequencies are displayed on a logarithmic color scale. The close visual agreement between experimental and simulated maps demonstrates that the model accurately reproduces chromosome-scale interaction patterns across diverse chromatin contexts. **(D–F)** Contact probability as a function of genomic distance, $P(s)$, derived from experimental Hi-C data (red) and polymer simulations (blue) for **GM12878** (D), **IMR90** (E), and **H1-hESC** (F). The fitted power-law scaling exponent ($\kappa$) is shown for both experiment and simulation in each panel, demonstrating that the simulations faithfully recapitulate the distance-dependent decay of chromatin contacts observed experimentally.

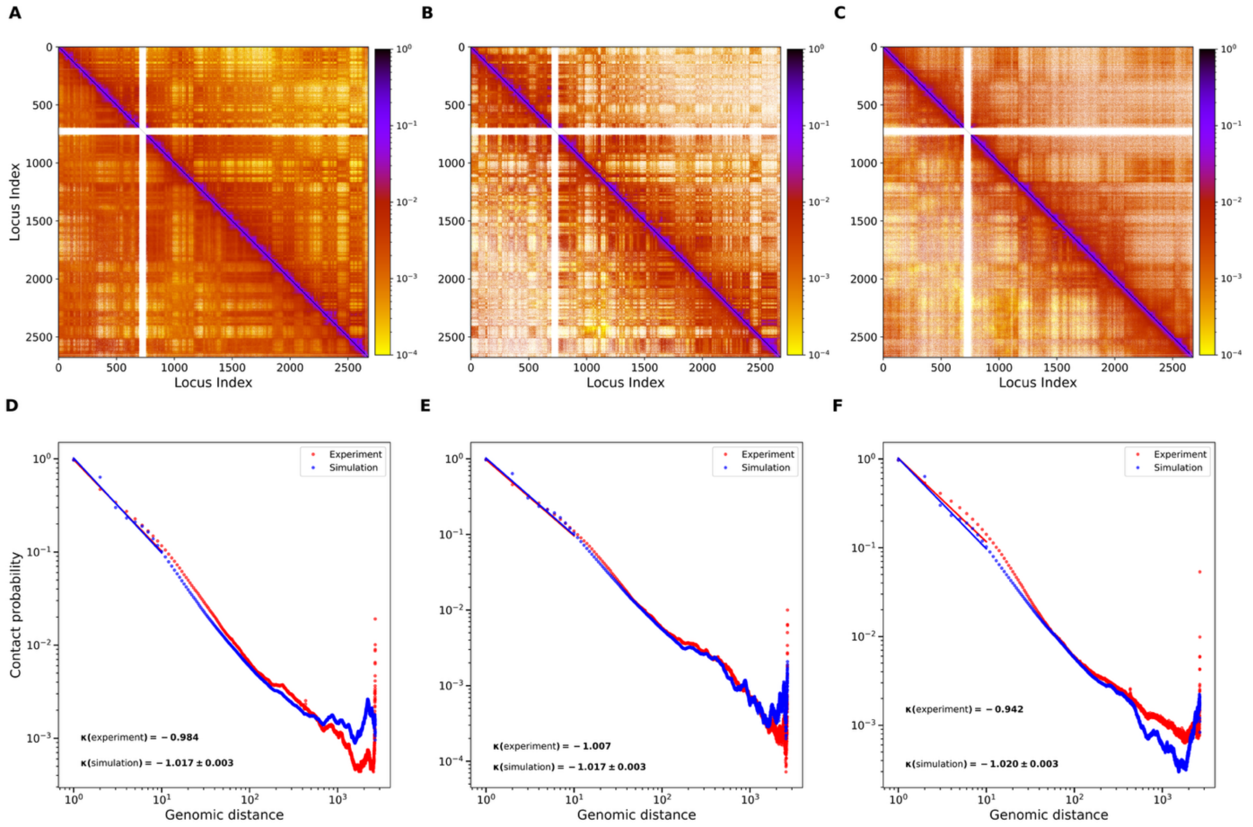


**Fig S11. Validation of chromosome 12 chromatin architecture simulated by the polymer model across three human cell types. (A–C)** Experimental Hi-C contact maps (upper triangles) are compared with polymer simulation contact maps (lower triangles) for chromosome 12 in **GM12878** (A), **IMR90** (B), and **H1-hESC** (C). Contact frequencies are displayed on a logarithmic color scale. The close visual agreement between experimental and simulated maps demonstrates that the model accurately reproduces chromosome-scale interaction patterns across diverse chromatin contexts. **(D–F)** Contact probability as a function of genomic distance, $P(s)$, derived from experimental Hi-C data (red) and polymer simulations (blue) for **GM12878** (D), **IMR90** (E), and **H1-hESC** (F). The fitted power-law scaling exponent ($\kappa$) is shown for both experiment and simulation in each panel, demonstrating that the simulations faithfully recapitulate the distance-dependent decay of chromatin contacts observed experimentally.

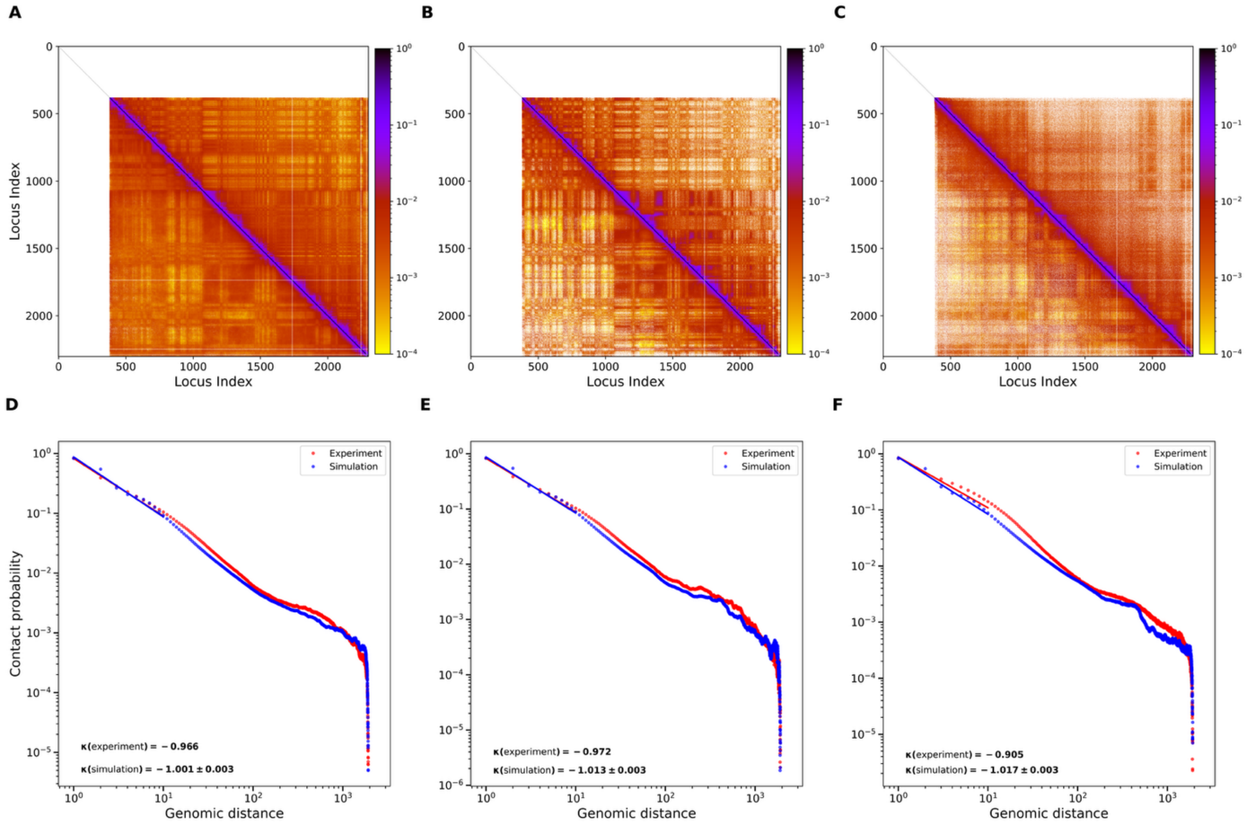


**Fig S12. Validation of chromosome 13 chromatin architecture simulated by the polymer model across three human cell types. (A–C)** Experimental Hi-C contact maps (upper triangles) are compared with polymer simulation contact maps (lower triangles) for chromosome 13 in **GM12878** (A), **IMR90** (B), and **H1-hESC** (C). Contact frequencies are displayed on a logarithmic color scale. The close visual agreement between experimental and simulated maps demonstrates that the model accurately reproduces chromosome-scale interaction patterns across diverse chromatin contexts. **(D–F)** Contact probability as a function of genomic distance, $P(s)$, derived from experimental Hi-C data (red) and polymer simulations (blue) for **GM12878** (D), **IMR90** (E), and **H1-hESC** (F). The fitted power-law scaling exponent ($\kappa$) is shown for both experiment and simulation in each panel, demonstrating that the simulations faithfully recapitulate the distance-dependent decay of chromatin contacts observed experimentally.

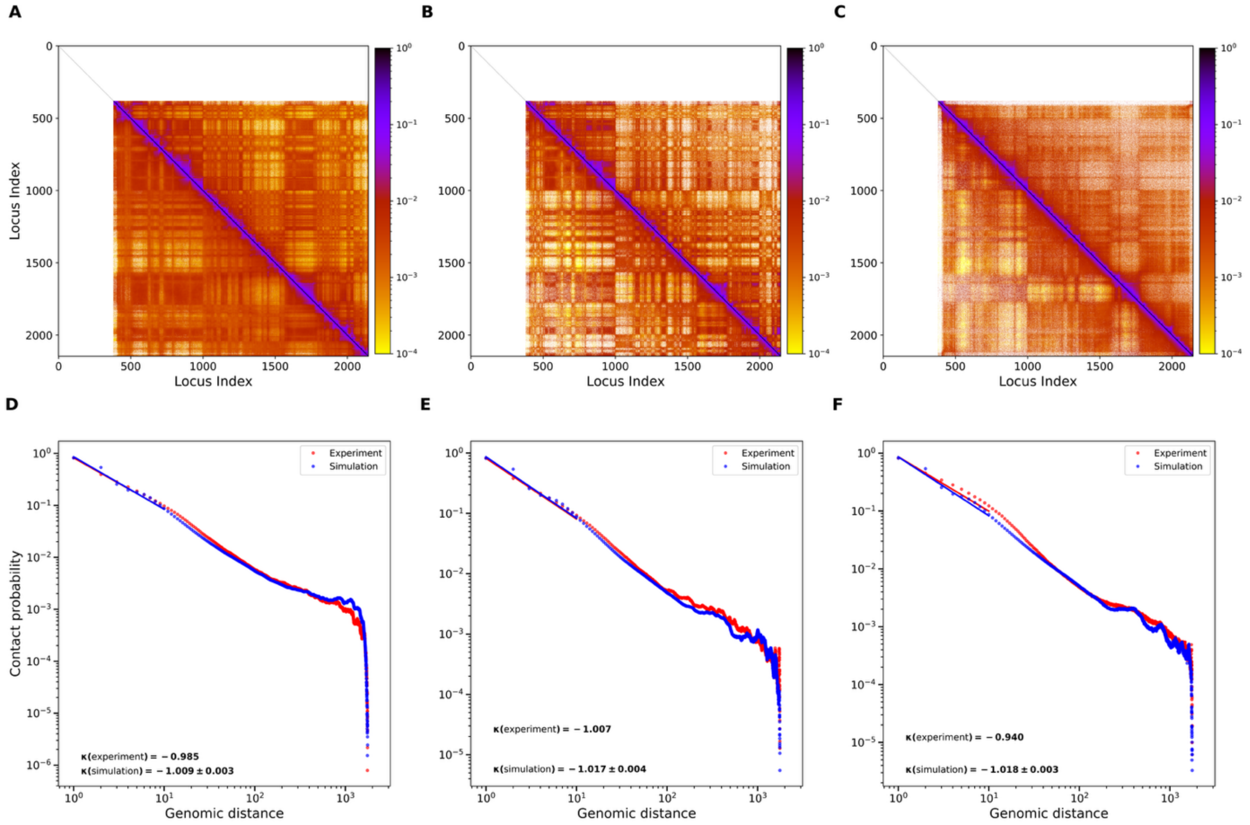


**Fig S13. Validation of chromosome 14 chromatin architecture simulated by the polymer model across three human cell types. (A–C)** Experimental Hi-C contact maps (upper triangles) are compared with polymer simulation contact maps (lower triangles) for chromosome 14 in **GM12878** (A), **IMR90** (B), and **H1-hESC** (C). Contact frequencies are displayed on a logarithmic color scale. The close visual agreement between experimental and simulated maps demonstrates that the model accurately reproduces chromosome-scale interaction patterns across diverse chromatin contexts. **(D–F)** Contact probability as a function of genomic distance, $P(s)$, derived from experimental Hi-C data (red) and polymer simulations (blue) for **GM12878** (D), **IMR90** (E), and **H1-hESC** (F). The fitted power-law scaling exponent ($\kappa$) is shown for both experiment and simulation in each panel, demonstrating that the simulations faithfully recapitulate the distance-dependent decay of chromatin contacts observed experimentally.

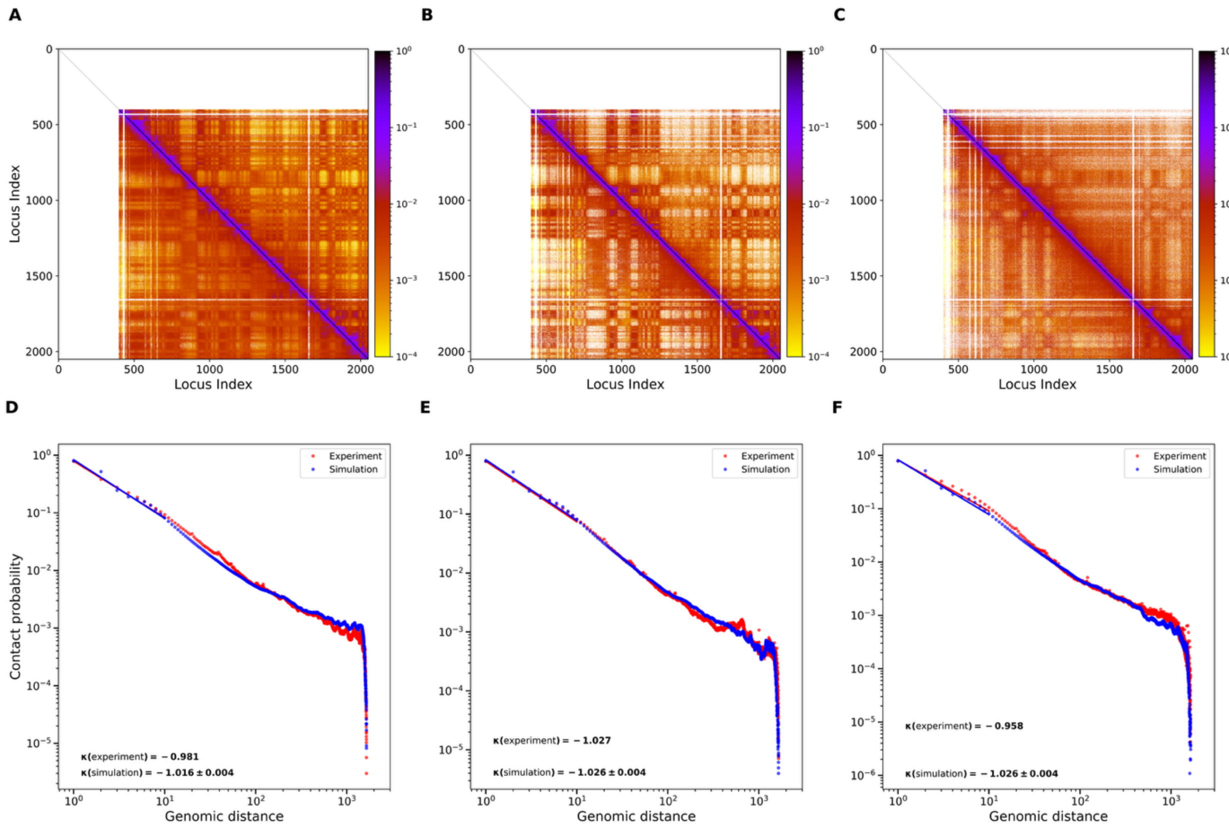


**Fig S14. Validation of chromosome 15 chromatin architecture simulated by the polymer model across three human cell types. (A–C)** Experimental Hi-C contact maps (upper triangles) are compared with polymer simulation contact maps (lower triangles) for chromosome 15 in **GM12878** (A), **IMR90** (B), and **H1-hESC** (C). Contact frequencies are displayed on a logarithmic color scale. The close visual agreement between experimental and simulated maps demonstrates that the model accurately reproduces chromosome-scale interaction patterns across diverse chromatin contexts. **(D–F)** Contact probability as a function of genomic distance, $P(s)$, derived from experimental Hi-C data (red) and polymer simulations (blue) for **GM12878** (D), **IMR90** (E), and **H1-hESC** (F). The fitted power-law scaling exponent ($\kappa$) is shown for both experiment and simulation in each panel, demonstrating that the simulations faithfully recapitulate the distance-dependent decay of chromatin contacts observed experimentally.

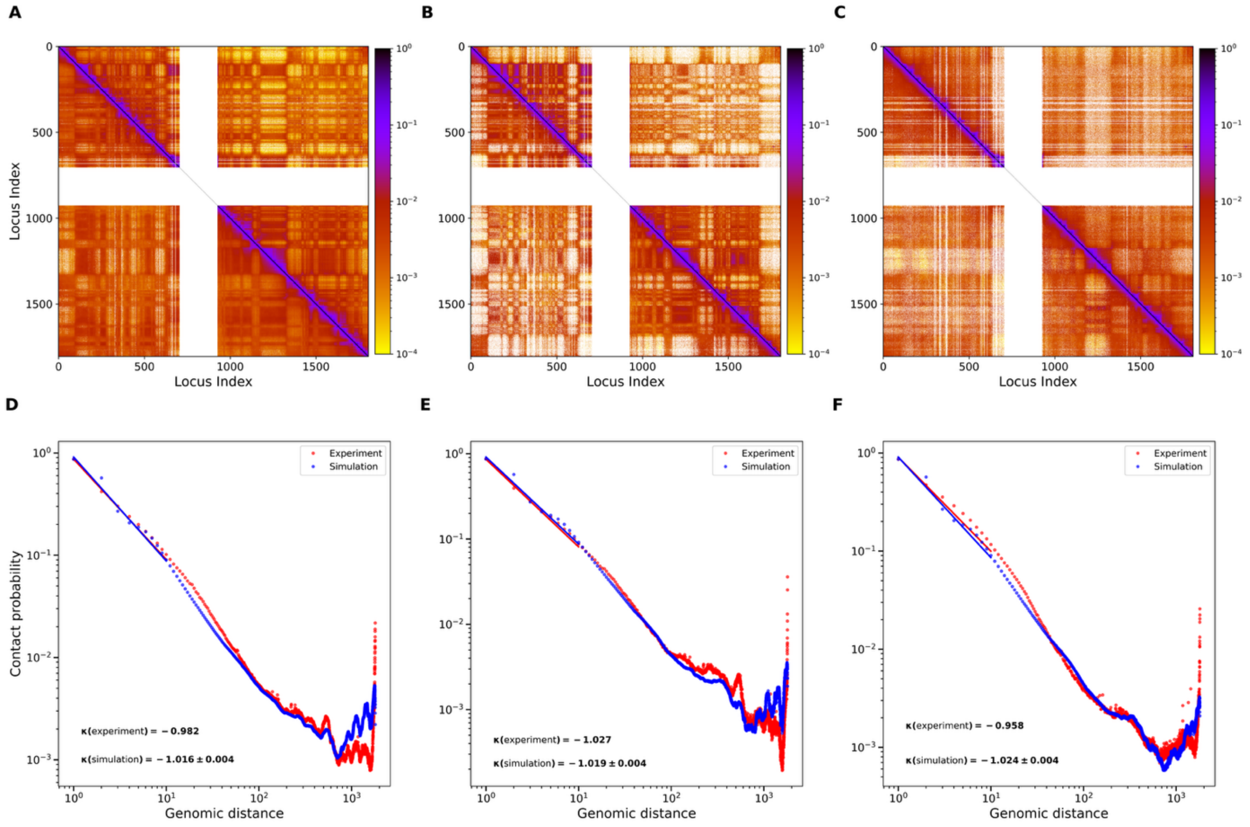


**Fig S15. Validation of chromosome 16 chromatin architecture simulated by the polymer model across three human cell types. (A–C)** Experimental Hi-C contact maps (upper triangles) are compared with polymer simulation contact maps (lower triangles) for chromosome 16 in **GM12878** (A), **IMR90** (B), and **H1-hESC** (C). Contact frequencies are displayed on a logarithmic color scale. The close visual agreement between experimental and simulated maps demonstrates that the model accurately reproduces chromosome-scale interaction patterns across diverse chromatin contexts. **(D–F)** Contact probability as a function of genomic distance, $P(s)$, derived from experimental Hi-C data (red) and polymer simulations (blue) for **GM12878** (D), **IMR90** (E), and **H1-hESC** (F). The fitted power-law scaling exponent ($\kappa$) is shown for both experiment and simulation in each panel, demonstrating that the simulations faithfully recapitulate the distance-dependent decay of chromatin contacts observed experimentally.

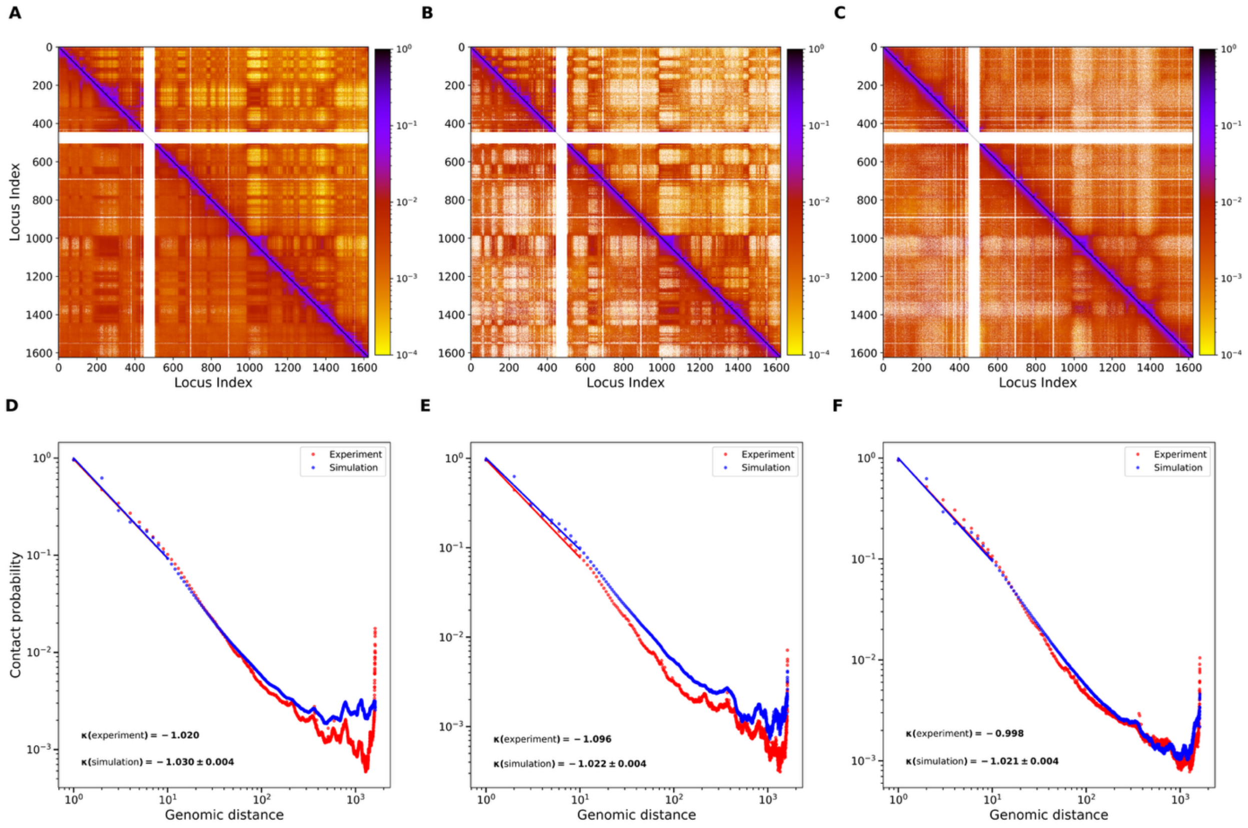


**Fig S16. Validation of chromosome 17 chromatin architecture simulated by the polymer model across three human cell types. (A–C)** Experimental Hi-C contact maps (upper triangles) are compared with polymer simulation contact maps (lower triangles) for chromosome 17 in **GM12878** (A), **IMR90** (B), and **H1-hESC** (C). Contact frequencies are displayed on a logarithmic color scale. The close visual agreement between experimental and simulated maps demonstrates that the model accurately reproduces chromosome-scale interaction patterns across diverse chromatin contexts. **(D–F)** Contact probability as a function of genomic distance, $P(s)$, derived from experimental Hi-C data (red) and polymer simulations (blue) for **GM12878** (D), **IMR90** (E), and **H1-hESC** (F). The fitted power-law scaling exponent ($\kappa$) is shown for both experiment and simulation in each panel, demonstrating that the simulations faithfully recapitulate the distance-dependent decay of chromatin contacts observed experimentally.

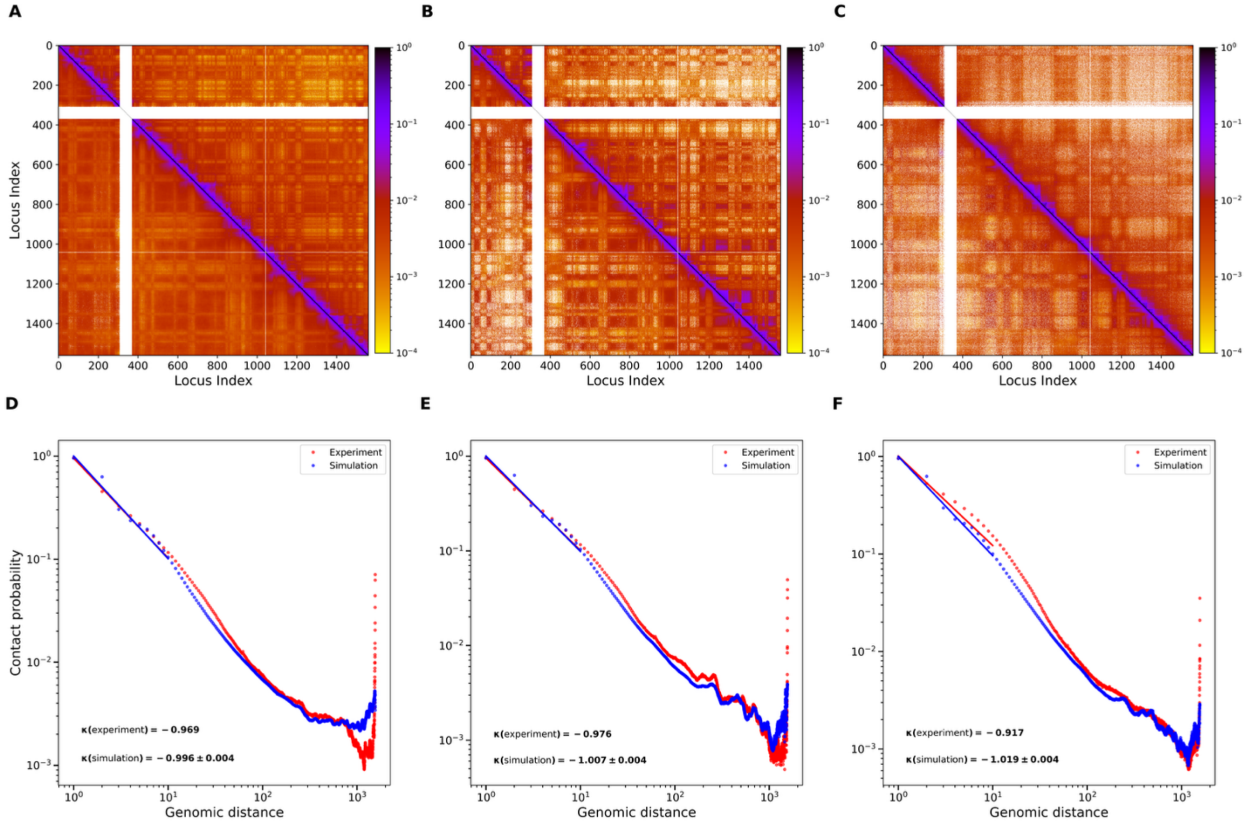


**Fig S17. Validation of chromosome 18 chromatin architecture simulated by the polymer model across three human cell types. (A–C)** Experimental Hi-C contact maps (upper triangles) are compared with polymer simulation contact maps (lower triangles) for chromosome 18 in **GM12878** (A), **IMR90** (B), and **H1-hESC** (C). Contact frequencies are displayed on a logarithmic color scale. The close visual agreement between experimental and simulated maps demonstrates that the model accurately reproduces chromosome-scale interaction patterns across diverse chromatin contexts. **(D–F)** Contact probability as a function of genomic distance, $P(s)$, derived from experimental Hi-C data (red) and polymer simulations (blue) for **GM12878** (D), **IMR90** (E), and **H1-hESC** (F). The fitted power-law scaling exponent ($\kappa$) is shown for both experiment and simulation in each panel, demonstrating that the simulations faithfully recapitulate the distance-dependent decay of chromatin contacts observed experimentally.

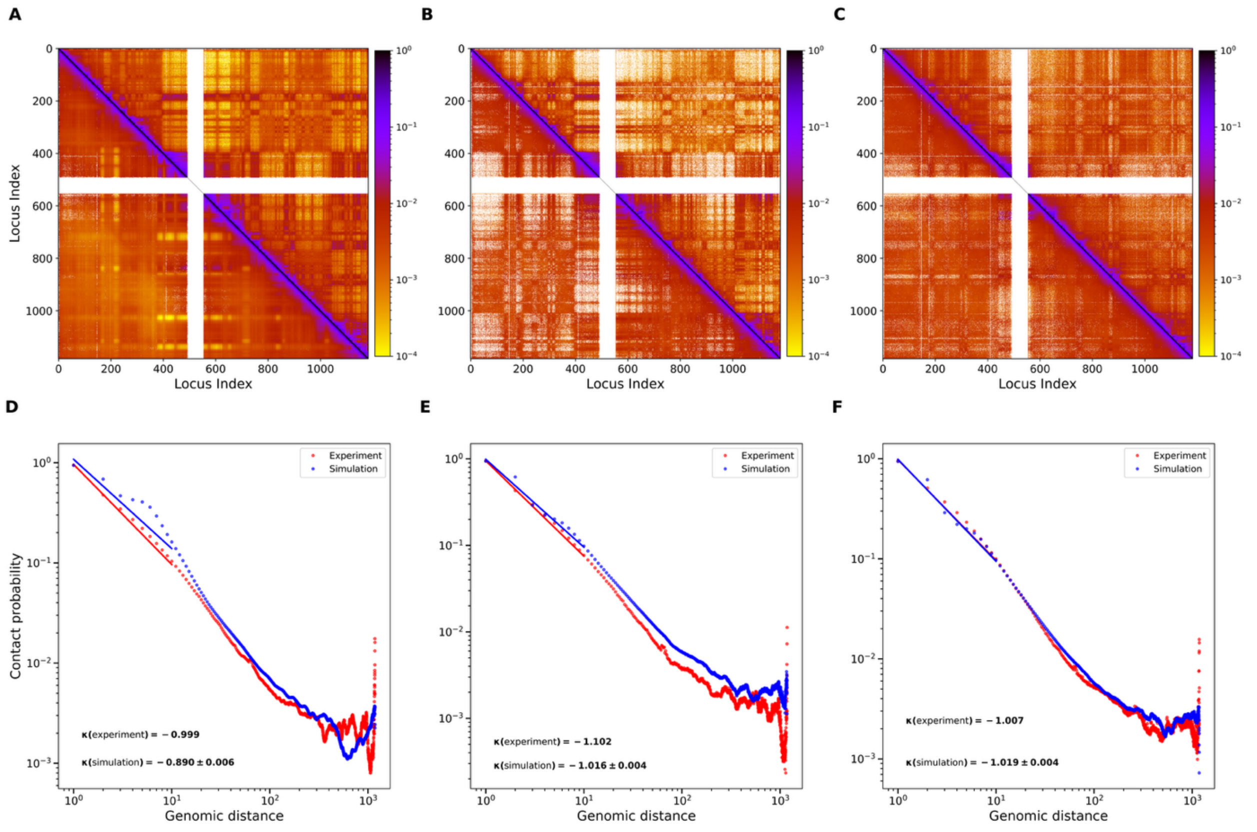


**Fig S18. Validation of chromosome 19 chromatin architecture simulated by the polymer model across three human cell types. (A–C)** Experimental Hi-C contact maps (upper triangles) are compared with polymer simulation contact maps (lower triangles) for chromosome 19 in **GM12878** (A), **IMR90** (B), and **H1-hESC** (C). Contact frequencies are displayed on a logarithmic color scale. The close visual agreement between experimental and simulated maps demonstrates that the model accurately reproduces chromosome-scale interaction patterns across diverse chromatin contexts. **(D–F)** Contact probability as a function of genomic distance, $P(s)$, derived from experimental Hi-C data (red) and polymer simulations (blue) for **GM12878** (D), **IMR90** (E), and **H1-hESC** (F). The fitted power-law scaling exponent ($\kappa$) is shown for both experiment and simulation in each panel, demonstrating that the simulations faithfully recapitulate the distance-dependent decay of chromatin contacts observed experimentally.

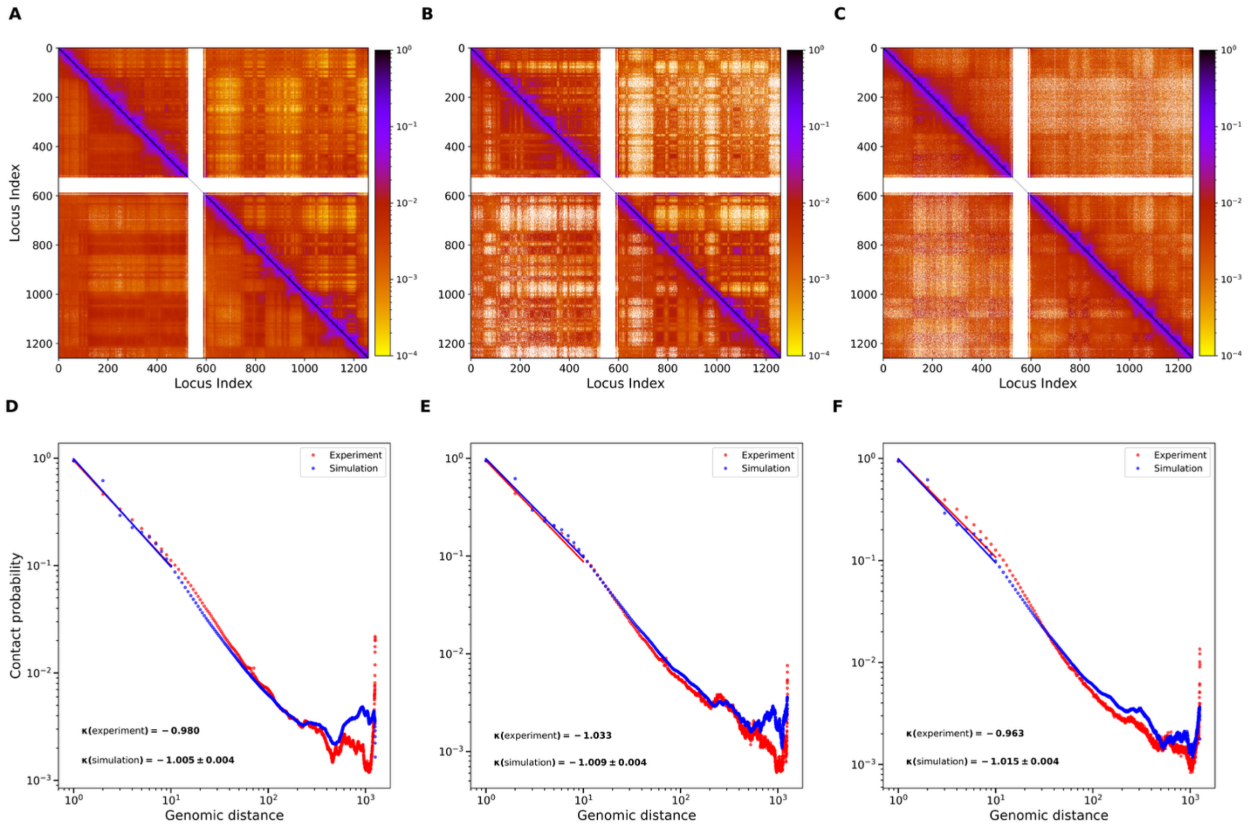


**Fig S19. Validation of chromosome 20 chromatin architecture simulated by the polymer model across three human cell types. (A–C)** Experimental Hi-C contact maps (upper triangles) are compared with polymer simulation contact maps (lower triangles) for chromosome 20 in **GM12878** (A), **IMR90** (B), and **H1-hESC** (C). Contact frequencies are displayed on a logarithmic color scale. The close visual agreement between experimental and simulated maps demonstrates that the model accurately reproduces chromosome-scale interaction patterns across diverse chromatin contexts. **(D–F)** Contact probability as a function of genomic distance, $P(s)$, derived from experimental Hi-C data (red) and polymer simulations (blue) for **GM12878** (D), **IMR90** (E), and **H1-hESC** (F). The fitted power-law scaling exponent ($\kappa$) is shown for both experiment and simulation in each panel, demonstrating that the simulations faithfully recapitulate the distance-dependent decay of chromatin contacts observed experimentally.

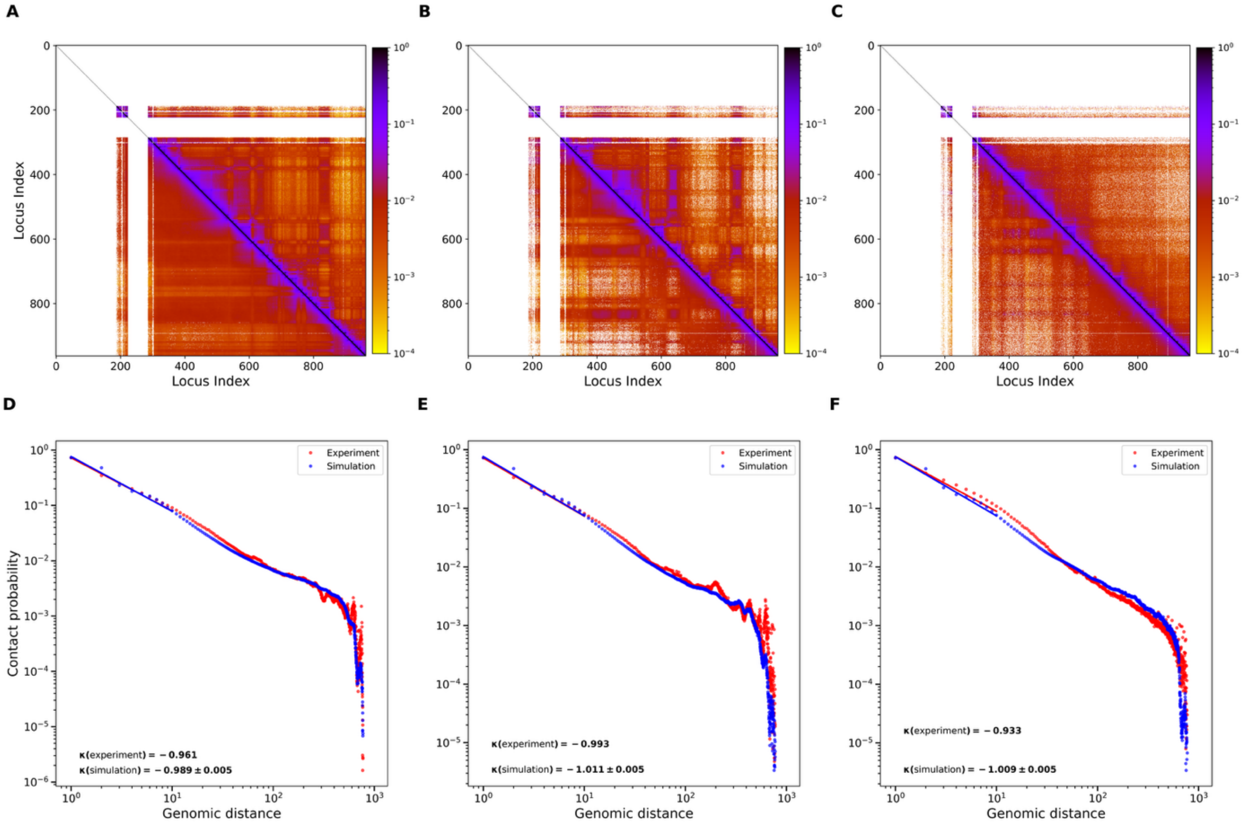


**Fig S20. Validation of chromosome 21 chromatin architecture simulated by the polymer model across three human cell types. (A–C)** Experimental Hi-C contact maps (upper triangles) are compared with polymer simulation contact maps (lower triangles) for chromosome 21 in **GM12878** (A), **IMR90** (B), and **H1-hESC** (C). Contact frequencies are displayed on a logarithmic color scale. The close visual agreement between experimental and simulated maps demonstrates that the model accurately reproduces chromosome-scale interaction patterns across diverse chromatin contexts. **(D–F)** Contact probability as a function of genomic distance, $P(s)$, derived from experimental Hi-C data (red) and polymer simulations (blue) for **GM12878** (D), **IMR90** (E), and **H1-hESC** (F). The fitted power-law scaling exponent ($\kappa$) is shown for both experiment and simulation in each panel, demonstrating that the simulations faithfully recapitulate the distance-dependent decay of chromatin contacts observed experimentally.

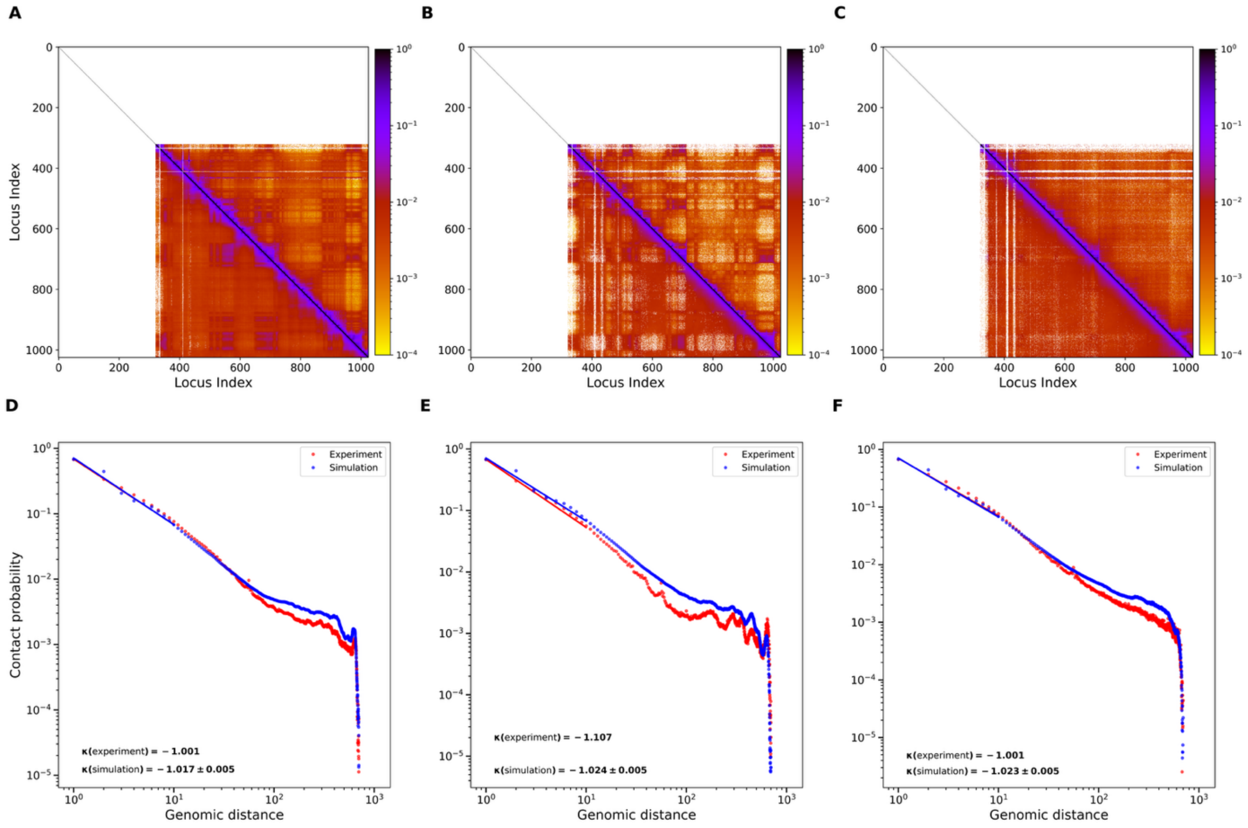


**Fig S21**. **Validation of chromosome 22 chromatin architecture simulated by the polymer model across three human cell types. (A–C)** Experimental Hi-C contact maps (upper triangles) are compared with polymer simulation contact maps (lower triangles) for chromosome 22 in **GM12878** (A), **IMR90** (B), and **H1-hESC** (C). Contact frequencies are displayed on a logarithmic color scale. The close visual agreement between experimental and simulated maps demonstrates that the model accurately reproduces chromosome-scale interaction patterns across diverse chromatin contexts. **(D–F)** Contact probability as a function of genomic distance, $P(s)$, derived from experimental Hi-C data (red) and polymer simulations (blue) for **GM12878** (D), **IMR90** (E), and **H1-hESC** (F). The fitted power-law scaling exponent ($\kappa$) is shown for both experiment and simulation in each panel, demonstrating that the simulations faithfully recapitulate the distance-dependent decay of chromatin contacts observed experimentally.

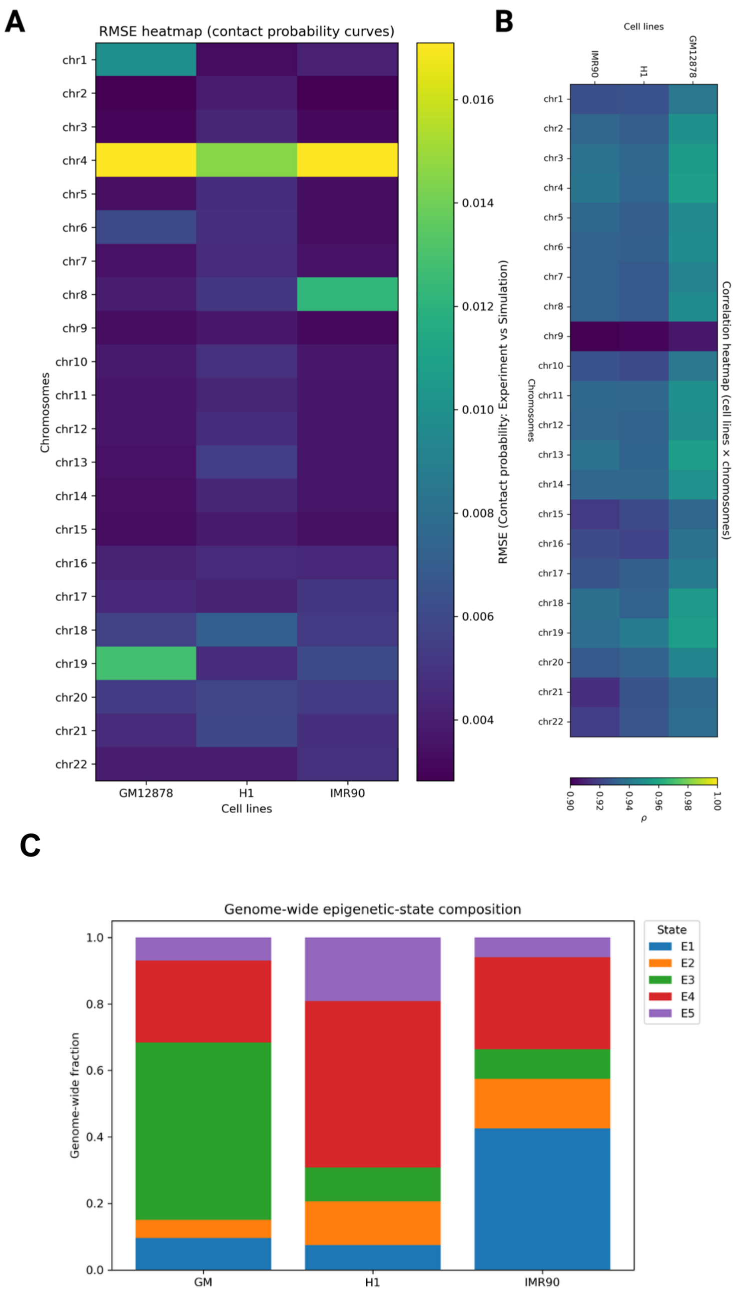


**Fig S22. The optimized model reproduces experimental Hi-C organization, and cell-type-specific folding is epigenetically encoded. (A)** RMSE between experimental and simulated contact-probability curves per chromosome and cell line (color scale ≈0.004–0.017), uniformly low with modest peaks at chromosome 4 (all lines), chromosome 19 (GM12878), and chromosome 8 (IMR90). **(B)** Pearson correlation between simulated and experimental contact matrices for chromosome 1–chromosome 22 across GM12878, H1-hESC, and IMR90; correlations are uniformly high.; (C) Genome-wide epigenetic-state composition (E1–E5) across GM12878, H1-hESC, and IMR90, showing distinct chromatin-state distributions, with E3 predominating in GM12878, E4 in H1-hESC, and E1 in IMR90.

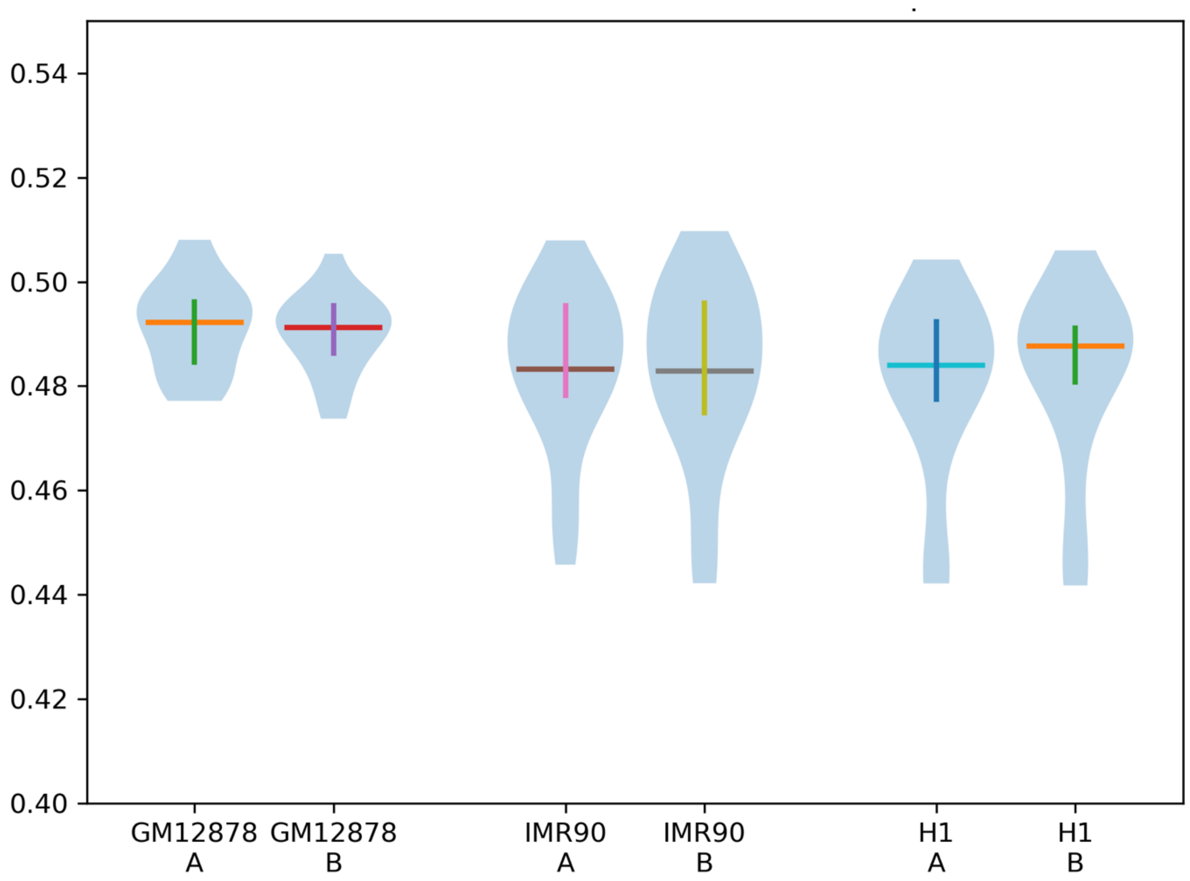


**Fig S23.** Distribution of the MSD exponent α (MSD ~ $\tau^{\alpha}$) across cell lines and A/B compartments; violins show the distribution and crosses denote mean ± spread. All distributions are subdiffusive (α ≈ 0.44–0.50) with overlapping error bars.

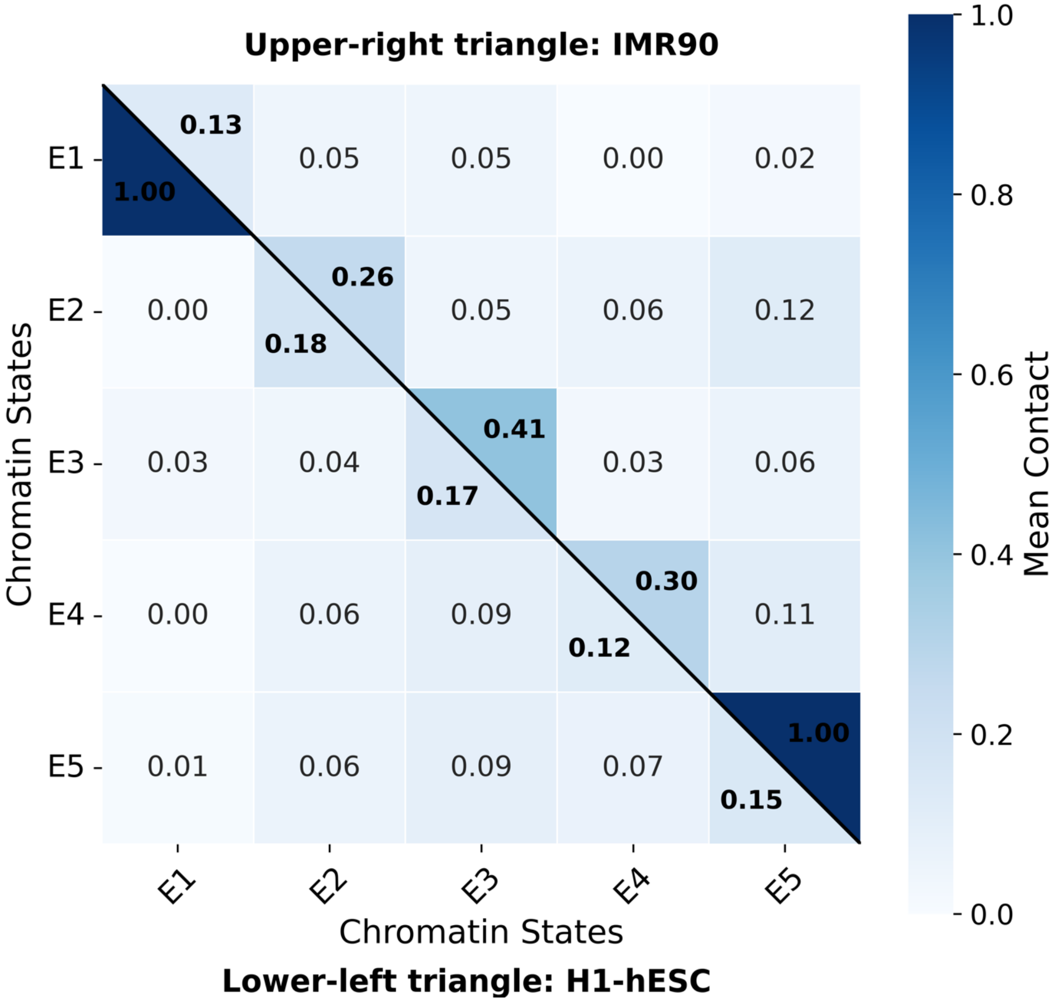


**Fig S24.** State–state interaction heatmaps (normalized mean contact, 0–1) for epigenetic states E1–E5; diagonal = self-interactions, off-diagonal = cross-state mixing. Each panel overlays two cell lines (IMR90 (upper) vs. H1-hESC (lower triangle)).

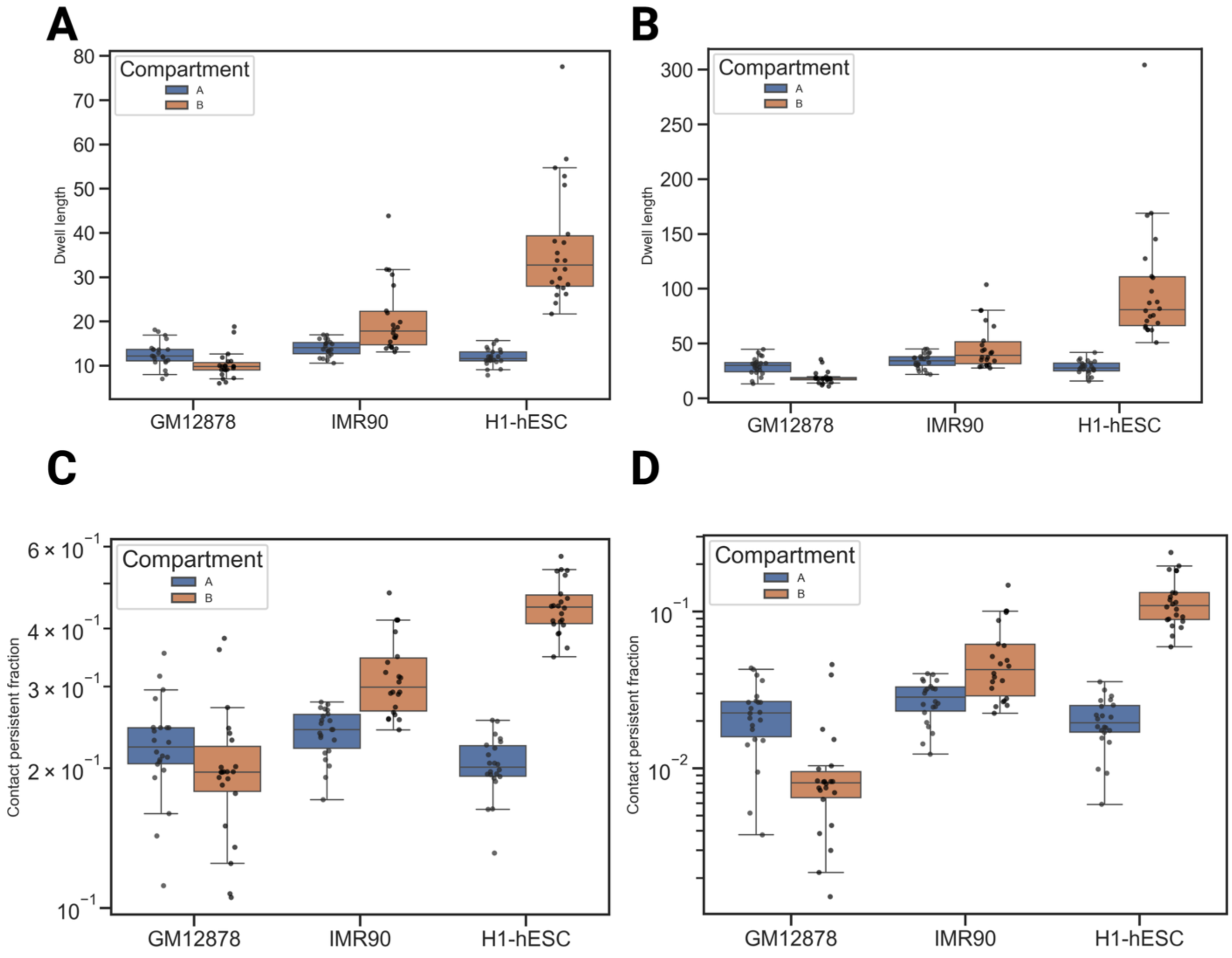


**Fig S25.** Compartment-resolved dwell length metrics Q95 (A) and Q99 (B) for GM12878, IMR90, and H1-hESC, shown separately for the euchromatin and heterochromatin compartments. long-lived-contact survival fractions S≥5 (C) and S≥20 (D) for GM12878, IMR90, and H1-hESC are displayed separately for two distinct compartments.

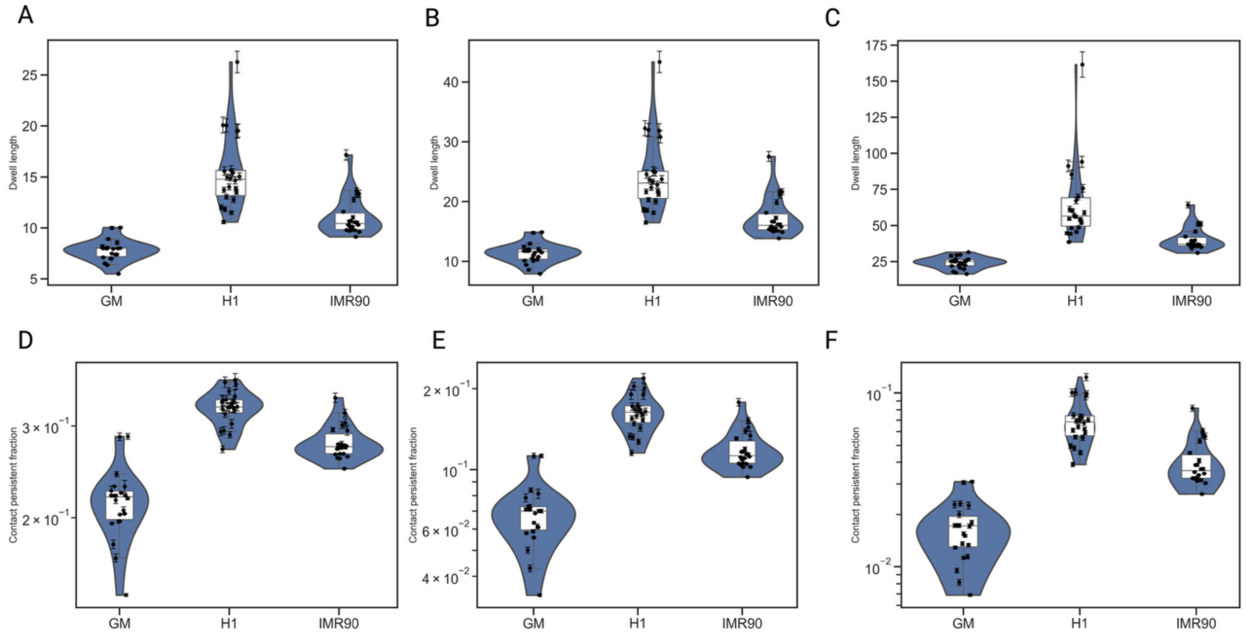


**Fig S26.** Whole-genome dwell length metrics reconstructed by recombining euchromatin and heterochromatin statistics as contact-weighted averages Q90 (**A**), Q95 (**B**), and Q99 (**C**); for every metric, the recombined mean follows GM12878 < IMR90 < H1-hESC, with the largest separation at Q99. (D-F) Whole-genome persistence metrics reconstructed by recombining euchromatin and heterochromatin statistics as contact-weighted averages S≥5 (**D**), S≥10 (**E**), and S≥20 (**F**).

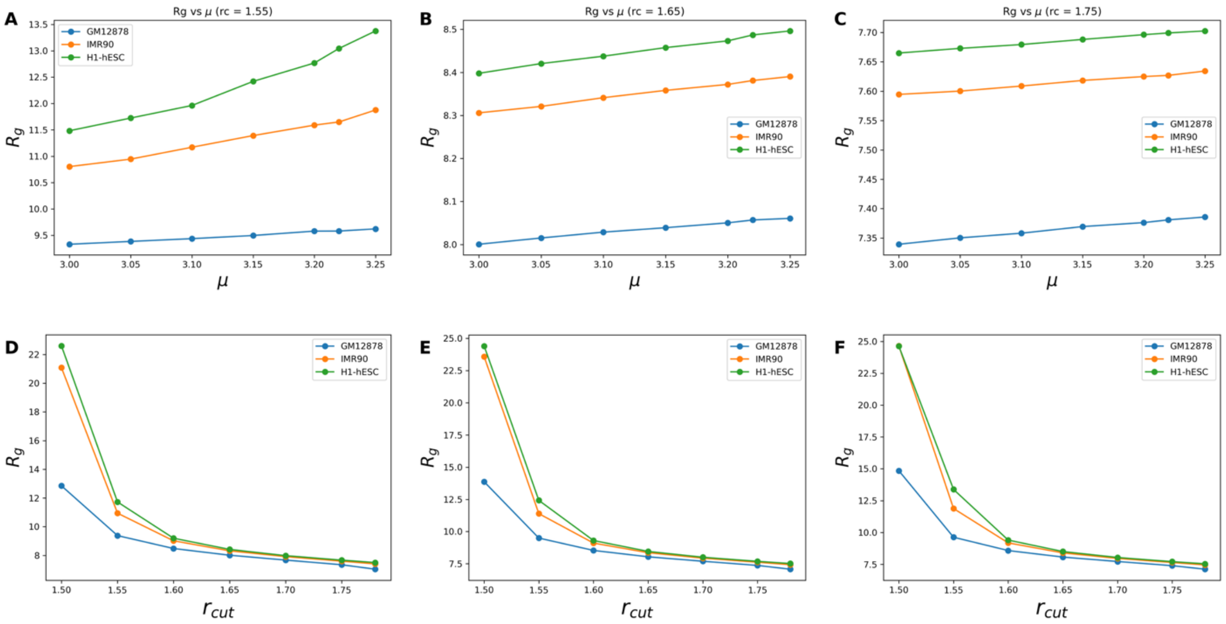


**Fig S27. Sensitivity of the radius of gyration ($R_g$) to the model parameters $\mu$ and $r_{cut}$ across three human cell types.** Each panel reports the ensemble-averaged radius of gyration $R_g$ of the reconstructed 3D chromatin conformations for GM12878 (blue), IMR90 (orange), and H1-hESC (green). **(A–C)** Dependence of $R_g$ on the contact-to-distance conversion exponent $\mu$ over the range $\mu$ from 3.00 to 3.25, shown at three fixed interaction cutoffs $r_{cut} = 1.55$(A), 1.65(B), and 1.75(C). Across all cutoffs $R_g$ increases monotonically and approximately linearly with $\mu$, with the cell-line ordering ($R_g^{\text{H1-hESC}} > R_g^{\text{IMR90}} > R_g^{\text{GM12878}}$) preserved throughout; the absolute spread between cell lines and the sensitivity to $\mu$ both diminish sharply as $r_{cut}$ increases. **(D–F)** Dependence of $R_g$ on the cutoff $r_{cut}$ from 1.50 to 1.78 at the correspondingly fixed values of $\mu$. $R_g$ decays steeply and nonlinearly with increasing $r_{cut}$, converging toward a common plateau ($R_g \approx 7$–8 in the model unit) for $r_{cut} \gtrsim 1.65$, where the three cell lines become nearly indistinguishable. Markers denote computed values at each parameter setting and solid lines are guides to the eye. Together, the panels indicate that $r_{cut}$ is the dominant determinant of overall chain compaction, whereas $\mu$ provides finer-grained tuning, and that cell-line-specific differences in $R_g$ are most pronounced in the small-cutoff, expanded-conformation regime.

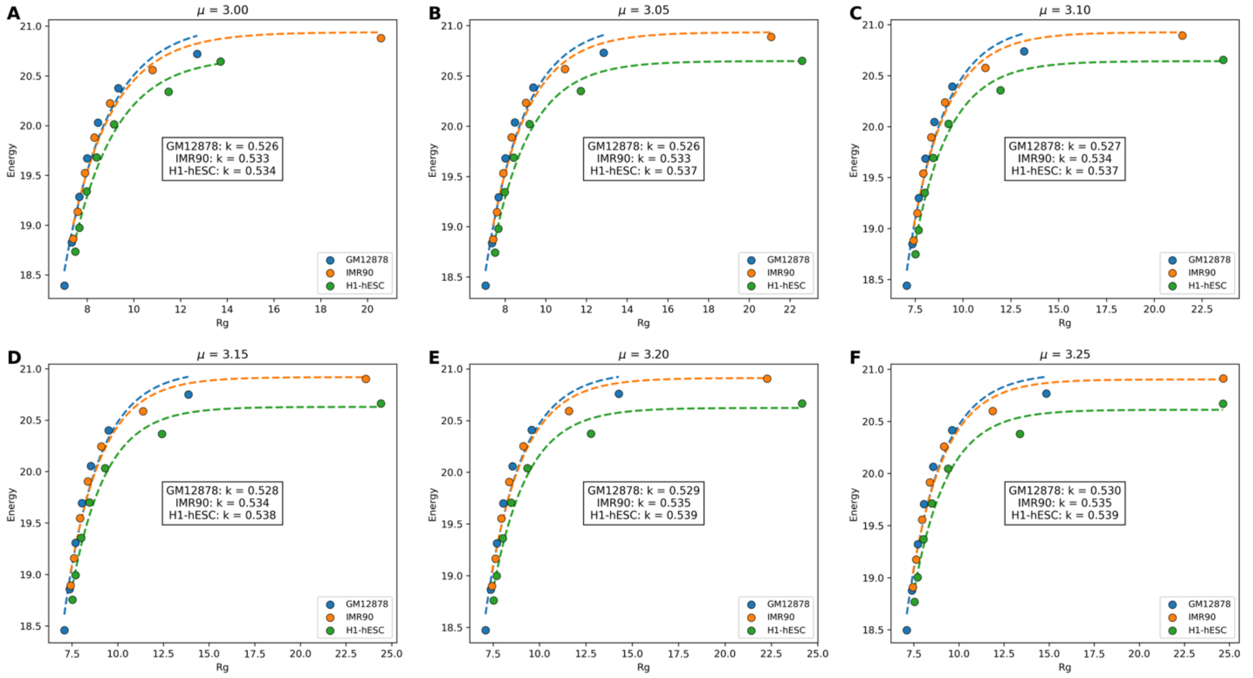


**Fig S28. Relationship between conformational energy and radius of gyration ($R_g$) across the interaction-parameter sweep for three human cell lines.** Each panel plots the model energy against the radius of gyration $R_g$ for reconstructed 3D chromatin conformations of GM12878 (blue), IMR90 (orange), and H1-hESC (green), with points generated by sweeping the cutoff-distance parameter $r_{cut}$. Panels **(A–F)** correspond to increasing values of the interaction parameter $\mu = 3.00$, 3.05, 3.10, 3.15, 3.20, and 3.25, respectively. In every panel the energy rises steeply at small $R_g$ and saturates toward a plateau as $R_g$ increases, indicating that the more expanded conformations obtained at small $r_{cut}$ approach an energetic ceiling. Dashed curves are per-cell-line fits to this saturating relationship, and the inset reports the corresponding fitted scaling exponent $k$ for each cell line. The exponent is remarkably stable across the full $\mu$ range, varying by less than 0.005 per cell line, and preserves a consistent ordering ($k_{\text{H1-hESC}} > k_{\text{IMR90}} > k_{\text{GM12878}}$, spanning $\approx$ 0.526–0.539). This near-invariance demonstrates that the energy–$R_g$ scaling is governed by an intrinsic, cell-line-specific property that is robust to the choice of interaction parameter $\mu$, with H1-hESC exhibiting the strongest energy response to compaction and GM12878 the weakest.

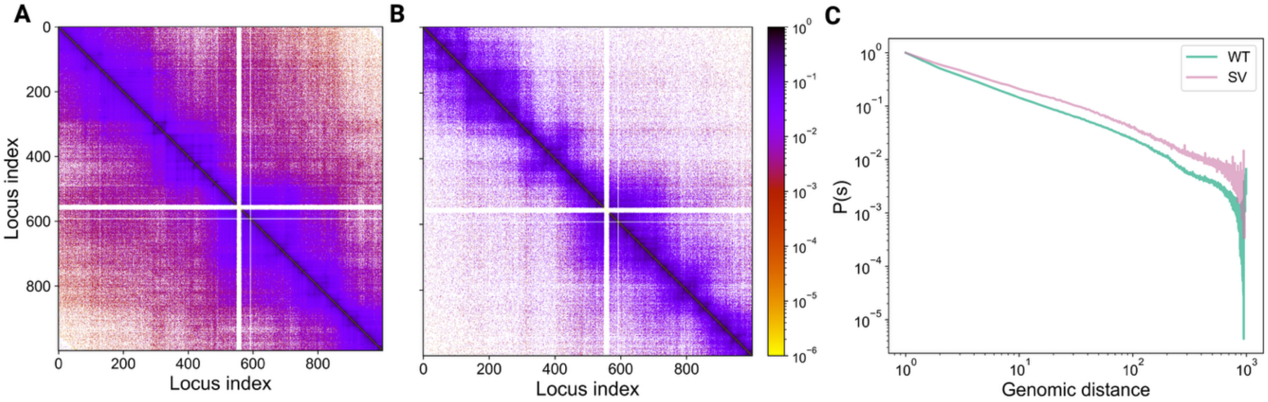


**Fig S29. Structural-variant (SV) rearrangement reshapes the chromatin contact landscape and its distance-scaling behavior for Chromosome 8. (A)** Contact frequency matrix for the WT-genome (GM19036) and **(B)** the corresponding SV matrix (HG00864), both plotted as heatmaps of pairwise contact probability as a function of locus index. Both maps display the expected strong diagonal, block-like topologically associating domain (TAD) structure, and higher-order compartmentalization. **(C)** Genome-wide contact probability $P(s)$ as a function of genomic separation $s$ (both axes logarithmic) for WT (green) and SV (pink). The two curves overlap at short range but diverge at intermediate-to-long separations, where the SV genome maintains systematically higher contact probability, indicating that the rearrangement promotes additional long-range contacts and a shallower decay of $P(s)$ characteristic of a more compact or reconfigured large-scale organization.

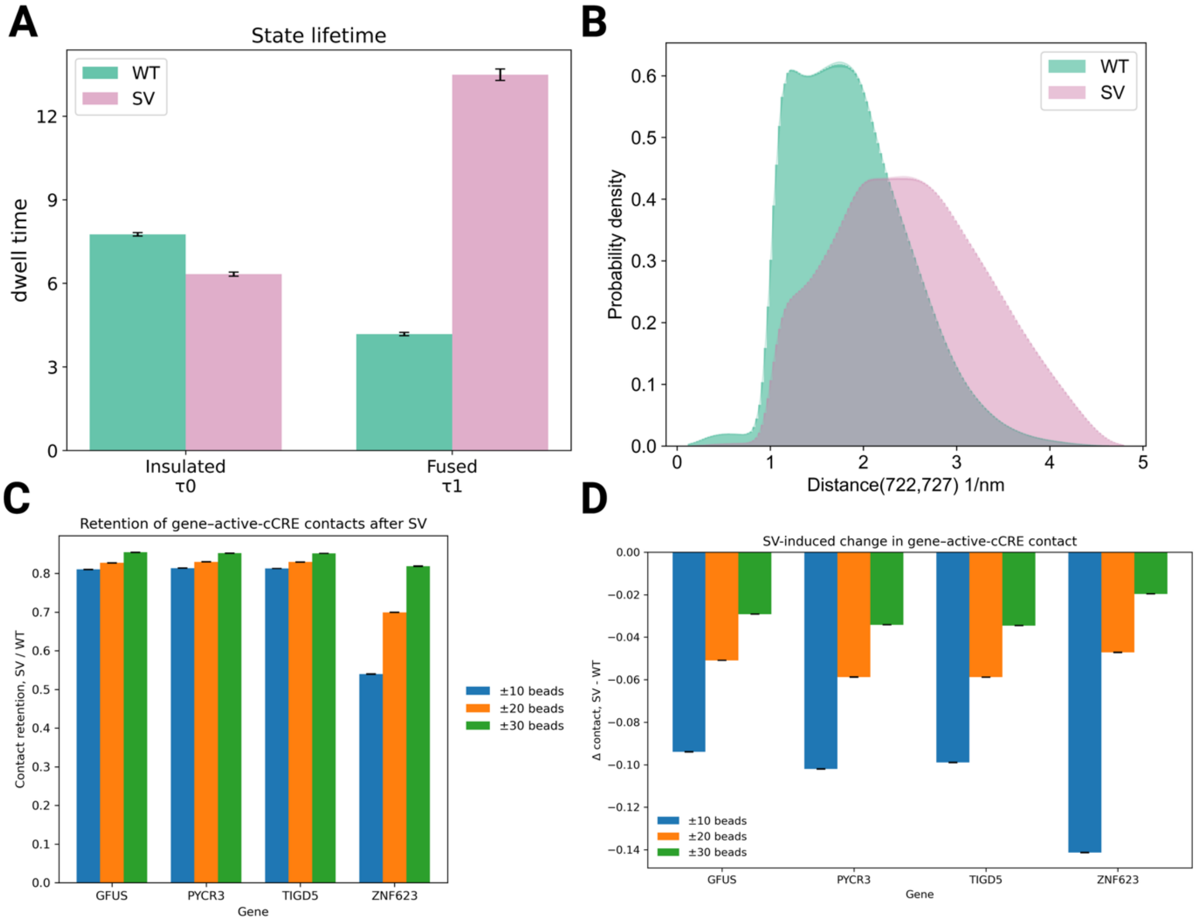


**Fig S30. Structural variation alters chromatin-state dynamics and disrupts local gene–regulatory contacts. (A)** Mean dwell time of chromatin conformational states before (wild type; WT/NOSV) and after structural variation (SV). Following the deletion, the insulated state is short-lived while the fused state is markedly stabilized, indicating a shift toward persistent domain fusion. Error bars denote the standard error across simulation replicates. **(B)** Probability density of the inter-locus distance between beads 722 and 727, which become juxtaposed after the deletion. SV shifts the distribution toward larger equilibrium separations and broadens it, reflecting increased conformational heterogeneity and reorganized local chromatin architecture. **(C)** Retention of gene–active cCRE (candidate cis-regulatory element) contacts after SV, evaluated within ±10-, ±20-, and ±30-bead windows around each gene. Retention increases with window size for most genes; ZNF623 is a notable exception, showing sharply reduced retention within the ±10-bead window and thus heightened sensitivity to local structural perturbation. **(D)** Mean change in gene–active cCRE contact frequency (SV − WT) across the same windows. All genes lose regulatory contacts after the deletion, with the largest losses in the immediate neighborhood and progressively smaller effects at broader genomic scales. ZNF623 shows the greatest local loss, consistent with pronounced SV-induced disruption of its regulatory landscape.

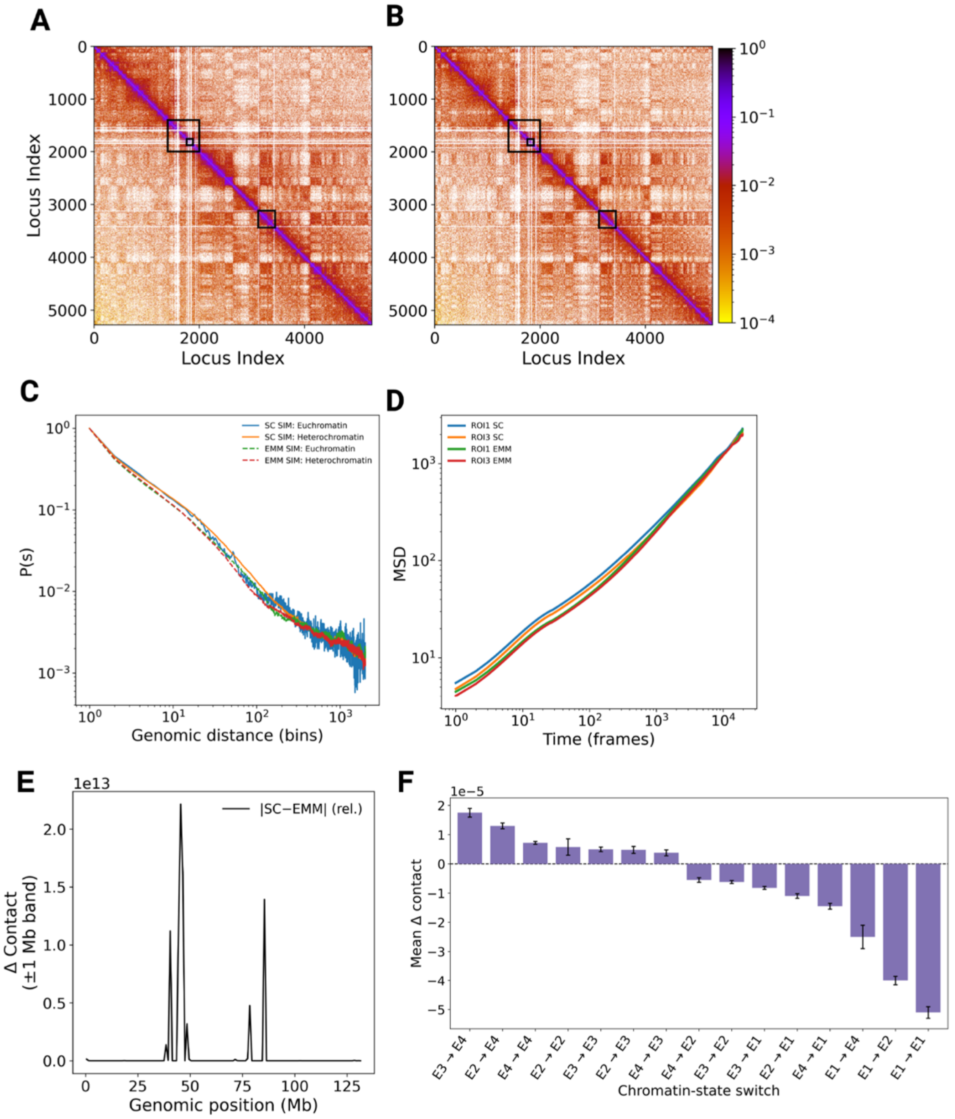


**Fig S31. The maximum-entropy model faithfully reproduces chromatin organization and dynamics.** **(A)** Reference Hi-C contact-probability matrix from the sigmoid colon (SC) tissue; black boxes mark representative regions used for local structural comparison. **(B)** Contact-probability matrix for esophageal muscularis mucosa (EMM) tissue, showing close agreement with the reference across both local and long-range interactions. **(C)** Contact probability $P(s)$versus genomic separation $s$for euchromatic and heterochromatic regions. The EMM reproduces the reference scaling across genomic distances, indicating that polymer-folding properties are preserved in both chromatin environments. **(D)** Mean-squared displacement (MSD) of representative loci within the highlighted regions (ROI1 and ROI3). The near-overlapping SC and EMM trajectories show that the inferred model preserves chromatin dynamics as well as structure. **(E)** Sliding-window analysis of mean relative absolute contact differences between SC and EMM across chromosome 10 (±1 Mb), highlighting regions of tissue-specific contact divergence. **(F)** Mean contact-frequency changes (SC − EMM) across chromatin-state transition categories. Positive and negative values indicate stronger contacts in SC and EMM, respectively. Error bars represent standard deviations across 40 simulation runs.

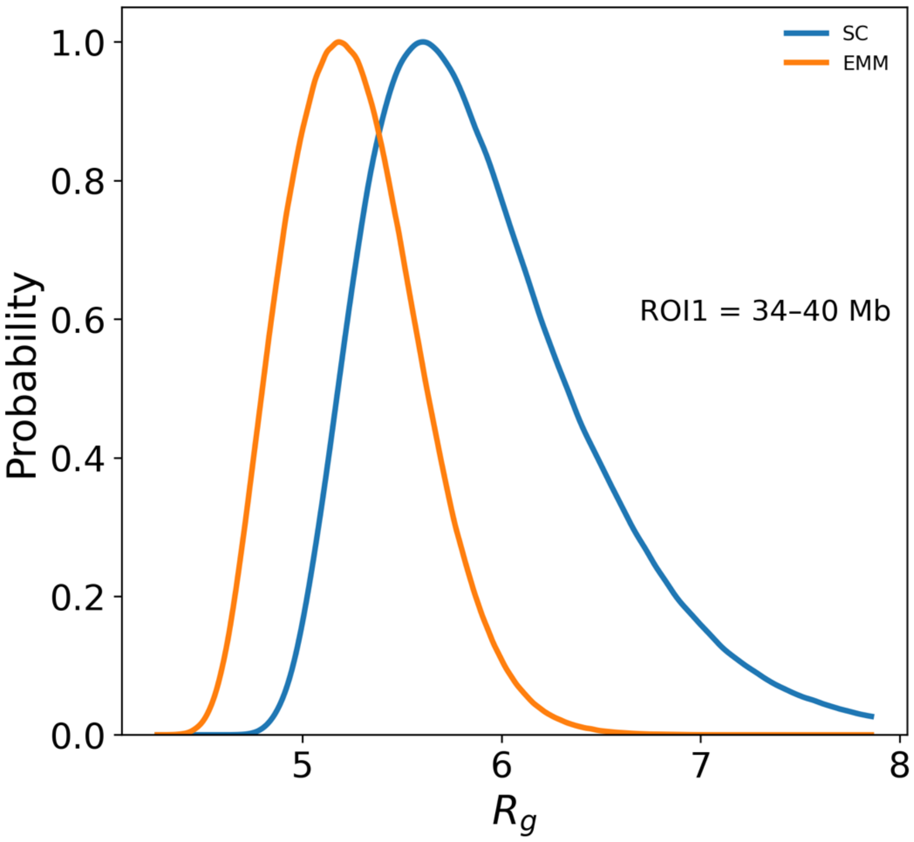


**Fig S32.** The radius of gyration distribution comparison for conformational ensembles generated by modeling sigmoid colon (SC) and esophageal muscularis mucosa (EMM) tissues.
